# Combining Multi-Dimensional Molecular Fingerprints to Predict hERG Cardiotoxicity of Compounds

**DOI:** 10.1101/2021.06.06.447291

**Authors:** Weizhe Ding, Li Zhang, Yang Nan, Juanshu Wu, Xiangxin Xin, Chenyang Han, Siyuan Li, Hongsheng Liu

## Abstract

At present, drug toxicity has become a critical problem with heavy medical and economic burdens. acLQTS (acquired Long QT Syndrome) is acquired cardiac ion channel disease caused by drugs blocking the hERG channel. Therefore, it is necessary to avoid cardiotoxicity in the drug design and computer models have been widely used to fix this plight. In this study, we present a molecular fingerprint based on the molecular dynamic simulation and uses it combined with other molecular fingerprints (multi-dimensional molecular fingerprints) to predict hERG cardiotoxicity of compounds. 203 compounds with hERG inhibitory activity (pIC50) were retrieved from a previous study and predicting models were established using four machine learning algorithms based on the single and multi-dimensional molecular fingerprints. Results showed that MDFP has the potential to be an alternative to traditional molecular fingerprints and the combination of MDFP and traditional molecular fingerprints can achieve higher prediction accuracy. Meanwhile, the accuracy of the best model, which was generated by consensus of four algorithms with multi-dimensional molecular fingerprints, was 0.694 (RMSE) in the test dataset. Besides, the number of hydrogen bonds from MDFP has been determined as a critical factor in the predicting models, followed by rgyr and sasa. Our findings provide a new sight of MDFP and multi-dimensional molecular fingerprints in building models of hERG cardiotoxicity prediction.

## 1. Introduction

Drug-induced toxicity has become a critical reason for the failure of drug discovery and development in recent years (Wallace, 2015). A previous study showed that there were more than half of drugs failed (54%) in clinical development among 640 novel therapeutics, while 17% of them failed because of drug-induced toxicity (Hwang et al., 2016). Besides, it has also been reported that the mean costs required to bring a new drug to market increased from $374.1 million to $1335.9 million after counting for costs of failed trials (Wouters et al., 2016). Thus, it has become an urgent task to find ways to identify the toxicity of compounds on a large scale in drug development.

Acquired Long QT syndrome (acLQTS), one of the most important diseases caused by drug-induced toxicity, is a potentially life-threatening cardiac arrhythmia disease that increases the risk for syncope, sudden cardiac death (SCD), and seizures (Tester & Ackerman, 2014). The hERG protein is a tetrameric potassium ion channel and mainly relates to cardiotoxicity and acLQTS (Liu et al., 2020). It has been reported that the potassium ion channel (hERG channel) may be blocked caused by antiarrhythmic drug binding, which leads to prolonged repolarization time and acLQTS (Witchel, 2007). At present, multiple drug candidates have failed due to the cardiotoxicity of hERG, such as cisapride, terfenadine, sertindole, pimozide, and astemizole, which have become a significant limiting factor in drug discovery and development (Bergström & Lindmark, 2019; Villoutreix & Taboureau, 2019).

Computer-aided drug design (CADD) has been thought of as an alternate choice to reduce the amount of time and money in the development of drug design, especially in predicting drug toxicity (Maia et al., 2020). Molecular fingerprints are a way of CADD and are used to encoding the structure of molecules (O’Boyle et al., 2011). It has been deployed as descriptors for predicting biological activities and compound properties (Muegge & Mukherjee, 2014). Frequently used molecular fingerprints are structure-based and property-based (Kelley, 2018; Rogers & Hahn, 2010; Riniker & Landrum, 2013; Riniker, 2017). A previous study of hERG cardiotoxicity prediction showed that the accuracy of the best model developed by molecular descriptors reached 0.54 (R^2^), while RMSE was 0.63 (Johnson et al., 2007). Another study of the hERG channel also showed that the accuracy of the regression model by descriptors was 0.60 (Q^2^) and 0.55 (RMSE) for pIC50 (Radchenko et al., 2017). These results showed the practicalities and effectiveness based on commonly used molecular fingerprints. However, there are still no fingerprints that considered the time factor applied on the cardiotoxicity prediction of hERG.

Molecular dynamics fingerprints (MDFP) are the fingerprints based on calculating the trajectory of molecular dynamic simulation and have rapidly become a hotspot. After adding the dimension of time, MDFP can be seen as a choice of the traditional molecular fingerprint. The study of p-glycoprotein substrates prediction showed that gradient tree boosting (GTB) methods in combination with MDFP was the only model which achieved a good accuracy on the in-house dataset (Esposito et al., 2020). Meanwhile, the research of free-energy prediction showed good performance with a heterogeneous fusion model by MDFP (Riniker, 2017). Besides, studies of self-solvation free energies and application of MDFP in SAMPL6 octanol–water log P blind challenge also revealed a high prediction rate (Gebhardt et al., 2020; Wang & Riniker, 2019). As a consequence, MDFP can be an alternative choice of traditional molecular fingerprints and has great application potential on the cardiotoxicity prediction of hERG.

Multi-dimensional molecular fingerprints are indicated as multiple molecular fingerprints combining together in order to predict more accurately. Previous studies showed that multi-dimensional molecular fingerprints were better than the single molecular fingerprint in drug development (Kyaw et al., 2020). Thus, in this study, we studied MDFP and multi-dimensional molecular fingerprints (MDFP with other molecular fingerprints) in predicting hERG cardiotoxicity of compounds. The extensive open dataset of hERG compounds with IC50 values has been collected from previous studies. Then, molecular dynamic simulation was conducted to generate MDFP and traditional molecular fingerprints have also been generated by Baseline2D, ECFP4, and PropertyFP. Finally, the regression models were built by machine learning with four algorithms. Our study provides new sights on the combination of multi-dimensional molecular fingerprints and the research of predicting the hERG cardiotoxicity of compounds.

## 2. Methods

### 2.1. Toxicity Datasets

A high-quality hERG inhibitor dataset has been collected from the previous study (Munawar et al., 2019). The IC50 value is the biochemical half-maximal inhibitory concentration and has been used to represent the inhibiting abilities of compounds on hERG in this dataset (Kalliokoski et al., 2013). The data of toxicity have been eliminated if the name and IC50 values were repeated. The repeated molecules have also been averaged if the difference IC50 values were less than one order of magnitude (Feng et al., 2021). Finally, 203 compounds have been collected with specific IC50 values of the hERG. The distribution of training and testing sets followed by 80% and 20%, respectively. The training sets were used for 5-fold cross-validation and the testing sets were used to check the prediction performance of the established model for new compounds. Besides, pIC50 is the negative log unit of the IC50 values and has been used to represent inhibiting abilities better than IC50 (Cortés-Ciriano et al., 2020). Therefore, IC50 of compounds was converted to pIC50.

### 2.2. MD Simulations

Molecular dynamics (MD) simulation was performed by GROMACS (2020.4). For compounds in the dataset, mol2 files were obtained from Zinc15 (http://zinc15.docking.org/) by using SMILES files. The topology of compounds was generated with AMBER14SB force field by ACPYPE (https://www.bio2byte.be/) (Sousa da Silva et al., 2012). Afterward, the compounds were placed in a dodecahedron box with a size of 1.0 nm centrally and solvated with the TIP3P water model. Then, the descent energy minimization with 100ps was applied to the system. An additional equilibration of 1ns under NVT and NPT conditions was carried out, while the constant temperature was 300 K and the constant pressure was 1 bar, respectively (Sun et al., 2020). Finally, the system was performed with running 5 ns MD simulation and coordinates were written every 10ps, energies every 1ps.

### 2.3. 2D Molecular Fingerprints

Three types of molecular fingerprints have been used in this study. Baseline2D was obtained using RDKit and its elements mainly consisted of 10 counts: number of heavy atoms, number of rotatable bonds, number of N, O, F, P, S, Cl, Br, and I atoms (Riniker, 2017; Wang & Riniker, 2019). The PropertyFP fingerprint was also obtained using the Descriptastorus package from RDKit (Kelley, 2018). It contained nearly 200 atoms features and properties. Besides, ECFP4 was generated using the RDKit implementation of the Morgan algorithm with a vector length of 2048 and a radius of 2 ( Rogers & Hahn, 2010).

### 2.4. MD Fingerprints

The MD trajectories were analyzed by the GROMACS toolkit (Ogunwa, 2019). Following features has been generated: radius of gyration (rgyr), solvent-accessible surface area (sasa), root mean squared error (rmsd), total energy (tenergy), hydrogen bonds (hbond), kinetic energy (kinetic), Lennard-Jones short-range energies (LJ-SR) and Lennard-Jones 1-4 energies (LJ-14). The average (avr), median (mid), and standard deviation (std) of features were calculated using the R version 3.6.1 (Team, 2013). Fig. 1 showed the MDFP with all properties.

**Fig.1.**
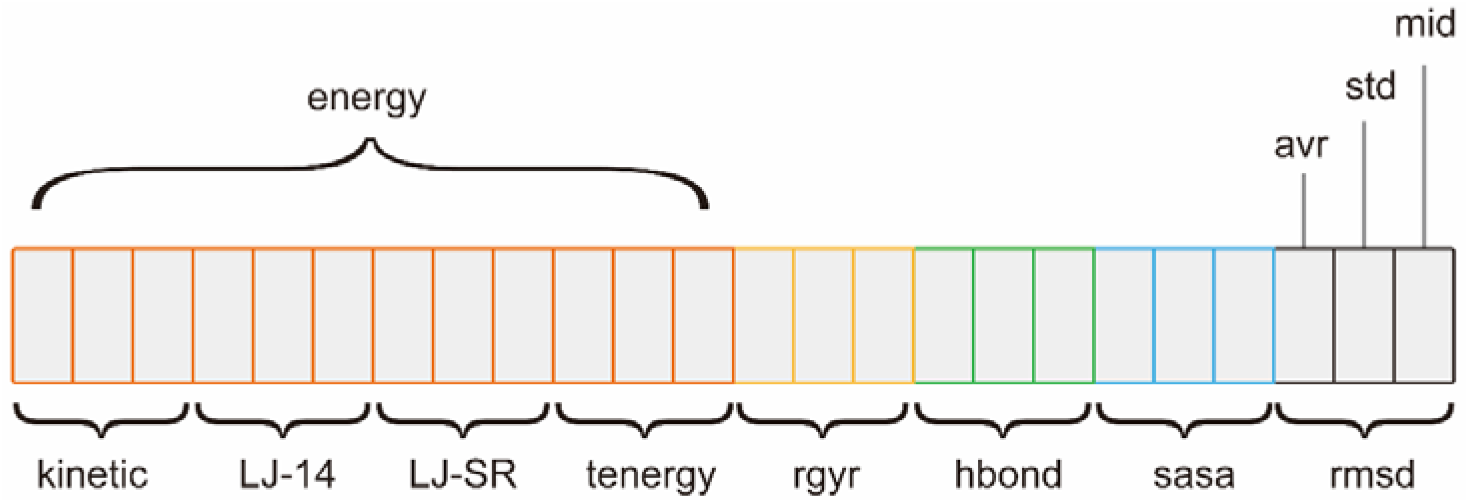
Schematic representation of the MDFP variant with all properties: kinetic, LJ-14, LJ-SR, tenergy, rgyr, hbond, sasa, rmsd. Each property is represented by the avr (average), std (standard deviation), and mid (median).

### 2.5. Feature Selection

Feature selection is critically important for predictive models, especially in machine learning (Johnson et al., 2018). It provides an effective way to reduce the dimensionality of data sets, identify informative features, and remove irrelevant features, improving the learning accuracy of machine learning models (Holder et al., 2017). In this study, zero variation and near-zero variation features were deleted firstly using the nearZeroVar function in the R package caret (version 6.0–84) (Kuhn, 2008). Then, recursive feature elimination (RFE) was performed to select the optimal feature subset using the rfe function in the caret package in a 10 times 5-fold cross-validation setting (Darst et al., 2018). In the RFE process, all features are first ranked according to the feature importance values obtained by the random forest (RF) algorithm, and then RF models are trained iteratively on the features that are gradually reduced according to the ranking to evaluate the performance of the feature subsets (Tang et al., 2020).

### 2.6. Model Construction

In this study, RF, SVM, gradient boosting machine (GBM), and partial least square regression (PLS) was used for machine learning model construction. All models were executed beyond R (version 3.6.1) with using the randomForest (version 4.6–12) (Liaw & Wiener, 2002), the kernlab (version 0.9-25) (Karatzoglou et al., 2004), the gbm (version 2.1.5) (Brandon et al., 2019), and the pls (version 2.7-1) packages (Bjørn-Helge et al., 2019), respectively.

#### 2.6.1 Random forest

RF is the machine learning ensemble classifier and has been applied in many fields (Breiman, 2001). By constructing multiple decision trees, the RF classifier has been considered as better performance than the single decision tree (Gandhi et al., 2018). In the current study, the randomforest function has been used to build RF classifiers. The number of classification trees and variables randomly selected for each node spilt have been set as ntree = 500, while mtry was optimized from one-third of the number of features minus 10 to plus 15. The relative importance of molecular fingerprints has also been calculated by the important function of the package.

#### 2.6.2 Support vector machine

SVM is a generalized linear classifier based on the principle of structural risk reduction for pattern recognition (Huang et al., 2018). It is well known as a supervised learning algorithm that analyzes data and recognizes patterns (Nedaie et al., 2018). In this study, the radial basis function (RBF) kernel was used for building the SVM classifier. Meanwhile, the random search method (Bergstra & Bengio, 2012) was also applied to optimize specific SVM parameters with the regularization parameter C and σ parameter by using the caret package, while C was from e^-2^ to e^6^, σ was from e^-7^ to e with the step of e^0.5^.

#### 2.6.3 Gradient boosting machine

GBM is also a tree-based machine learning model. It has been considered as a step-wise, additive type model which sequentially fits new-tree-based models (Golden et al., 2019). Meanwhile, it also has many advantages, especially worked well in practice (Cho et al., 2019). In this study, the total number of trees (n.trees) and the maximum depth of each tree (interaction. depth) have been optimized by using the caret package and have been set from 1 to 3000 and 1 to 10, respectively. Besides, shrinkage and n.minobsinnode were set as 0.005 and 10.

#### 2.6.4 Partial least square regression

PLS calculates a group of latent variables in connection with the output maximally and determines the relationship between the input and output data (Foodeh et al., 2020). It is a stretch of the multiple linear regression models and is widely used in many domains (Wu et al., 2020). Unlike multiple linear regression (MLR), it can handle the data with noisy, strongly collinear, and X-variables (Dong et al., 2018). In this study, n_components for PLS were optimized from 1 to the greatest features or sample sizes.

### 2.7. Model Evaluation

In order to test the predictive performance of the models, 5-fold cross-validation with 10 repeats has been used to evaluate the models. After randomly divided the original dataset into five equal subsets, four of them were used for training and the other was used for testing. Then the 5-fold cross-validation was repeated ten times to reduce the randomness. This cross-validation progress was performed 10 times with different random seeds of 2, 4, 8, 16, 32, 64, 128, 256, 512, and 1024. Then, average values were calculated to evaluate the prediction performance of the models.

Root-mean-squared error (RMSE), mean unsigned error (MUE), and R^2^ has been used to evaluate the predictive performance of the models. These indicators were calculated by the following formulas:

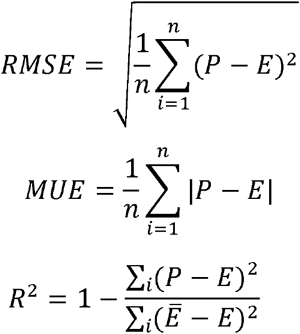

Where P, Ē, E, n represent predictive value, the average of experimental value, experimental value, and compound numbers, respectively.

## 3. Results and discussion

### 3.1. Feature selection

In this study, 203 compounds were collected from the previous study and divided into training and testing datasets with 80% to 20%, respectively. In order to build models to predict hERG cardiotoxicity, MDFP, Baseline2D, ECFP4, and PropertyFP have been calculated for the compounds in the dataset. Table 1 illustrated the number of features calculated from each type of molecular fingerprint and the detailed description of these features is shown in the supplementary files (Table S1 and Table S2). After the feature selection by RF-RFE, 11 and 6 features have been selected from MDFP and Baseline2D, respectively. Meanwhile, there were also 99 features selected from ECFP4 and 71 from PropertyFP. Percentage increase in MSE (%lncMSE) obtained by RF was used to evaluate the importance of features. Fig. 2 showed the top ten features (Baseline2D for six) which important to the prediction models. The results of MDFP showed that the number of hydrogen bonds between compounds and water has a significant effect on predicting hERG cardiotoxicity, followed by kinetic energy and surface area. Besides, the results of 2D molecular fingerprints indicated that the number of heavy atoms, number of O atoms (oxygens), and number of F atoms (fluorines) were the most important features in Baseline2D, while MolLog P in PropertyFP and 3218693969 in ECFP4. Above all, after calculating features in all molecular fingerprints, the following features have been selected as the most critical with heavyatoms, oxygens, fluorines, the median of hydrogen bonds, and 3218693969. These features may be played important roles in predicting the hERG cardiotoxicity and should be paid extra attention in the development of drug candidates.

**Table 1.** The number of features for the different molecular fingerprints.

| fingerprints | number of<br>features | number<br>of<br>selected<br>feature |
| --- | --- | --- |
| MDFP | 24 | 11 |
| Baseline2D | 10 | 6 |
| ECFP4 | 2298 | 99 |
| PropertyFP | 200 | 71 |

**Fig. 2.**
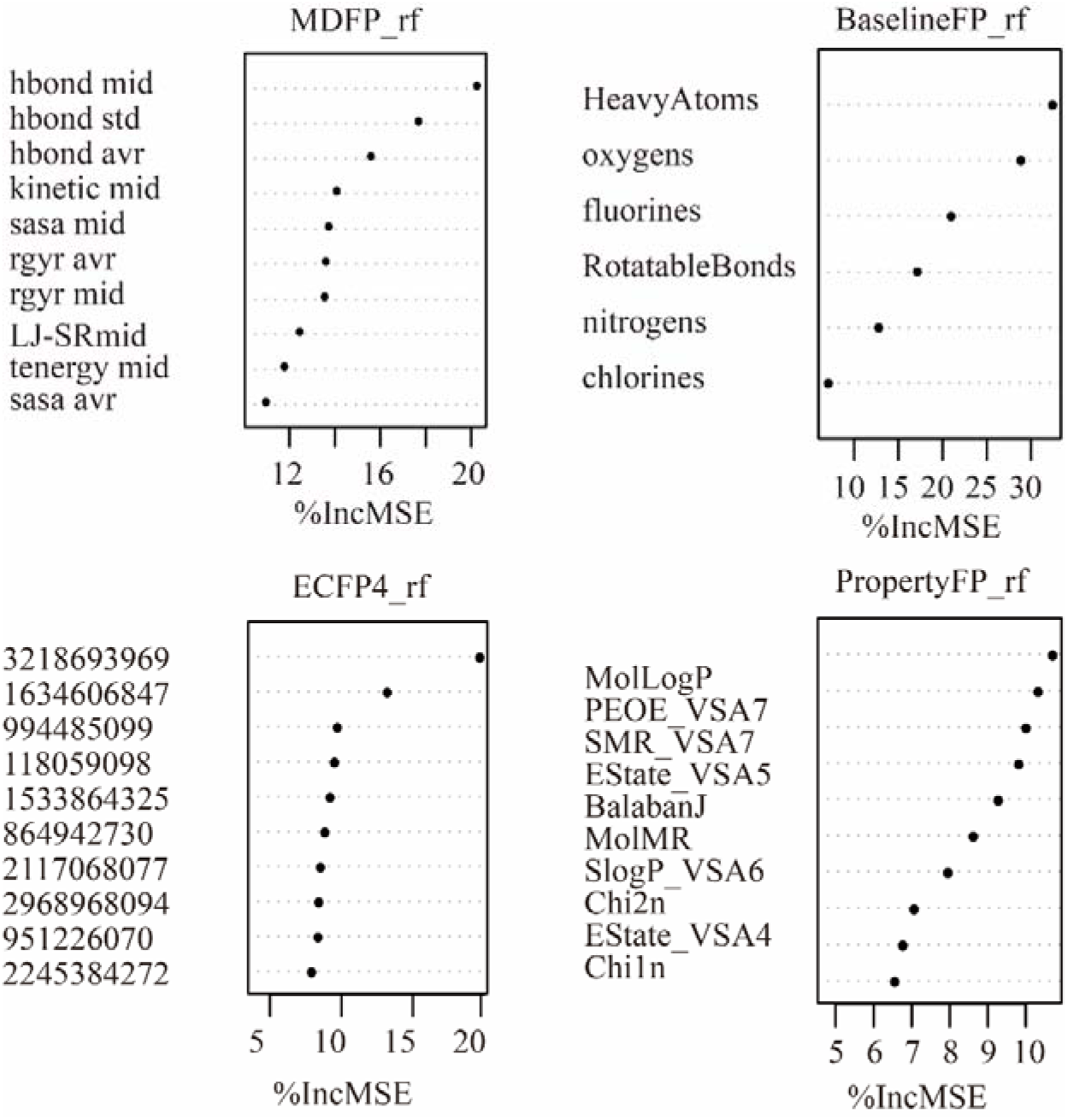
The most important features selected by RF-RFE from MDFP, Baseline2D, ECFP4, and PropertyFP fingerprints.

### 3.2. Prediction performance of the models

After performing feature selection, the GBM, PLS, RF, and SVM algorithms were used for generating ML models based on the resulting fingerprints. The performance of these machine learning models was evaluated by 10 times 5-fold cross-validation and their performances were presented in Table 2. The results showed that the RMSE of each machine learning model based on PropertyFP is the lowest, with a range of 0.860-0.960, followed by MDFP, with a range of 0.967-1.039, while ECFP4 and Baseline2D are poor quality. R^2^ and MUE also showed the same pattern. Table 3 illustrated the performance of these models which were used to predict the pIC50 of the molecules in the testing set. In general, the models show better RMSE values in the test set than in the 5-fold cross-validation, indicating that the model has not been overfitted. Meanwhile, compared with the models based on different molecular fingerprints, the performance in the testing set was similar, while Baseline2D was slightly better (RMSE=0.721 to 0.795) and MDFP also obtained a good performance (RMSE=0.755 to 0.819). These results indicated that MDFP can effectively predict the activity of hERG inhibitors, and the predictive performance of the MDFP was similar to the traditional molecular fingerprints.

**Table 2.** Cross-validation performance for models trained using different ML algorithms on the molecular fingerprints (MDFP, Baseline2D, ECFP4, PropertyFP). Performance metrics are represented as average and standard deviation of 10 times 5-fold cross-validation runs of different random seeds.

| fingerprint | ML models | RMSE | R<sup>2</sup> | MUE |
| --- | --- | --- | --- | --- |
| MDFP | GBM | 0.985±0.005 | 0.523±0.004 | 0.774±0.005 |
|  | PLS | 1.039±0.005 | 0.482±0.006 | 0.797±0.003 |
|  | RF | 0.977±0.005 | 0.534±0.006 | 0.768±0.004 |
|  | SVM | 0.967±0.007 | 0.541±0.006 | 0.745±0.007 |
| Baseline2D | GBM | 1.112±0.009 | 0.394±0.009 | 0.884±0.006 |
|  | PLS | 1.189±0.004 | 0.321±0.007 | 0.956±0.003 |
|  | RF | 1.036±0.011 | 0.465±0.011 | 0.813±0.008 |
|  | SVM | 1.014±0.006 | 0.492±0.006 | 0.791±0.006 |
| ECFP4 | GBM | 1.072±0.006 | 0.433±0.007 | 0.837±0.007 |
|  | PLS | 1.084±0.004 | 0.433±0.004 | 0.850±0.006 |
|  | RF | 1.043±0.004 | 0.464±0.004 | 0.827±0.004 |
|  | SVM | 1.009±0.004 | 0.497±0.004 | 0.800±0.004 |
| PropertyFP | GBM | 0.941±0.008 | 0.562±0.006 | 0.747±0.007 |
|  | PLS | 0.959±0.008 | 0.551±0.006 | 0.776±0.008 |
|  | RF | 0.960±0.004 | 0.559±0.004 | 0.763±0.005 |
|  | SVM | 0.860±0.006 | 0.634±0.006 | 0.676±0.009 |

**Table 3.** Cross-validation performance for models tested using different ML algorithms on the molecular fingerprints (MDFP, Baseline2D, ECFP4, PropertyFP). Performance metrics are represented as average and standard deviation of 10 times 5-fold cross-validation runs of different random seeds.

| fingerprint | ML models | RMSE | R<sup>2</sup> | MUE |
| --- | --- | --- | --- | --- |
| MDFP | GBM | 0.772±0.008 | 0.479±0.008 | 0.582±0.009 |
|  | PLS | 0.755±0 | 0.494±0 | 0.564±0 |
|  | RF | 0.819±0.011 | 0.398±0.012 | 0.570±0.006 |
|  | SVM | 0.802±0.010 | 0.458±0.007 | 0.586±0.005 |
|  | consensus | 0.745±0.005 | 0.495±0.005 | 0.524±0.003 |
| Baseline2D | GBM | 0.794±0.005 | 0.472±0.004 | 0.568±0.004 |
|  | PLS | 0.772±0.000 | 0.441±0.000 | 0.548±0.000 |
|  | RF | 0.795±0.015 | 0.423±0.015 | 0.545±0.011 |
|  | SVM | 0.721±0.005 | 0.525±0.005 | 0.520±0.011 |
|  | consensus | 0.713±0.003 | 0.520±0.004 | 0.507±0.002 |
| ECFP4 | GBM | 0.858±0.008 | 0.348±0.010 | 0.664±0.009 |
|  | PLS | 0.752±0.001 | 0.495±0.010 | 0.578±0.006 |
|  | RF | 0.865±0.009 | 0.315±0.011 | 0.635±0.011 |
|  | SVM | 0.737±0 | 0.491±0 | 0.553±0 |
|  | consensus | 0.761±0.001 | 0.457±0.003 | 0.571±0.003 |
| PropertyFP | GBM | 0.813±0.005 | 0.432±0.006 | 0.632±0.008 |
|  | PLS | 0.764±0.002 | 0.492±0.003 | 0.596±0.001 |
|  | RF | 0.709±0.006 | 0.529±0.009 | 0.540±0.006 |
|  | SVM | 0.761±0.035 | 0.488±0.040 | 0.605±0.033 |
|  | consensus | 0.730±0.008 | 0.508±0.010 | 0.560±0.008 |

The predictive performance of the MDFP model combined with other molecular fingerprints was also investigated in this study. Table 4 and Table 5 showed the performance of models in the 5-fold cross-validation sets and testing sets while MDFP combined with other molecular fingerprints, respectively. The results showed that the combination of MDFP and other molecular fingerprints can obtain a model with better prediction performance. For example, the model established by the single molecular fingerprint (MDFP or PropertyFP) in the 5-fold cross-validation had the best performance as PropertyFP-SVM (RMSE=0.860). However, the model established by multi-dimensional molecular fingerprints (MDFP and PropertyFP) was MDFP+PropertyFP-SVM (RMSE=0.837), which showed a better performance than using the single molecular fingerprints. Besides, models combining MDFP with other molecular fingerprints also showed better predictive performance in the testing set (Table 5), while the best model was the SVM model trained on MDFP++ (MDFP with all other fingerprints) (RMSE=0.696±0.015). These results illustrated that the performance of multi-dimensional molecular fingerprints was better than the single molecular fingerprints and MDFP may provide additional effective predictors for the prediction of hERG inhibitor activity.

**Table 4.** Cross-validation performance for models trained using different ML algorithms on the molecular fingerprints (MDFP + Baseline2D, MDFP + ECFP4, MDFP + PropertyFP, MDFP++). Performance metrics are represented as average and standard deviation of 10 times 5-fold cross-validation runs of different random seeds.

| fingerprint | ML models | RMSE | R<sup>2</sup> | MUE |
| --- | --- | --- | --- | --- |
| MDFP + Baseline2D | GBM | 0.991±0.005 | 0.516±0.005 | 0.767±0.005 |
|  | PLS | 1.068±0.007 | 0.458±0.004 | 0.820±0.004 |
|  | RF | 0.950±0.006 | 0.560±0.006 | 0.738±0.005 |
|  | SVM | 0.938±0.008 | 0.568±0.007 | 0.717±0.008 |
| MDFP + ECFP4 | GBM | 0.975±0.005 | 0.529±0.006 | 0.745±0.006 |
|  | PLS | 1.021±0.010 | 0.509±0.005 | 0.797±0.009 |
|  | RF | 0.945±0.005 | 0.566±0.004 | 0.740±0.005 |
|  | SVM | 0.935±0.005 | 0.569±0.004 | 0.740±0.005 |
| MDFP + PropertyFP | GBM | 0.915±0.008 | 0.585±0.006 | 0.722±0.009 |
|  | PLS | 0.948±0.011 | 0.568±0.009 | 0.754±0.011 |
|  | RF | 0.944±0.005 | 0.578±0.004 | 0.742±0.004 |
|  | SVM | 0.837±0.006 | 0.654±0.006 | 0.659±0.007 |
| MDFP++ | GBM | 0.920±0.008 | 0.580±0.006 | 0.723±0.008 |
|  | PLS | 0.958±0.007 | 0.556±0.005 | 0.754±0.007 |
|  | RF | 0.940±0.005 | 0.578±0.004 | 0.742±0.005 |
|  | SVM | 0.873±0.007 | 0.623±0.005 | 0.686±0.007 |

**Table 5.** Predictions were generated using different ML models trained on MDFP combined with multi-dimensional molecular fingerprints (MDFP + Baseline2D, MDFP + ECFP4, MDFP + PropertyFP, MDFP++) in test. MDFP++ including MDFP, Baseline2D, ECFP4, and PropertyFP.

| fingerprint | ML models | RMSE | R<sup>2</sup> | MUE |
| --- | --- | --- | --- | --- |
| MDFP + Baseline2D | GBM | 0.728±0.008 | 0.525±0.008 | 0.544±0.008 |
|  | PLS | 0.751±0.007 | 0.502±0.011 | 0.559±0.006 |
|  | RF | 0.789±0.009 | 0.427±0.011 | 0.560±0.008 |
|  | SVM | 0.781±0.003 | 0.494±0.002 | 0.551±0.001 |
|  | consensus | 0.721±0.003 | 0.524±0.003 | 0.518±0.003 |
| MDFP + ECFP4 | GBM | 0.758±0.007 | 0.491±0.007 | 0.569±0.004 |
|  | PLS | 0.702±0 | 0.555±0 | 0.535±0 |
|  | RF | 0.750±0.012 | 0.472±0.016 | 0.553±0.007 |
|  | SVM | 0.698±0.003 | 0.550±0.004 | 0.522±0.008 |
|  | consensus | 0.694±0.002 | 0.548±0.003 | 0.515±0.004 |
| MDFP + PropertyFP | GBM | 0.799±0.009 | 0.456±0.010 | 0.615±0.008 |
|  | PLS | 0.794±0.000 | 0.481±0.004 | 0.610±0.003 |
|  | RF | 0.709±0.008 | 0.527±0.011 | 0.549±0.009 |
|  | SVM | 0.723±0.011 | 0.518±0.012 | 0.578±0.012 |
|  | consensus | 0.719±0.003 | 0.523±0.003 | 0.554±0.003 |
| MDFP++ | GBM | 0.811±0.008 | 0.448±0.009 | 0.619±0.009 |
|  | PLS | 0.702±0 | 0.587±0 | 0.526±0 |
|  | RF | 0.718±0.010 | 0.513±0.014 | 0.554±0.006 |
|  | SVM | 0.696±0.015 | 0.564±0.011 | 0.516±0.017 |
|  | consensus | 0.695±0.004 | 0.557±0.004 | 0.518±0.003 |

In order to improve the prediction performance of the model, we further averaged the prediction results of the four machine learning models to obtain a consensus value. The prediction performance was shown in Table 3 and Table 5. Fig. 3 and Fig. 4 showed the predicted values vs experimental values for MDFP and MDFP++, respectively. The values of other molecular fingerprints have been demonstrated in the supplementary files (Fig. S1 to S6). It was found that the performance of consensus models was significantly better than the other models (except PropertyFP). Among the models established by a single molecular fingerprint, the consensus model based on Baseline2D had the highest accuracy (RMSE=0.713), while the consensus model based on MDFP also obtained a better RMSE of 0.745. Meanwhile, in the model based on the multi-dimensional molecular fingerprints, MDFP+ECFP4 and MDFP++ obtained high accuracy with RMSE of 0.694 and 0.695, respectively. These results indicated that the integrated model can obtain a better method for predicting the activity of hERG inhibitors.

**Fig. 3.**
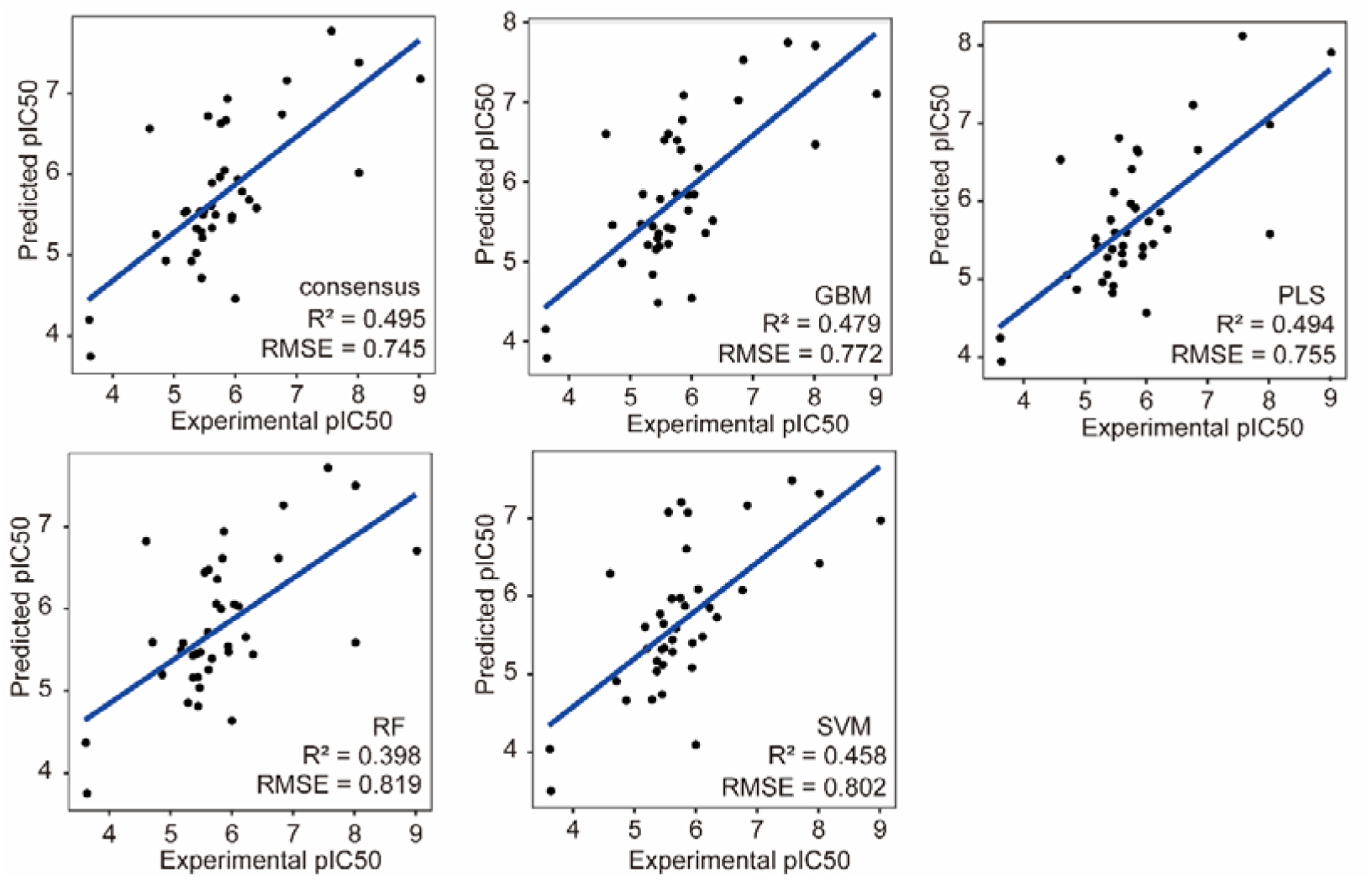
pIC50: The experimental values of the 10^th^ operation for the data set. Predictions were generated using consensus, GBM, PLS, RF, SVM trained on MDFP. The linear regression lines are shown in blue.

**Fig. 4.**
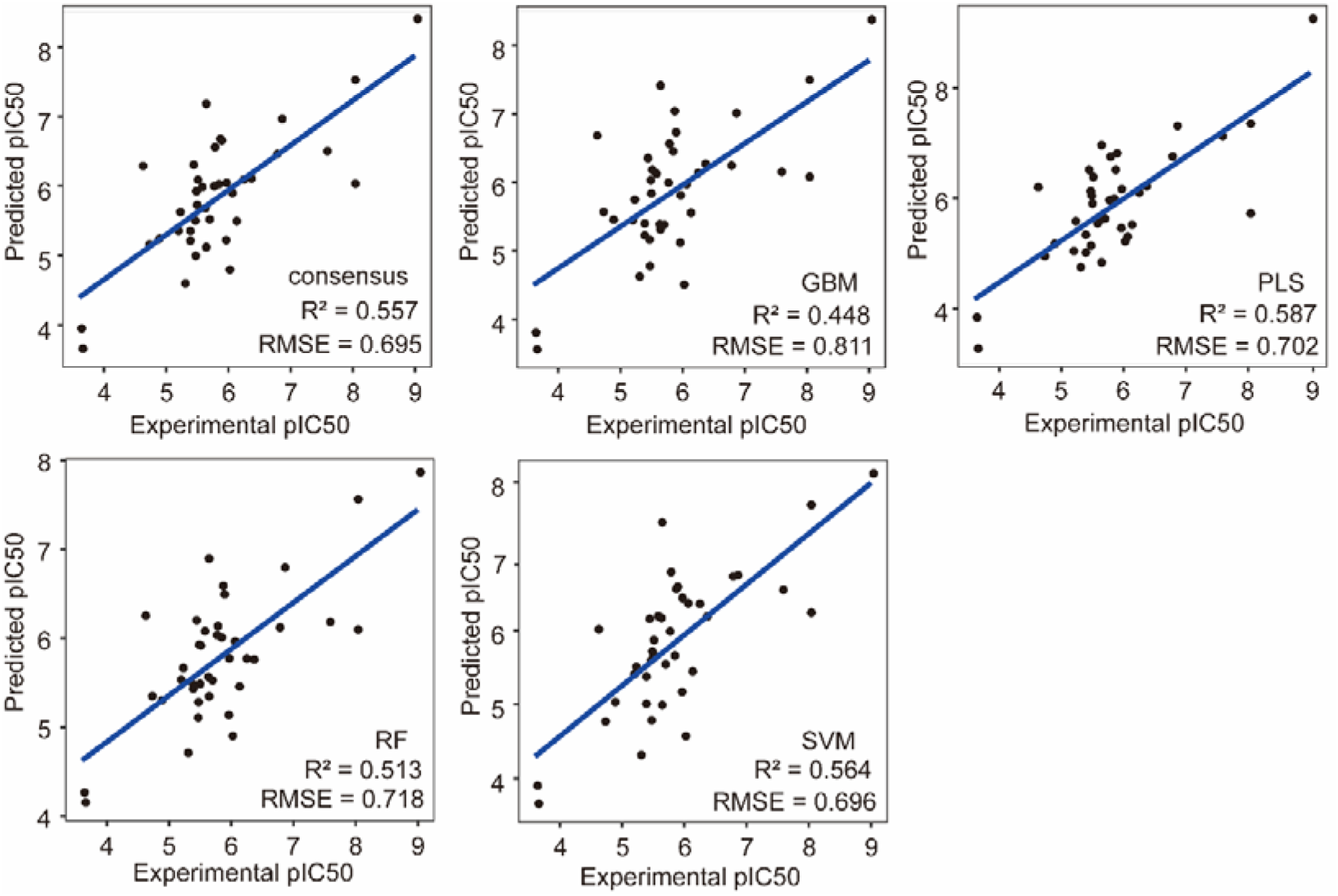
pIC50: The experimental values of the 10^th^ operation for the data set. Predictions were generated using consensus, GBM, PLS, RF, SVM trained on MDFP++. The linear regression lines are shown in blue. MDFP++ including MDFP, Baseline2D, ECFP4, and PropertyFP.

In summary, these results illustrated that the MDFP was effective compared with traditional molecular fingerprints and can truly be an alternative to the other molecular fingerprints. Meanwhile, the prediction accuracies of all ML models on multi-dimensional molecular fingerprints were better than the single molecular fingerprints in predicting the hERG cardiotoxicity. Besides, the integrated models showed the best prediction than the single models among most of the molecular fingerprints. Thus, the models obtained by multiple machine learning methods could be more accurate in predicting the hERG cardiotoxicity of compounds.

### 3.3. MDFP features associated with cardiotoxicity

To further reveal the contributions of fingerprint features associated with cardiotoxicity, the correlation coefficient has been used to determine the feature between MDFP and pIC50. Correlation is a measure of a monotonic association between 2 variables and Pearson’s correlation coefficient has become one of the most frequently used statistics (Armstrong, 2019). In this study, Pearson, Kendall, and Spearman correlations were used to evaluate the important features of MDFP with pIC50. Table 6 showed the correlation coefficient between the feature of MDFP and pIC50. The median of rgyr has been determined as the most relevant feature with pIC50 (Kendall = 0.35, Pearson = 0.51, and Spearman = 0.49), followed by the median of sasa and kinetic with the high correlation coefficient. These results showed the features which extracted from MDFP had strong correlations with pIC50 and can be used to predict cardiotoxicity in the future study.

**Table 6.** Correlation coefficient between the features of MDFP and pIC50.

| feature | kendall | pearson | spearman |
| --- | --- | --- | --- |
| rgyr mid | 0.35 | 0.51 | 0.49 |
| sasa mid | 0.30 | 0.41 | 0.43 |
| kinetic mid | 0.28 | 0.32 | 0.42 |
| LJ-SR mid | 0.28 | 0.25 | 0.41 |
| rgyr avr | 0.23 | 0.20 | 0.33 |
| sasa avr | 0.20 | 0.16 | 0.29 |
| sasa std | 0.17 | 0.01 | 0.25 |
| hbond avr | -0.08 | -0.14 | -0.12 |
| hbond std | -0.09 | -0.09 | -0.13 |
| hbond mid | -0.12 | -0.19 | -0.17 |
| tenergy mid | -0.28 | -0.42 | -0.41 |

### 3.4. Compared with other models

Recently, a couple of computational models have been developed for toxicity prediction. Among them, cardiotoxicity prediction has become a hotspot with multiple studies. Table 7 showed the comparisons between our model and other models for cardiotoxicity prediction. Compared with other models, the consensus model with MDFP and ECFP4 showed the lowest RMSE and MUE, with higher R^2^. Meanwhile, the molecular fingerprints of previous studies were used by only one dimension, which may prove that multi-dimensional fingerprints performed well in predicting the cardiotoxicity of hERG. Besides, although it was lower than QSAR-SVM, the consensus with MDFP still better than the other models as 0.745±0.005 (RMSE), which illustrated the advantages of MDFP. These findings showed that MDFP and multi-dimensional molecular fingerprints with machine learning methods can be an outstanding model in predicting cardiotoxicity.

**Table. 7.** Performance indicators of several cardiotoxicity prediction models reported in the literature.

| models | RMSE | $R^2$ | MUE | Reference |
| --- | --- | --- | --- | --- |
| QSAR-SVM | $0.79 \pm 0.05$ | $0.58 \pm 0.05$ | - | (Simeon & Jongkon, 2019) |
| QSAR-DNN | $0.90 \pm 0.06$ | $0.49 \pm 0.04$ | - | |
| MLR-Canvas | 1.186 | 0.191 | 0.941 | (Subramanian et al., 2016) |
| DNN-DeepChem | 1.03 | 0.351 | 0.763 |  |
| PLS-FFD | 1.07 | 0.48 | - | (Munawar et al., 2019) |
| consensus-MDFP | $0.745 \pm 0.045$ | $0.495 \pm 0.005$ | $0.524 \pm 0.003$ | |
| consensus-MDFP+ECFP4 | $0.694 \pm 0.002$ | $0.548 \pm 0.003$ | $0.515 \pm 0.004$ | |

## 4. Conclusion

In this study, MDFP and multi-dimensional molecular fingerprints were used for building machine learning models to predict the hERG cardiotoxicity of compounds. 203 compounds were firstly identified to establish the 5-fold cross-validation and testing datasets. Then molecular dynamic simulation has been used to generate molecular dynamic molecular fingerprints. Baseline2D, ECFP4, and PropertyFP were used to generate traditional molecular fingerprints. After that, critical features have been selected by RF-RFE and 4 machine learning algorithms, namely RF, SVM, GBM, and PLS were used for building predicting models based on the single fingerprints and multi-dimensional molecular fingerprints. Besides, the correlation between MDFP and pIC50 has also been surveyed. Results showed that MDFP has the potential to be an alternative choice of molecular fingerprints and multi-dimensional molecular fingerprints are better than single fingerprints in predicting cardiotoxicity. It also illustrated that the consensus model with MDFP and ECFP4 has the optimum prediction effect and hydrogen bonds are critically important in the models with MDFP. Our finding provides a new sight into the application of MDFP and multi-dimensional molecular fingerprints in predicting the hERG cardiotoxicity of compounds. Cell and animal experiments will be carried out to validate further.

## Acknowledgements

This study was supported by the National Natural Science Foundation of China (No. 82003655), the Key R&D Program of Liaoning Province (No. 2019JH2/10300041), Scientific Research Project from Department of Education of Liaoning Province (No. LQN201906), Shenyang Science and Technology Plan Project (No. 17-65-7-00, 19-302-3-04).

## Data Availability Statement

All data and models generated or used during the study appear in the submitted article.

## Author contributions

WZD, LZ, and HSL conceived the project, developed the prediction method, designed, and implemented the experiments, analyzed the result, and wrote the paper. YN, JSW, and XXX implemented the experiments, analyzed the result, and wrote the paper. SYH and SYL analyzed the result. All authors read and approved the final manuscript.

