## Supplementary material for "Combining Multi-Dimensional Molecular Fingerprints to Predict hERG Cardiotoxicity of Compounds": fig.S1

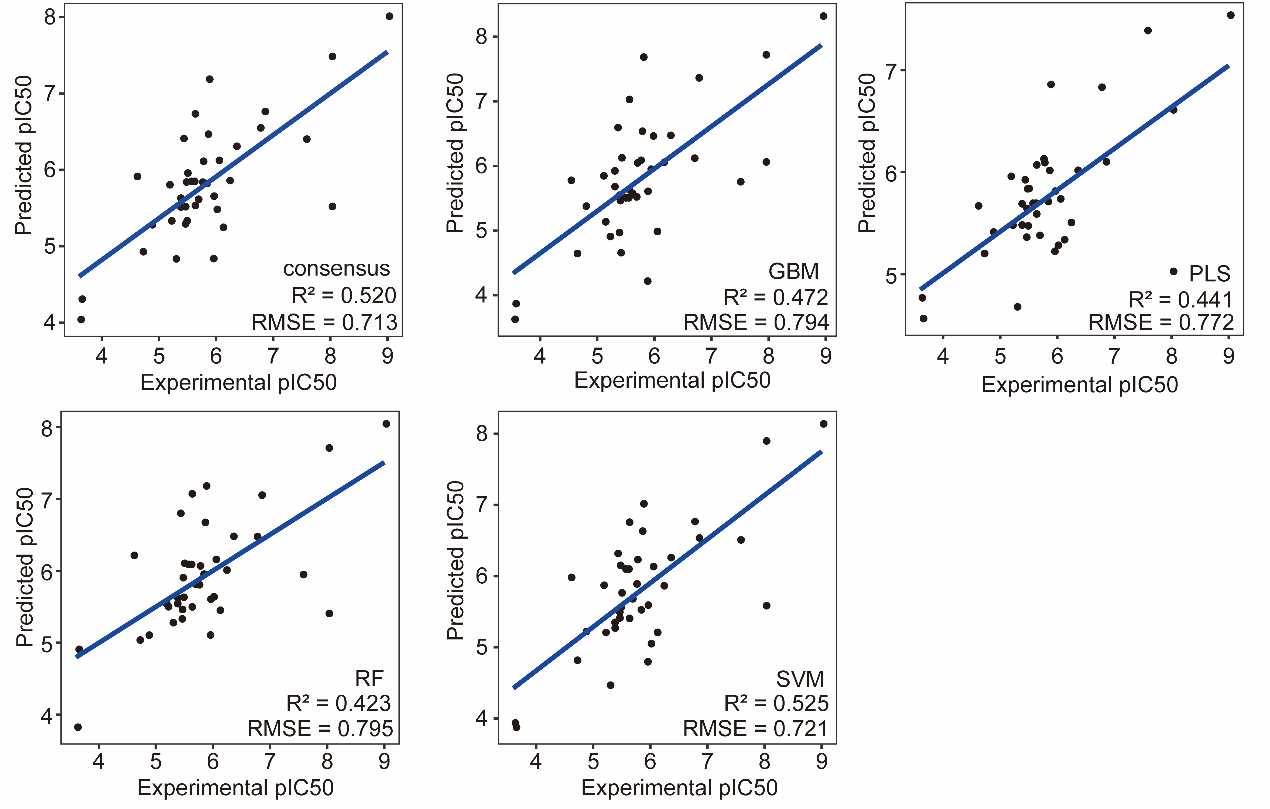

**Fig. S1.** pIC50: The experimental values of the 10^th^ operation for the data set. Predictions were generated using consensus, GBM, PLS, RF, SVM trained on Baseline2D. The linear regression lines are shown in blue.
