## Supplementary material for "Combining Multi-Dimensional Molecular Fingerprints to Predict hERG Cardiotoxicity of Compounds": table S1

**Table S1** Details of the features generated in different molecular fingerprints (MDFP, Baseline2D, PropertyFP). Selected features are marked with “+”.

| fingerprints | features | Description | feature selection |
| --- | --- | --- | --- |
| MDFP | sasa avr | The average of sasa | + |
|  | sasa std | The standard of sasa | + |
|  | sasa mid | The median of sasa | + |
|  | kinetic avr | The average of kinetic |  |
|  | kinetic std | The standard of kinetic |  |
|  | kinetic mid | The median of kinetic | + |
|  | LJ-14 avr | The average of LJ-14 |  |
|  | LJ-14 std | The standard of LJ-14 |  |
|  | LJ-14 mid | The median of LJ-14 |  |
|  | LJ-SR avr | The average of LJ-SR |  |
|  | LJ-SR std | The standard of LJ-SR |  |
|  | LJ-SR mid | The median of LJ-SR | + |
|  | tenergy avr | The average of tenergy |  |
|  | tenergy std | The standard of tenergy |  |
|  | tenergy mid | The median of tenergy | + |
|  | hbond avr | The average of hbond | + |
|  | hbond std | The standard of hbond | + |
|  | hbond mid | The median of hbond | + |
|  | rgyr avr | The average of rgyr | + |
|  | rgyr std | The standard of rgyr |  |
|  | rgyr mid | The median of rgyr | + |
|  | rmsd_avr | The average of rmsd |  |
|  | rmsd_std | The standard of rmsd |  |
|  | rmsd_mid | The median of rmsd |  |
| Baseline2D | HeavyAtoms |  | + |
|  | RotatableBonds |  | + |
|  | nitrogens |  | + |
|  | oxygens |  | + |
|  | fluorines |  | + |
|  | phosphorous |  |  |
|  | sulfurs |  |  |
|  | chlorines |  | + |
|  | bromines |  |  |
|  | iodines |  |  |
| PropertyFP | BalabanJ |  | + |
|  | BertzCT |  | + |
|  | Chi0 |  | + |
|  | Chi0n |  | + |
|  | Chi0v |  | + |
|  | Chi1 |  | + |
|  | Chi1n |  | + |
|  | Chi1v |  | + |
|  | Chi2n |  | + |
|  | Chi2v |  | + |
|  | Chi3n |  | + |
|  | Chi3v |  | + |
|  | Chi4n |  | + |
|  | Chi4v |  |  |
|  | EState_VSA1 |  | + |
|  | EState_VSA10 |  | + |
|  | EState_VSA11 |  |  |
|  | EState_VSA2 |  | + |
|  | EState_VSA3 |  |  |
|  | EState_VSA4 |  | + |
|  | EState_VSA5 |  | + |
|  | EState_VSA6 |  |  |
|  | EState_VSA7 |  |  |
|  | EState_VSA8 |  | + |
|  | EState_VSA9 |  |  |
|  | ExactMolWt |  | + |
|  | FpDensityMorgan1 |  | + |
|  | FpDensityMorgan2 |  |  |
|  | FpDensityMorgan3 |  |  |
|  | FractionCSP3 |  | + |
|  | HallKierAlpha |  | + |
|  | HeavyAtomCount |  | + |
|  | HeavyAtomMolWt |  | + |
|  | Ipc |  |  |
|  | Kappa1 |  | + |
|  | Kappa2 |  | + |
|  | Kappa3 |  | + |
|  | LabuteASA |  | + |
|  | MaxAbsEStateIndex |  | + |
|  | MaxAbsPartialCharge |  | + |
|  | MaxEStateIndex |  | + |
|  | MaxPartialCharge |  | + |
|  | MinAbsEStateIndex |  | + |
|  | MinAbsPartialCharge |  | + |
|  | MinEStateIndex |  | + |
|  | MinPartialCharge |  | + |
|  | MolLogP |  | + |
|  | MolMR |  | + |
|  | MolWt |  | + |
|  | NHOHCount |  |  |
|  | NOCount |  |  |
|  | NumAliphaticCarbocycles |  |  |
|  | NumAliphaticHeterocycles |  |  |
|  | NumAliphaticRings |  |  |
|  | NumAromaticCarbocycles |  | + |
|  | NumAromaticHeterocycles |  |  |
|  | NumAromaticRings |  | + |
|  | NumHAcceptors |  |  |
|  | NumHDonors |  |  |
|  | NumHeteroatoms |  |  |
|  | NumRadicalElectrons |  |  |
|  | NumRotatableBonds |  |  |
|  | NumSaturatedCarbocycles |  |  |
|  | NumSaturatedHeterocycles |  |  |
|  | NumSaturatedRings |  |  |
|  | NumValenceElectrons |  | + |
|  | PEOE_VSA1 |  | + |
|  | PEOE_VSA10 |  |  |
|  | PEOE_VSA11 |  |  |
|  | PEOE_VSA12 |  |  |
|  | PEOE_VSA13 |  |  |
|  | PEOE_VSA14 |  |  |
|  | PEOE_VSA2 |  |  |
|  | PEOE_VSA3 |  | + |
|  | PEOE_VSA4 |  |  |
|  | PEOE_VSA5 |  |  |
|  | PEOE_VSA6 |  | + |
|  | PEOE_VSA7 |  | + |
|  | PEOE_VSA8 |  | + |
|  | PEOE_VSA9 |  | + |
|  | RingCount |  |  |
|  | SMR_VSA1 |  | + |
|  | SMR_VSA10 |  | + |
|  | SMR_VSA2 |  |  |
|  | SMR_VSA3 |  | + |
|  | SMR_VSA4 |  | + |
|  | SMR_VSA5 |  | + |
|  | SMR_VSA6 |  | + |
|  | SMR_VSA7 |  | + |
|  | SMR_VSA8 |  |  |
|  | SMR_VSA9 |  |  |
|  | SlogP_VSA1 |  |  |
|  | SlogP_VSA10 |  | + |
|  | SlogP_VSA11 |  |  |
|  | SlogP_VSA12 |  |  |
|  | SlogP_VSA2 |  | + |
|  | SlogP_VSA3 |  |  |
|  | SlogP_VSA4 |  | + |
|  | SlogP_VSA5 |  | + |
|  | SlogP_VSA6 |  | + |
|  | SlogP_VSA7 |  |  |
|  | SlogP_VSA8 |  | + |
|  | SlogP_VSA9 |  |  |
|  | TPSA |  | + |
|  | VSA_EState1 |  |  |
|  | VSA_EState10 |  |  |
|  | VSA_EState2 |  |  |
|  | VSA_EState3 |  |  |
|  | VSA_EState4 |  |  |
|  | VSA_EState5 |  |  |
|  | VSA_EState6 |  |  |
|  | VSA_EState7 |  |  |
|  | VSA_EState8 |  | + |
|  | VSA_EState9 |  |  |
|  | fr_Al_COO |  |  |
|  | fr_Al_OH |  |  |
|  | fr_Al_OH_noTert |  |  |
|  | fr_ArN |  |  |
|  | fr_Ar_COO |  | + |
|  | fr_Ar_N |  |  |
|  | fr_Ar_NH |  |  |
|  | fr_Ar_OH |  |  |
|  | fr_COO |  | + |
|  | fr_COO2 |  | + |
|  | fr_C_O |  |  |
|  | fr_C_O_noCOO |  |  |
|  | fr_C_S |  |  |
|  | fr_HOCCN |  |  |
|  | fr_Imine |  |  |
|  | fr_NH0 |  |  |
|  | fr_NH1 |  |  |
|  | fr_NH2 |  |  |
|  | fr_N_O |  |  |
|  | fr_Ndealkylation1 |  |  |
|  | fr_Ndealkylation2 |  |  |
|  | fr_Nhpyrrole |  |  |
|  | fr_SH |  |  |
|  | fr_aldehyde |  |  |
|  | fr_alkyl_carbamate |  |  |
|  | fr_alkyl_halide |  |  |
|  | fr_allylic_oxid |  |  |
|  | fr_amide |  |  |
|  | fr_amidine |  |  |
|  | fr_aniline |  |  |
|  | fr_aryl_methyl |  |  |
|  | fr_azide |  |  |
|  | fr_azo |  |  |
|  | fr_barbitur |  |  |
|  | fr_benzene |  | + |
|  | fr_benzodiazepine |  |  |
|  | fr_bicyclic |  | + |
|  | fr_diazo |  |  |
|  | fr_dihydropyridine |  |  |
|  | fr_epoxide |  |  |
|  | fr_ester |  |  |
|  | fr_ether |  |  |
|  | fr_furan |  |  |
|  | fr_guanido |  |  |
|  | fr_halogen |  | + |
|  | fr_hdrzine |  |  |
|  | fr_hdrzone |  |  |
|  | fr_imidazole |  |  |
|  | fr_imide |  |  |
|  | fr_isocyan |  |  |
|  | fr_isothiocyan |  |  |
|  | fr_ketone |  |  |
|  | fr_ketone_Topliss |  |  |
|  | fr_lactam |  |  |
|  | fr_lactone |  |  |
|  | fr_methoxy |  |  |
|  | fr_morpholine |  |  |
|  | fr_nitrile |  |  |
|  | fr_nitro |  |  |
|  | fr_nitro_arom |  |  |
|  | fr_nitro_arom_nonortho |  |  |
|  | fr_nitroso |  |  |
|  | fr_oxazole |  |  |
|  | fr_oxime |  |  |
|  | fr_para_hydroxylation |  |  |
|  | fr_phenol |  |  |
|  | fr_phenol_noOrthoHbond |  |  |
|  | fr_phos_acid |  |  |
|  | fr_phos_ester |  |  |
|  | fr_piperdine |  |  |
|  | fr_piperzine |  |  |
|  | fr_priamide |  |  |
|  | fr_prisulfonamd |  |  |
|  | fr_pyridine |  |  |
|  | fr_quatN |  |  |
|  | fr_sulfide |  |  |
|  | fr_sulfonamd |  |  |
|  | fr_sulfone |  |  |
|  | fr_term_acetylene |  |  |
|  | fr_tetrazole |  |  |
|  | fr_thiazole |  |  |
|  | fr_thiocyan |  |  |
|  | fr_thiophene |  |  |
|  | fr_unbrch_alkane |  |  |
|  | fr_urea |  |  |
|  | qed |  | + |
