## Supplementary material for "Combining Multi-Dimensional Molecular Fingerprints to Predict hERG Cardiotoxicity of Compounds": table S2

**Table S2** All the features were selected on importance by %lncMSE in ECFP4 molecular fingerprints.

| fingerprints | features | feature selection |
| --- | --- | --- |
| ECFP4 | 852462 | 3624155 |
|  | 3624155 | 10565946 |
|  | 5851216 | 19852674 |
|  | 8212726 | 98513984 |
|  | 8757308 | 118059098 |
|  | 9713885 | 132611095 |
|  | 10565946 | 256204769 |
|  | 13251412 | 310388129 |
|  | 14919950 | 409801989 |
|  | 15839042 | 422715066 |
|  | 19852674 | 572760528 |
|  | 20567188 | 586636525 |
|  | 21464752 | 603510687 |
|  | 26234434 | 787069595 |
|  | 30707086 | 847957139 |
|  | 31076933 | 847961216 |
|  | 34006608 | 848128881 |
|  | 35067595 | 864662311 |
|  | 36538691 | 864674487 |
|  | 37182385 | 864942730 |
|  | 37269444 | 882399112 |
|  | 38478160 | 934550081 |
|  | 43575877 | 951226070 |
|  | 43606954 | 994485099 |
|  | 46138220 | 1016841875 |
|  | 49003182 | 1083852209 |
|  | 50279957 | 1100037548 |
|  | 53971451 | 1101907775 |
|  | 54861895 | 1109942725 |
|  | 54936104 | 1167322652 |
|  | 55489393 | 1173125914 |
|  | 62093196 | 1313967653 |
|  | 65918436 | 1376881293 |
|  | 65941455 | 1510328189 |
|  | 67332941 | 1510461303 |
|  | 69470421 | 1533864325 |
|  | 73766099 | 1542633699 |
|  | 74537039 | 1583799011 |
|  | 81315441 | 1634606847 |
|  | 81456112 | 1637836422 |
|  | 83000178 | 1790668568 |
|  | 86544075 | 2021812431 |
|  | 88831783 | 2036328569 |
|  | 92620239 | 2041434490 |
|  | 96001975 | 2092489639 |
|  | 96783414 | 2117068077 |
|  | 98513984 | 2142032900 |
|  | 110033661 | 2228558432 |
|  | 115579364 | 2245277810 |
|  | 116091479 | 2245384272 |
|  | 116374980 | 2246340824 |
|  | 117265359 | 2246699815 |
|  | 117396011 | 2246728737 |
|  | 118059098 | 2257970297 |
|  | 119221004 | 2435680164 |
|  | 119562847 | 2463614838 |
|  | 122001321 | 2594220197 |
|  | 128522177 | 2763854213 |
|  | 132611095 | 2784506312 |
|  | 134920792 | 2803848648 |
|  | 136353966 | 2811394787 |
|  | 136366266 | 2832976762 |
|  | 137636017 | 2968968094 |
|  | 137723646 | 2976033787 |
|  | 143628737 | 2976816164 |
|  | 144568923 | 3121777292 |
|  | 144870008 | 3160601726 |
|  | 145374537 | 3182824521 |
|  | 149653088 | 3189457552 |
|  | 160260385 | 3217380708 |
|  | 160723288 | 3218693969 |
|  | 161963127 | 3315826729 |
|  | 165848974 | 3334525955 |
|  | 166544320 | 3337745083 |
|  | 170741831 | 3452535345 |
|  | 171200514 | 3491109101 |
|  | 172238518 | 3542456614 |
|  | 172337835 | 3594646603 |
|  | 176036124 | 3631761933 |
|  | 176087340 | 3657471097 |
|  | 179318893 | 3661262542 |
|  | 180633200 | 3684013815 |
|  | 185358682 | 3692055567 |
|  | 190140222 | 3776905034 |
|  | 190406487 | 3818546315 |
|  | 191014185 | 3820826669 |
|  | 193481629 | 3848368466 |
|  | 195323148 | 3892941493 |
|  | 195764383 | 3907878801 |
|  | 197218356 | 3933537673 |
|  | 200184615 | 3965194470 |
|  | 202865053 | 3982076256 |
|  | 204065626 | 3983062349 |
|  | 204549064 | 3997240246 |
|  | 206372905 | 3999906991 |
|  | 207732580 | 4036277955 |
|  | 211458732 | 4042373501 |
|  | 211903143 | 4055698890 |
|  | 212358896 | 4197146180 |
|  | 214126897 |  |
|  | 215117572 |  |
|  | 218474789 |  |
|  | 219610753 |  |
|  | 219906052 |  |
|  | 219974458 |  |
|  | 222262242 |  |
|  | 228862830 |  |
|  | 229197718 |  |
|  | 230478851 |  |
|  | 232011924 |  |
|  | 232497375 |  |
|  | 237615532 |  |
|  | 239295302 |  |
|  | 239490685 |  |
|  | 240803077 |  |
|  | 242694095 |  |
|  | 244052397 |  |
|  | 250769854 |  |
|  | 250814365 |  |
|  | 252874892 |  |
|  | 253115797 |  |
|  | 255802399 |  |
|  | 256204769 |  |
|  | 257851290 |  |
|  | 264762917 |  |
|  | 264906369 |  |
|  | 265892994 |  |
|  | 266249624 |  |
|  | 266675433 |  |
|  | 269171368 |  |
|  | 270475737 |  |
|  | 271903915 |  |
|  | 273756909 |  |
|  | 276046393 |  |
|  | 283306526 |  |
|  | 285599556 |  |
|  | 287232915 |  |
|  | 288315894 |  |
|  | 288722959 |  |
|  | 289080917 |  |
|  | 289801995 |  |
|  | 292399449 |  |
|  | 293645588 |  |
|  | 293924501 |  |
|  | 293971392 |  |
|  | 298300585 |  |
|  | 298374928 |  |
|  | 299167261 |  |
|  | 302572264 |  |
|  | 303135741 |  |
|  | 304034656 |  |
|  | 304986910 |  |
|  | 305828102 |  |
|  | 306288037 |  |
|  | 307525970 |  |
|  | 308810678 |  |
|  | 310388129 |  |
|  | 310913359 |  |
|  | 311972411 |  |
|  | 313683681 |  |
|  | 315011794 |  |
|  | 319552149 |  |
|  | 320205079 |  |
|  | 321337029 |  |
|  | 326707068 |  |
|  | 327061258 |  |
|  | 328936174 |  |
|  | 330223697 |  |
|  | 332354558 |  |
|  | 335270011 |  |
|  | 337627472 |  |
|  | 345392702 |  |
|  | 347565724 |  |
|  | 351819320 |  |
|  | 357342195 |  |
|  | 365002352 |  |
|  | 368531313 |  |
|  | 368713488 |  |
|  | 369315086 |  |
|  | 374795611 |  |
|  | 374977434 |  |
|  | 379825674 |  |
|  | 384152397 |  |
|  | 385544376 |  |
|  | 386092920 |  |
|  | 390426252 |  |
|  | 396311007 |  |
|  | 398568954 |  |
|  | 400500269 |  |
|  | 405578005 |  |
|  | 405900472 |  |
|  | 408293823 |  |
|  | 409801989 |  |
|  | 410310586 |  |
|  | 416356657 |  |
|  | 422165373 |  |
|  | 422715066 |  |
|  | 423609967 |  |
|  | 424376859 |  |
|  | 426242490 |  |
|  | 427867059 |  |
|  | 431195058 |  |
|  | 431202968 |  |
|  | 433289869 |  |
|  | 433773193 |  |
|  | 434423882 |  |
|  | 434543776 |  |
|  | 435232729 |  |
|  | 437805903 |  |
|  | 438669819 |  |
|  | 441521794 |  |
|  | 443741425 |  |
|  | 446551469 |  |
|  | 447529997 |  |
|  | 448302737 |  |
|  | 450365582 |  |
|  | 452417285 |  |
|  | 456978550 |  |
|  | 459358826 |  |
|  | 463959832 |  |
|  | 467071127 |  |
|  | 467603055 |  |
|  | 468055343 |  |
|  | 469168129 |  |
|  | 472281497 |  |
|  | 476060908 |  |
|  | 478352517 |  |
|  | 480680885 |  |
|  | 482332985 |  |
|  | 483001034 |  |
|  | 483752110 |  |
|  | 484705467 |  |
|  | 485463469 |  |
|  | 485641844 |  |
|  | 486030630 |  |
|  | 486705461 |  |
|  | 487673803 |  |
|  | 491884665 |  |
|  | 493191496 |  |
|  | 496179091 |  |
|  | 496445357 |  |
|  | 497870099 |  |
|  | 501836124 |  |
|  | 503074166 |  |
|  | 509026204 |  |
|  | 509217717 |  |
|  | 511170092 |  |
|  | 516041383 |  |
|  | 517457164 |  |
|  | 518687148 |  |
|  | 519369723 |  |
|  | 523461156 |  |
|  | 525850584 |  |
|  | 530246988 |  |
|  | 530656108 |  |
|  | 535828179 |  |
|  | 540046244 |  |
|  | 543897685 |  |
|  | 544755086 |  |
|  | 544863012 |  |
|  | 548571000 |  |
|  | 548665696 |  |
|  | 550081331 |  |
|  | 550276949 |  |
|  | 550953794 |  |
|  | 551287920 |  |
|  | 552722929 |  |
|  | 553412256 |  |
|  | 556594412 |  |
|  | 565690435 |  |
|  | 568634294 |  |
|  | 571127535 |  |
|  | 571978829 |  |
|  | 572760528 |  |
|  | 574797690 |  |
|  | 577408712 |  |
|  | 579286581 |  |
|  | 579898624 |  |
|  | 584893129 |  |
|  | 585340062 |  |
|  | 586636525 |  |
|  | 586976137 |  |
|  | 589930590 |  |
|  | 592593828 |  |
|  | 592655526 |  |
|  | 600377156 |  |
|  | 600629739 |  |
|  | 601159014 |  |
|  | 602889953 |  |
|  | 603510687 |  |
|  | 603997856 |  |
|  | 605268295 |  |
|  | 606428793 |  |
|  | 607100956 |  |
|  | 610814047 |  |
|  | 611982871 |  |
|  | 616374597 |  |
|  | 616574971 |  |
|  | 618008133 |  |
|  | 618671879 |  |
|  | 622233516 |  |
|  | 623728226 |  |
|  | 625123882 |  |
|  | 628583830 |  |
|  | 631553871 |  |
|  | 636971230 |  |
|  | 638050764 |  |
|  | 642367021 |  |
|  | 643572069 |  |
|  | 645084142 |  |
|  | 649201019 |  |
|  | 649517798 |  |
|  | 662511025 |  |
|  | 662595032 |  |
|  | 665316820 |  |
|  | 667123616 |  |
|  | 667404468 |  |
|  | 667497574 |  |
|  | 674450788 |  |
|  | 675765711 |  |
|  | 679391948 |  |
|  | 684208217 |  |
|  | 685125387 |  |
|  | 685706799 |  |
|  | 693302733 |  |
|  | 694069322 |  |
|  | 696250882 |  |
|  | 696361425 |  |
|  | 698112012 |  |
|  | 698893736 |  |
|  | 699795807 |  |
|  | 701541392 |  |
|  | 704691639 |  |
|  | 706972984 |  |
|  | 711358894 |  |
|  | 711814864 |  |
|  | 711849114 |  |
|  | 712145326 |  |
|  | 714883344 |  |
|  | 715404902 |  |
|  | 716189338 |  |
|  | 716321151 |  |
|  | 717151242 |  |
|  | 717512901 |  |
|  | 717723595 |  |
|  | 719193781 |  |
|  | 719320964 |  |
|  | 719832797 |  |
|  | 720956516 |  |
|  | 721333722 |  |
|  | 725338437 |  |
|  | 725727510 |  |
|  | 729796064 |  |
|  | 731987238 |  |
|  | 737524545 |  |
|  | 737872620 |  |
|  | 738659967 |  |
|  | 739021901 |  |
|  | 739041005 |  |
|  | 740717125 |  |
|  | 745024969 |  |
|  | 746386425 |  |
|  | 746613721 |  |
|  | 747735251 |  |
|  | 754166207 |  |
|  | 756562620 |  |
|  | 762909776 |  |
|  | 766491448 |  |
|  | 773457308 |  |
|  | 773607102 |  |
|  | 778085759 |  |
|  | 780020950 |  |
|  | 781912670 |  |
|  | 784131575 |  |
|  | 787045417 |  |
|  | 787069595 |  |
|  | 787341104 |  |
|  | 787495672 |  |
|  | 789496286 |  |
|  | 792968423 |  |
|  | 794201745 |  |
|  | 798961065 |  |
|  | 799373156 |  |
|  | 800085213 |  |
|  | 805455350 |  |
|  | 805987832 |  |
|  | 808416955 |  |
|  | 817588605 |  |
|  | 818776141 |  |
|  | 819348444 |  |
|  | 822222489 |  |
|  | 823820999 |  |
|  | 827346468 |  |
|  | 829754261 |  |
|  | 830340805 |  |
|  | 831281868 |  |
|  | 831463242 |  |
|  | 833123915 |  |
|  | 833901780 |  |
|  | 834846795 |  |
|  | 836300471 |  |
|  | 837992725 |  |
|  | 838905077 |  |
|  | 838909898 |  |
|  | 839602758 |  |
|  | 840271901 |  |
|  | 840355693 |  |
|  | 844021776 |  |
|  | 844521347 |  |
|  | 845764036 |  |
|  | 846808146 |  |
|  | 847028072 |  |
|  | 847307936 |  |
|  | 847336149 |  |
|  | 847433064 |  |
|  | 847934112 |  |
|  | 847957139 |  |
|  | 847961216 |  |
|  | 848127915 |  |
|  | 848128881 |  |
|  | 849275503 |  |
|  | 854241849 |  |
|  | 858362104 |  |
|  | 859793819 |  |
|  | 859799282 |  |
|  | 860193140 |  |
|  | 864503681 |  |
|  | 864662311 |  |
|  | 864674487 |  |
|  | 864942730 |  |
|  | 864942795 |  |
|  | 868877654 |  |
|  | 869121947 |  |
|  | 871877728 |  |
|  | 871944527 |  |
|  | 872351080 |  |
|  | 873832266 |  |
|  | 874637027 |  |
|  | 875116816 |  |
|  | 876016289 |  |
|  | 876335404 |  |
|  | 876651869 |  |
|  | 880794658 |  |
|  | 881070323 |  |
|  | 881979466 |  |
|  | 882399112 |  |
|  | 882686432 |  |
|  | 884707938 |  |
|  | 885026269 |  |
|  | 885223974 |  |
|  | 889594580 |  |
|  | 892756502 |  |
|  | 894436332 |  |
|  | 902470469 |  |
|  | 905544157 |  |
|  | 906905665 |  |
|  | 910184811 |  |
|  | 910450790 |  |
|  | 911833712 |  |
|  | 912632640 |  |
|  | 913917326 |  |
|  | 913947419 |  |
|  | 914762037 |  |
|  | 915017141 |  |
|  | 916634498 |  |
|  | 920763749 |  |
|  | 921426030 |  |
|  | 921629099 |  |
|  | 926215114 |  |
|  | 927052640 |  |
|  | 927745747 |  |
|  | 932551375 |  |
|  | 932711668 |  |
|  | 932712697 |  |
|  | 934550081 |  |
|  | 937526175 |  |
|  | 940732351 |  |
|  | 950023157 |  |
|  | 951226070 |  |
|  | 951239203 |  |
|  | 951469037 |  |
|  | 954562416 |  |
|  | 954800030 |  |
|  | 957251995 |  |
|  | 957810395 |  |
|  | 958885167 |  |
|  | 960667422 |  |
|  | 962042210 |  |
|  | 963029399 |  |
|  | 964520091 |  |
|  | 964913175 |  |
|  | 966311396 |  |
|  | 970102735 |  |
|  | 970433791 |  |
|  | 974654488 |  |
|  | 978432910 |  |
|  | 979903539 |  |
|  | 980020035 |  |
|  | 980291093 |  |
|  | 986616778 |  |
|  | 987235844 |  |
|  | 988854583 |  |
|  | 989257658 |  |
|  | 989280568 |  |
|  | 989543450 |  |
|  | 992462230 |  |
|  | 994237755 |  |
|  | 994485099 |  |
|  | 994494548 |  |
|  | 996913272 |  |
|  | 999334238 |  |
|  | 999602603 |  |
|  | 1000622147 | |
|  | 1006167676 | |
|  | 1007283054 | |
|  | 1007683032 | |
|  | 1009042072 | |
|  | 1010142783 | |
|  | 1010672141 | |
|  | 1012780892 | |
|  | 1016053650 | |
|  | 1016093624 | |
|  | 1016841875 | |
|  | 1021101407 | |
|  | 1021342803 | |
|  | 1022974558 | |
|  | 1025559095 | |
|  | 1026654305 | |
|  | 1026928756 | |
|  | 1028073125 | |
|  | 1031129086 | |
|  | 1036411339 | |
|  | 1040143037 | |
|  | 1040797185 | |
|  | 1041832133 | |
|  | 1041876320 | |
|  | 1044405971 | |
|  | 1044537682 | |
|  | 1045831773 | |
|  | 1046110189 | |
|  | 1046313701 | |
|  | 1047069352 | |
|  | 1051848033 | |
|  | 1054767590 | |
|  | 1055081950 | |
|  | 1066486397 | |
|  | 1067305901 | |
|  | 1067317301 | |
|  | 1071810269 | |
|  | 1077373113 | |
|  | 1078211001 | |
|  | 1078276907 | |
|  | 1083852209 | |
|  | 1084224517 | |
|  | 1085526790 | |
|  | 1086305639 | |
|  | 1086480009 | |
|  | 1089584628 | |
|  | 1089888822 | |
|  | 1090103393 | |
|  | 1090571167 | |
|  | 1091346796 | |
|  | 1094126528 | |
|  | 1094481266 | |
|  | 1095816046 | |
|  | 1097652442 | |
|  | 1098159556 | |
|  | 1099011003 | |
|  | 1100037548 | |
|  | 1101399647 | |
|  | 1101907775 | |
|  | 1103426160 | |
|  | 1105397096 | |
|  | 1105669173 | |
|  | 1109942725 | |
|  | 1112379768 | |
|  | 1112821593 | |
|  | 1114299067 | |
|  | 1116101896 | |
|  | 1117976776 | |
|  | 1118326103 | |
|  | 1126371749 | |
|  | 1130345056 | |
|  | 1134608389 | |
|  | 1135286194 | |
|  | 1136130381 | |
|  | 1138084776 | |
|  | 1138487535 | |
|  | 1140133616 | |
|  | 1145656637 | |
|  | 1146516343 | |
|  | 1147977085 | |
|  | 1153219603 | |
|  | 1153303704 | |
|  | 1161921078 | |
|  | 1163692816 | |
|  | 1165927732 | |
|  | 1167268241 | |
|  | 1167322652 | |
|  | 1169545996 | |
|  | 1173125914 | |
|  | 1173528643 | |
|  | 1174508388 | |
|  | 1181021693 | |
|  | 1182622762 | |
|  | 1184237381 | |
|  | 1184436594 | |
|  | 1185093695 | |
|  | 1186654798 | |
|  | 1187730664 | |
|  | 1189967413 | |
|  | 1191428569 | |
|  | 1192366885 | |
|  | 1195162643 | |
|  | 1199016914 | |
|  | 1199792329 | |
|  | 1207044812 | |
|  | 1207774339 | |
|  | 1211233089 | |
|  | 1212160130 | |
|  | 1213407188 | |
|  | 1214058429 | |
|  | 1214060489 | |
|  | 1215457691 | |
|  | 1216095805 | |
|  | 1217367676 | |
|  | 1219508249 | |
|  | 1219984293 | |
|  | 1221464654 | |
|  | 1222399993 | |
|  | 1222707958 | |
|  | 1226975992 | |
|  | 1228263945 | |
|  | 1228528465 | |
|  | 1229325558 | |
|  | 1231564189 | |
|  | 1233505769 | |
|  | 1233810244 | |
|  | 1235335132 | |
|  | 1235524787 | |
|  | 1236033658 | |
|  | 1236342681 | |
|  | 1236995907 | |
|  | 1239899672 | |
|  | 1241059305 | |
|  | 1244401270 | |
|  | 1246481034 | |
|  | 1247998866 | |
|  | 1249334698 | |
|  | 1250165012 | |
|  | 1251599658 | |
|  | 1251717127 | |
|  | 1255403697 | |
|  | 1256751600 | |
|  | 1256946769 | |
|  | 1257718710 | |
|  | 1260363809 | |
|  | 1264267632 | |
|  | 1266193593 | |
|  | 1266789729 | |
|  | 1267137497 | |
|  | 1267207243 | |
|  | 1267407792 | |
|  | 1271427701 | |
|  | 1271688533 | |
|  | 1271691605 | |
|  | 1272192549 | |
|  | 1276414529 | |
|  | 1276993226 | |
|  | 1279513460 | |
|  | 1279636133 | |
|  | 1279665880 | |
|  | 1280488931 | |
|  | 1285475934 | |
|  | 1286704427 | |
|  | 1287954200 | |
|  | 1289643292 | |
|  | 1292701557 | |
|  | 1301991264 | |
|  | 1305042253 | |
|  | 1306816608 | |
|  | 1308598964 | |
|  | 1309208283 | |
|  | 1312131881 | |
|  | 1313764286 | |
|  | 1313811741 | |
|  | 1313967653 | |
|  | 1315950583 | |
|  | 1316507708 | |
|  | 1320519598 | |
|  | 1320875999 | |
|  | 1322529363 | |
|  | 1323452743 | |
|  | 1323467736 | |
|  | 1323887020 | |
|  | 1324775727 | |
|  | 1325617440 | |
|  | 1327153794 | |
|  | 1336346143 | |
|  | 1340247152 | |
|  | 1342487149 | |
|  | 1344062198 | |
|  | 1348250504 | |
|  | 1348783936 | |
|  | 1349242618 | |
|  | 1349263792 | |
|  | 1349266638 | |
|  | 1349287913 | |
|  | 1349611886 | |
|  | 1350922703 | |
|  | 1352399629 | |
|  | 1354803640 | |
|  | 1362202978 | |
|  | 1362425715 | |
|  | 1362518133 | |
|  | 1363569421 | |
|  | 1363660037 | |
|  | 1369588494 | |
|  | 1370727332 | |
|  | 1371111668 | |
|  | 1376853888 | |
|  | 1376881293 | |
|  | 1378792052 | |
|  | 1378815686 | |
|  | 1381999759 | |
|  | 1382479327 | |
|  | 1390505218 | |
|  | 1390916045 | |
|  | 1391940372 | |
|  | 1392283410 | |
|  | 1392646552 | |
|  | 1395869423 | |
|  | 1396119199 | |
|  | 1396348621 | |
|  | 1397494279 | |
|  | 1397786340 | |
|  | 1398128024 | |
|  | 1398897557 | |
|  | 1399598682 | |
|  | 1399966818 | |
|  | 1402541001 | |
|  | 1403876697 | |
|  | 1407821339 | |
|  | 1408978711 | |
|  | 1409732253 | |
|  | 1410865100 | |
|  | 1421345390 | |
|  | 1423151108 | |
|  | 1425045427 | |
|  | 1428903162 | |
|  | 1429182865 | |
|  | 1429883190 | |
|  | 1433386768 | |
|  | 1435511471 | |
|  | 1440420545 | |
|  | 1441199729 | |
|  | 1441523996 | |
|  | 1445852595 | |
|  | 1446204275 | |
|  | 1449131080 | |
|  | 1449220749 | |
|  | 1450335512 | |
|  | 1451062712 | |
|  | 1455164111 | |
|  | 1455165215 | |
|  | 1456084163 | |
|  | 1458875980 | |
|  | 1460642244 | |
|  | 1461502191 | |
|  | 1464119786 | |
|  | 1466409066 | |
|  | 1468316360 | |
|  | 1470963619 | |
|  | 1472142383 | |
|  | 1475039305 | |
|  | 1477785326 | |
|  | 1477870366 | |
|  | 1478031619 | |
|  | 1479995930 | |
|  | 1480134744 | |
|  | 1485981353 | |
|  | 1486598263 | |
|  | 1491567894 | |
|  | 1494778871 | |
|  | 1499507695 | |
|  | 1500223086 | |
|  | 1500549366 | |
|  | 1500702723 | |
|  | 1500929892 | |
|  | 1501125405 | |
|  | 1502513189 | |
|  | 1502988933 | |
|  | 1505769767 | |
|  | 1506563592 | |
|  | 1506993418 | |
|  | 1507955614 | |
|  | 1510328189 | |
|  | 1510355397 | |
|  | 1510461303 | |
|  | 1513049439 | |
|  | 1514095308 | |
|  | 1514987235 | |
|  | 1525423190 | |
|  | 1526453647 | |
|  | 1528282272 | |
|  | 1529733929 | |
|  | 1532069362 | |
|  | 1532878780 | |
|  | 1533767736 | |
|  | 1533864325 | |
|  | 1534862878 | |
|  | 1535166686 | |
|  | 1536224221 | |
|  | 1540337387 | |
|  | 1542631284 | |
|  | 1542633699 | |
|  | 1543552144 | |
|  | 1545387776 | |
|  | 1547879760 | |
|  | 1549284008 | |
|  | 1558658757 | |
|  | 1559170183 | |
|  | 1561850253 | |
|  | 1564219984 | |
|  | 1568268146 | |
|  | 1568451843 | |
|  | 1568606682 | |
|  | 1569502690 | |
|  | 1570300669 | |
|  | 1570377143 | |
|  | 1570535615 | |
|  | 1572207475 | |
|  | 1573444561 | |
|  | 1573683112 | |
|  | 1574248945 | |
|  | 1578218178 | |
|  | 1581512783 | |
|  | 1581919998 | |
|  | 1582333603 | |
|  | 1582607016 | |
|  | 1582611257 | |
|  | 1583009052 | |
|  | 1583799011 | |
|  | 1584990857 | |
|  | 1586850127 | |
|  | 1588193232 | |
|  | 1599425600 | |
|  | 1602566724 | |
|  | 1602567453 | |
|  | 1602809129 | |
|  | 1603195963 | |
|  | 1604304397 | |
|  | 1611751080 | |
|  | 1613850255 | |
|  | 1614150208 | |
|  | 1615344303 | |
|  | 1616436608 | |
|  | 1617664615 | |
|  | 1618787606 | |
|  | 1620258790 | |
|  | 1620460207 | |
|  | 1621884272 | |
|  | 1622795192 | |
|  | 1624437721 | |
|  | 1626105231 | |
|  | 1628605685 | |
|  | 1632503489 | |
|  | 1634017026 | |
|  | 1634606847 | |
|  | 1636327326 | |
|  | 1637836422 | |
|  | 1638695876 | |
|  | 1638849234 | |
|  | 1640551393 | |
|  | 1642318146 | |
|  | 1646750527 | |
|  | 1657871527 | |
|  | 1664608228 | |
|  | 1665524040 | |
|  | 1665909570 | |
|  | 1667185460 | |
|  | 1669960073 | |
|  | 1671692019 | |
|  | 1673980810 | |
|  | 1674219510 | |
|  | 1674860378 | |
|  | 1675092201 | |
|  | 1680467653 | |
|  | 1683958275 | |
|  | 1684459365 | |
|  | 1684525404 | |
|  | 1685248591 | |
|  | 1686095578 | |
|  | 1693267448 | |
|  | 1693331843 | |
|  | 1696659438 | |
|  | 1697851766 | |
|  | 1700605146 | |
|  | 1702335205 | |
|  | 1703184139 | |
|  | 1705316165 | |
|  | 1705430016 | |
|  | 1710869618 | |
|  | 1712080978 | |
|  | 1713195601 | |
|  | 1713555506 | |
|  | 1714595447 | |
|  | 1715903102 | |
|  | 1719784028 | |
|  | 1721067998 | |
|  | 1721149895 | |
|  | 1721719357 | |
|  | 1723713874 | |
|  | 1725931447 | |
|  | 1729463752 | |
|  | 1730555933 | |
|  | 1732562638 | |
|  | 1732621785 | |
|  | 1732628033 | |
|  | 1733160153 | |
|  | 1735424888 | |
|  | 1735672143 | |
|  | 1736026813 | |
|  | 1738265797 | |
|  | 1740651977 | |
|  | 1740768360 | |
|  | 1741427994 | |
|  | 1742208574 | |
|  | 1742698971 | |
|  | 1746956570 | |
|  | 1749795017 | |
|  | 1749886582 | |
|  | 1754248939 | |
|  | 1754370480 | |
|  | 1754393348 | |
|  | 1755363164 | |
|  | 1755577996 | |
|  | 1759357401 | |
|  | 1762382665 | |
|  | 1768844655 | |
|  | 1774019924 | |
|  | 1774142290 | |
|  | 1775810647 | |
|  | 1782011267 | |
|  | 1785683843 | |
|  | 1786160715 | |
|  | 1787372080 | |
|  | 1788683205 | |
|  | 1790668568 | |
|  | 1791250428 | |
|  | 1791271074 | |
|  | 1792562719 | |
|  | 1794378966 | |
|  | 1794757945 | |
|  | 1795978025 | |
|  | 1805352855 | |
|  | 1807812884 | |
|  | 1814116706 | |
|  | 1816178031 | |
|  | 1819912581 | |
|  | 1822335788 | |
|  | 1824088295 | |
|  | 1825586641 | |
|  | 1830939119 | |
|  | 1832793247 | |
|  | 1839403113 | |
|  | 1840994228 | |
|  | 1842114921 | |
|  | 1842608396 | |
|  | 1842898132 | |
|  | 1842929960 | |
|  | 1845306853 | |
|  | 1851683108 | |
|  | 1854141252 | |
|  | 1856075173 | |
|  | 1857049193 | |
|  | 1857396528 | |
|  | 1858577693 | |
|  | 1860548965 | |
|  | 1861965050 | |
|  | 1865863787 | |
|  | 1868602760 | |
|  | 1870383208 | |
|  | 1871221615 | |
|  | 1871590209 | |
|  | 1876820278 | |
|  | 1888311577 | |
|  | 1890487293 | |
|  | 1891839201 | |
|  | 1893133531 | |
|  | 1894132300 | |
|  | 1896049752 | |
|  | 1900944773 | |
|  | 1901992722 | |
|  | 1902789874 | |
|  | 1903882444 | |
|  | 1906188330 | |
|  | 1906719888 | |
|  | 1909986973 | |
|  | 1910057321 | |
|  | 1911461466 | |
|  | 1919707488 | |
|  | 1922070456 | |
|  | 1923187777 | |
|  | 1923620478 | |
|  | 1924383628 | |
|  | 1926213669 | |
|  | 1926549088 | |
|  | 1929041313 | |
|  | 1931324871 | |
|  | 1933675971 | |
|  | 1935726747 | |
|  | 1936150468 | |
|  | 1937270128 | |
|  | 1937707958 | |
|  | 1941511195 | |
|  | 1942537345 | |
|  | 1945063726 | |
|  | 1945614622 | |
|  | 1954231082 | |
|  | 1955172768 | |
|  | 1958069366 | |
|  | 1961848637 | |
|  | 1962060547 | |
|  | 1963389888 | |
|  | 1974563496 | |
|  | 1976351578 | |
|  | 1978844073 | |
|  | 1979131117 | |
|  | 1979311206 | |
|  | 1979435572 | |
|  | 1981124422 | |
|  | 1985332818 | |
|  | 1989436032 | |
|  | 1991791946 | |
|  | 1994338942 | |
|  | 1995075671 | |
|  | 1997604307 | |
|  | 1999051638 | |
|  | 1999904870 | |
|  | 2000848104 | |
|  | 2002911368 | |
|  | 2004727027 | |
|  | 2009615374 | |
|  | 2010446245 | |
|  | 2013664411 | |
|  | 2014255590 | |
|  | 2014543234 | |
|  | 2017211826 | |
|  | 2017861928 | |
|  | 2021757847 | |
|  | 2021812431 | |
|  | 2022575067 | |
|  | 2025339464 | |
|  | 2027314762 | |
|  | 2030757576 | |
|  | 2031880624 | |
|  | 2032743038 | |
|  | 2034850751 | |
|  | 2034867880 | |
|  | 2035123431 | |
|  | 2035321105 | |
|  | 2036328569 | |
|  | 2038385394 | |
|  | 2041434490 | |
|  | 2052282471 | |
|  | 2053123511 | |
|  | 2053977436 | |
|  | 2054620141 | |
|  | 2059226165 | |
|  | 2059488795 | |
|  | 2060256112 | |
|  | 2060762800 | |
|  | 2063807541 | |
|  | 2064788354 | |
|  | 2067609940 | |
|  | 2070976245 | |
|  | 2072239802 | |
|  | 2076129780 | |
|  | 2076190208 | |
|  | 2082797629 | |
|  | 2083572173 | |
|  | 2083590407 | |
|  | 2084481909 | |
|  | 2085400973 | |
|  | 2089117413 | |
|  | 2090425988 | |
|  | 2092489639 | |
|  | 2094100161 | |
|  | 2097432113 | |
|  | 2098288598 | |
|  | 2099021818 | |
|  | 2099359955 | |
|  | 2100420998 | |
|  | 2104205232 | |
|  | 2104570636 | |
|  | 2106530579 | |
|  | 2107190963 | |
|  | 2110118862 | |
|  | 2110713573 | |
|  | 2113210395 | |
|  | 2115476908 | |
|  | 2117068077 | |
|  | 2117887031 | |
|  | 2119007969 | |
|  | 2119439498 | |
|  | 2122968379 | |
|  | 2125043625 | |
|  | 2125504268 | |
|  | 2125634645 | |
|  | 2126508391 | |
|  | 2128055421 | |
|  | 2129090136 | |
|  | 2131356872 | |
|  | 2132058917 | |
|  | 2132129411 | |
|  | 2132511834 | |
|  | 2132620936 | |
|  | 2140211822 | |
|  | 2140805174 | |
|  | 2142032900 | |
|  | 2142685628 | |
|  | 2143772823 | |
|  | 2144040306 | |
|  | 2146359984 | |
|  | 2147279852 | |
|  | 2152210560 | |
|  | 2154777022 | |
|  | 2154935424 | |
|  | 2154975788 | |
|  | 2158012165 | |
|  | 2159088240 | |
|  | 2160246537 | |
|  | 2160502559 | |
|  | 2160803309 | |
|  | 2161844995 | |
|  | 2171491174 | |
|  | 2172997423 | |
|  | 2173851476 | |
|  | 2179518760 | |
|  | 2184430117 | |
|  | 2185212638 | |
|  | 2188215619 | |
|  | 2189758440 | |
|  | 2192183111 | |
|  | 2194376677 | |
|  | 2196690495 | |
|  | 2201149108 | |
|  | 2206026610 | |
|  | 2206529641 | |
|  | 2211217220 | |
|  | 2211527187 | |
|  | 2211843251 | |
|  | 2211983264 | |
|  | 2212573984 | |
|  | 2214592779 | |
|  | 2215059400 | |
|  | 2215423748 | |
|  | 2218281189 | |
|  | 2222621677 | |
|  | 2222715027 | |
|  | 2223687999 | |
|  | 2226535532 | |
|  | 2228063684 | |
|  | 2228215562 | |
|  | 2228262129 | |
|  | 2228558432 | |
|  | 2232363332 | |
|  | 2233928532 | |
|  | 2237617467 | |
|  | 2238101436 | |
|  | 2240437263 | |
|  | 2244444028 | |
|  | 2244797474 | |
|  | 2245273601 | |
|  | 2245277810 | |
|  | 2245384272 | |
|  | 2245900962 | |
|  | 2246340824 | |
|  | 2246699815 | |
|  | 2246703798 | |
|  | 2246728737 | |
|  | 2246997334 | |
|  | 2249497313 | |
|  | 2250750806 | |
|  | 2251845666 | |
|  | 2251883808 | |
|  | 2251888095 | |
|  | 2252049393 | |
|  | 2257970297 | |
|  | 2258843522 | |
|  | 2260741221 | |
|  | 2261230370 | |
|  | 2262965892 | |
|  | 2264628208 | |
|  | 2264700157 | |
|  | 2269518716 | |
|  | 2273642216 | |
|  | 2275683444 | |
|  | 2276548146 | |
|  | 2280278597 | |
|  | 2281561161 | |
|  | 2284773903 | |
|  | 2285421074 | |
|  | 2290389376 | |
|  | 2295262118 | |
|  | 2295519362 | |
|  | 2295747111 | |
|  | 2296493092 | |
|  | 2296565457 | |
|  | 2297887526 | |
|  | 2299023194 | |
|  | 2299784278 | |
|  | 2302345434 | |
|  | 2304698736 | |
|  | 2305802494 | |
|  | 2308348490 | |
|  | 2309124039 | |
|  | 2310216636 | |
|  | 2310666682 | |
|  | 2315898571 | |
|  | 2319429949 | |
|  | 2322101270 | |
|  | 2324641173 | |
|  | 2325814211 | |
|  | 2329247646 | |
|  | 2331740049 | |
|  | 2333272823 | |
|  | 2334784814 | |
|  | 2335258807 | |
|  | 2338065186 | |
|  | 2341250177 | |
|  | 2341340169 | |
|  | 2341585586 | |
|  | 2342410226 | |
|  | 2344409743 | |
|  | 2345191348 | |
|  | 2347776210 | |
|  | 2349397595 | |
|  | 2352465170 | |
|  | 2353112200 | |
|  | 2353583745 | |
|  | 2355616982 | |
|  | 2360741695 | |
|  | 2363984686 | |
|  | 2368838505 | |
|  | 2368852967 | |
|  | 2369304345 | |
|  | 2370996728 | |
|  | 2373710881 | |
|  | 2373891708 | |
|  | 2374957460 | |
|  | 2378775366 | |
|  | 2378779377 | |
|  | 2380929092 | |
|  | 2390579542 | |
|  | 2392057161 | |
|  | 2394178657 | |
|  | 2399075688 | |
|  | 2399596470 | |
|  | 2401082571 | |
|  | 2401181567 | |
|  | 2402406502 | |
|  | 2409787606 | |
|  | 2411493228 | |
|  | 2414650566 | |
|  | 2415288965 | |
|  | 2418375248 | |
|  | 2420735149 | |
|  | 2422851564 | |
|  | 2423543607 | |
|  | 2424530656 | |
|  | 2424973678 | |
|  | 2425865203 | |
|  | 2428426313 | |
|  | 2428614265 | |
|  | 2429890643 | |
|  | 2431669681 | |
|  | 2434476610 | |
|  | 2434661383 | |
|  | 2435680164 | |
|  | 2439001532 | |
|  | 2441429409 | |
|  | 2442359259 | |
|  | 2442900373 | |
|  | 2443462774 | |
|  | 2443676976 | |
|  | 2445100084 | |
|  | 2446961558 | |
|  | 2447748155 | |
|  | 2448462321 | |
|  | 2449545466 | |
|  | 2450291699 | |
|  | 2451306927 | |
|  | 2451661632 | |
|  | 2452462412 | |
|  | 2455017113 | |
|  | 2455275232 | |
|  | 2457874744 | |
|  | 2458968089 | |
|  | 2459558207 | |
|  | 2459912894 | |
|  | 2463614838 | |
|  | 2473389857 | |
|  | 2473818866 | |
|  | 2479916646 | |
|  | 2482040061 | |
|  | 2486053381 | |
|  | 2487927669 | |
|  | 2490400667 | |
|  | 2491708295 | |
|  | 2494804746 | |
|  | 2495104816 | |
|  | 2502538098 | |
|  | 2504604572 | |
|  | 2505330992 | |
|  | 2506328221 | |
|  | 2506939770 | |
|  | 2509211297 | |
|  | 2509875913 | |
|  | 2510215966 | |
|  | 2511318058 | |
|  | 2513428378 | |
|  | 2514702557 | |
|  | 2515833467 | |
|  | 2516079810 | |
|  | 2516197204 | |
|  | 2516309301 | |
|  | 2524292098 | |
|  | 2524325328 | |
|  | 2524958857 | |
|  | 2525709234 | |
|  | 2527619210 | |
|  | 2529257294 | |
|  | 2529809651 | |
|  | 2530377886 | |
|  | 2531703152 | |
|  | 2535347081 | |
|  | 2536602154 | |
|  | 2537263309 | |
|  | 2542488922 | |
|  | 2543587995 | |
|  | 2543874606 | |
|  | 2549196227 | |
|  | 2551483158 | |
|  | 2552737839 | |
|  | 2557045420 | |
|  | 2558454195 | |
|  | 2558854650 | |
|  | 2559675920 | |
|  | 2560252747 | |
|  | 2560867404 | |
|  | 2562008222 | |
|  | 2564760642 | |
|  | 2564933757 | |
|  | 2565499684 | |
|  | 2565679125 | |
|  | 2567005256 | |
|  | 2567905687 | |
|  | 2569072435 | |
|  | 2572035617 | |
|  | 2572949265 | |
|  | 2574787838 | |
|  | 2575073586 | |
|  | 2575091433 | |
|  | 2575624905 | |
|  | 2577837950 | |
|  | 2579153554 | |
|  | 2582570729 | |
|  | 2583926374 | |
|  | 2584776234 | |
|  | 2585021504 | |
|  | 2591432844 | |
|  | 2592252298 | |
|  | 2592785365 | |
|  | 2592975058 | |
|  | 2594220197 | |
|  | 2595731191 | |
|  | 2596116113 | |
|  | 2598322190 | |
|  | 2602464909 | |
|  | 2603759688 | |
|  | 2603892347 | |
|  | 2604440622 | |
|  | 2607036443 | |
|  | 2607304338 | |
|  | 2609658248 | |
|  | 2610761730 | |
|  | 2613148343 | |
|  | 2614860224 | |
|  | 2617658656 | |
|  | 2621717915 | |
|  | 2622370625 | |
|  | 2626152126 | |
|  | 2627504773 | |
|  | 2629657136 | |
|  | 2629723425 | |
|  | 2629928014 | |
|  | 2635303940 | |
|  | 2636383078 | |
|  | 2637439965 | |
|  | 2637535902 | |
|  | 2640652321 | |
|  | 2640800466 | |
|  | 2644702627 | |
|  | 2644797347 | |
|  | 2645514824 | |
|  | 2646219661 | |
|  | 2649212702 | |
|  | 2649902721 | |
|  | 2650167165 | |
|  | 2651067438 | |
|  | 2664995851 | |
|  | 2667063169 | |
|  | 2667430530 | |
|  | 2668155513 | |
|  | 2668267711 | |
|  | 2668574517 | |
|  | 2669055056 | |
|  | 2669891477 | |
|  | 2672648630 | |
|  | 2673417767 | |
|  | 2673639235 | |
|  | 2674474075 | |
|  | 2674622095 | |
|  | 2677858541 | |
|  | 2687950697 | |
|  | 2688692504 | |
|  | 2691656950 | |
|  | 2693939312 | |
|  | 2697110228 | |
|  | 2697642734 | |
|  | 2698428365 | |
|  | 2700220651 | |
|  | 2705805446 | |
|  | 2706078914 | |
|  | 2712470536 | |
|  | 2713070966 | |
|  | 2719762529 | |
|  | 2720313463 | |
|  | 2724477936 | |
|  | 2726586130 | |
|  | 2726983370 | |
|  | 2728125556 | |
|  | 2729097954 | |
|  | 2729628705 | |
|  | 2730755516 | |
|  | 2731256589 | |
|  | 2731613698 | |
|  | 2731746285 | |
|  | 2739425568 | |
|  | 2741574634 | |
|  | 2744158313 | |
|  | 2746488638 | |
|  | 2746975098 | |
|  | 2749896868 | |
|  | 2751466952 | |
|  | 2752034647 | |
|  | 2752308248 | |
|  | 2756611893 | |
|  | 2757088131 | |
|  | 2759434019 | |
|  | 2760457081 | |
|  | 2762756625 | |
|  | 2762810942 | |
|  | 2763854213 | |
|  | 2766434492 | |
|  | 2768992039 | |
|  | 2771442675 | |
|  | 2776365053 | |
|  | 2777262656 | |
|  | 2782530898 | |
|  | 2782747791 | |
|  | 2783552463 | |
|  | 2784506312 | |
|  | 2786465058 | |
|  | 2787134657 | |
|  | 2787503806 | |
|  | 2789648773 | |
|  | 2791547925 | |
|  | 2793027215 | |
|  | 2803419857 | |
|  | 2803848648 | |
|  | 2804184868 | |
|  | 2804524294 | |
|  | 2806018737 | |
|  | 2807496773 | |
|  | 2809361097 | |
|  | 2810604122 | |
|  | 2811394787 | |
|  | 2812398603 | |
|  | 2814583100 | |
|  | 2817491392 | |
|  | 2819034113 | |
|  | 2825490293 | |
|  | 2826135114 | |
|  | 2827868305 | |
|  | 2829279174 | |
|  | 2832976762 | |
|  | 2833025332 | |
|  | 2834566420 | |
|  | 2835079913 | |
|  | 2836150064 | |
|  | 2839110432 | |
|  | 2841393247 | |
|  | 2843706522 | |
|  | 2843970853 | |
|  | 2844835166 | |
|  | 2845281347 | |
|  | 2847919965 | |
|  | 2848578460 | |
|  | 2849741637 | |
|  | 2850370298 | |
|  | 2850656190 | |
|  | 2856245513 | |
|  | 2857220865 | |
|  | 2859975680 | |
|  | 2863953098 | |
|  | 2864705169 | |
|  | 2867364119 | |
|  | 2869481095 | |
|  | 2871087835 | |
|  | 2873206065 | |
|  | 2875639393 | |
|  | 2877223105 | |
|  | 2878620822 | |
|  | 2885491308 | |
|  | 2889160788 | |
|  | 2890522819 | |
|  | 2892360967 | |
|  | 2892519151 | |
|  | 2892637219 | |
|  | 2895288982 | |
|  | 2895999632 | |
|  | 2896269280 | |
|  | 2898623098 | |
|  | 2899053136 | |
|  | 2903607685 | |
|  | 2904824113 | |
|  | 2904998726 | |
|  | 2905660137 | |
|  | 2905822145 | |
|  | 2907250678 | |
|  | 2908279847 | |
|  | 2909636892 | |
|  | 2910395211 | |
|  | 2912042324 | |
|  | 2912830040 | |
|  | 2914602668 | |
|  | 2915034670 | |
|  | 2922016024 | |
|  | 2922136249 | |
|  | 2923200147 | |
|  | 2924120306 | |
|  | 2929617213 | |
|  | 2931695851 | |
|  | 2931878080 | |
|  | 2932724336 | |
|  | 2933418651 | |
|  | 2936019033 | |
|  | 2939120473 | |
|  | 2944555726 | |
|  | 2946106206 | |
|  | 2950679810 | |
|  | 2951428091 | |
|  | 2952211277 | |
|  | 2952902656 | |
|  | 2954329948 | |
|  | 2956273779 | |
|  | 2958413073 | |
|  | 2959332130 | |
|  | 2959422742 | |
|  | 2963238476 | |
|  | 2965330009 | |
|  | 2967245471 | |
|  | 2968303586 | |
|  | 2968820007 | |
|  | 2968968094 | |
|  | 2969511302 | |
|  | 2971572579 | |
|  | 2972062794 | |
|  | 2972599459 | |
|  | 2974534217 | |
|  | 2975126068 | |
|  | 2975700596 | |
|  | 2976033787 | |
|  | 2976816164 | |
|  | 2977973464 | |
|  | 2978961685 | |
|  | 2979336775 | |
|  | 2979609357 | |
|  | 2981620181 | |
|  | 2985339471 | |
|  | 2986138612 | |
|  | 2987120039 | |
|  | 2989172416 | |
|  | 2989738444 | |
|  | 2992008339 | |
|  | 2994748777 | |
|  | 2995598328 | |
|  | 2997633404 | |
|  | 2999514534 | |
|  | 3000202447 | |
|  | 3002759401 | |
|  | 3002887899 | |
|  | 3004333805 | |
|  | 3005152558 | |
|  | 3005189210 | |
|  | 3008098818 | |
|  | 3008585314 | |
|  | 3009478870 | |
|  | 3010410747 | |
|  | 3011598321 | |
|  | 3011650553 | |
|  | 3012204942 | |
|  | 3012865236 | |
|  | 3022163420 | |
|  | 3023959105 | |
|  | 3025629386 | |
|  | 3026237809 | |
|  | 3026394695 | |
|  | 3027067501 | |
|  | 3028039221 | |
|  | 3039572119 | |
|  | 3042542132 | |
|  | 3042770624 | |
|  | 3044392905 | |
|  | 3044751281 | |
|  | 3046825176 | |
|  | 3051562454 | |
|  | 3058733262 | |
|  | 3059129780 | |
|  | 3065279093 | |
|  | 3068057572 | |
|  | 3074137104 | |
|  | 3076806434 | |
|  | 3084241488 | |
|  | 3084500287 | |
|  | 3087437574 | |
|  | 3088749993 | |
|  | 3088822697 | |
|  | 3090652201 | |
|  | 3095161989 | |
|  | 3099084201 | |
|  | 3099386124 | |
|  | 3103812183 | |
|  | 3106246680 | |
|  | 3106764902 | |
|  | 3109601040 | |
|  | 3111861837 | |
|  | 3111864946 | |
|  | 3113855253 | |
|  | 3116051204 | |
|  | 3118255683 | |
|  | 3119383463 | |
|  | 3120642300 | |
|  | 3120734294 | |
|  | 3120784405 | |
|  | 3121777292 | |
|  | 3124390475 | |
|  | 3124581743 | |
|  | 3129246025 | |
|  | 3129492592 | |
|  | 3129571007 | |
|  | 3131105066 | |
|  | 3134313966 | |
|  | 3135560665 | |
|  | 3137088118 | |
|  | 3142691941 | |
|  | 3143719699 | |
|  | 3146955833 | |
|  | 3147339068 | |
|  | 3148386287 | |
|  | 3149717256 | |
|  | 3149867162 | |
|  | 3152170373 | |
|  | 3156961613 | |
|  | 3157481675 | |
|  | 3157855630 | |
|  | 3158187473 | |
|  | 3160601726 | |
|  | 3162837314 | |
|  | 3163510888 | |
|  | 3163669616 | |
|  | 3164804235 | |
|  | 3171733977 | |
|  | 3175776621 | |
|  | 3176649511 | |
|  | 3176806076 | |
|  | 3177290410 | |
|  | 3177362943 | |
|  | 3179791401 | |
|  | 3180000854 | |
|  | 3181405140 | |
|  | 3181446473 | |
|  | 3182700296 | |
|  | 3182824521 | |
|  | 3183067622 | |
|  | 3183395256 | |
|  | 3187793195 | |
|  | 3189457552 | |
|  | 3190668333 | |
|  | 3192617127 | |
|  | 3194351646 | |
|  | 3194612514 | |
|  | 3195927545 | |
|  | 3198674699 | |
|  | 3207567135 | |
|  | 3212498122 | |
|  | 3216224414 | |
|  | 3217380708 | |
|  | 3218262540 | |
|  | 3218693969 | |
|  | 3221148524 | |
|  | 3224978178 | |
|  | 3228732977 | |
|  | 3228757619 | |
|  | 3230131948 | |
|  | 3234023386 | |
|  | 3234104871 | |
|  | 3244406875 | |
|  | 3246071060 | |
|  | 3248361648 | |
|  | 3249313900 | |
|  | 3253842763 | |
|  | 3254172035 | |
|  | 3255046070 | |
|  | 3255377480 | |
|  | 3260793816 | |
|  | 3260853321 | |
|  | 3261096889 | |
|  | 3261293650 | |
|  | 3262126146 | |
|  | 3263463138 | |
|  | 3265203944 | |
|  | 3266039179 | |
|  | 3266686894 | |
|  | 3267049022 | |
|  | 3268335377 | |
|  | 3269035194 | |
|  | 3269951047 | |
|  | 3271309638 | |
|  | 3272800686 | |
|  | 3273018953 | |
|  | 3275683399 | |
|  | 3277944515 | |
|  | 3278587996 | |
|  | 3279739845 | |
|  | 3285151283 | |
|  | 3286720230 | |
|  | 3288032115 | |
|  | 3290988458 | |
|  | 3291149959 | |
|  | 3292496536 | |
|  | 3308295291 | |
|  | 3309718011 | |
|  | 3311784590 | |
|  | 3311951855 | |
|  | 3313841562 | |
|  | 3315826729 | |
|  | 3320432666 | |
|  | 3321054387 | |
|  | 3324153899 | |
|  | 3324518393 | |
|  | 3324835923 | |
|  | 3324998366 | |
|  | 3327598791 | |
|  | 3327625720 | |
|  | 3327911390 | |
|  | 3328145258 | |
|  | 3328900012 | |
|  | 3329119289 | |
|  | 3332019264 | |
|  | 3332711906 | |
|  | 3334525955 | |
|  | 3335427733 | |
|  | 3337420400 | |
|  | 3337745083 | |
|  | 3340462994 | |
|  | 3341916169 | |
|  | 3342901245 | |
|  | 3345225135 | |
|  | 3346539298 | |
|  | 3346582092 | |
|  | 3351556771 | |
|  | 3352490390 | |
|  | 3354929391 | |
|  | 3358702130 | |
|  | 3358736304 | |
|  | 3369800066 | |
|  | 3372972590 | |
|  | 3375560558 | |
|  | 3376021752 | |
|  | 3377254929 | |
|  | 3377547859 | |
|  | 3380438856 | |
|  | 3381412387 | |
|  | 3381665419 | |
|  | 3381668292 | |
|  | 3386795352 | |
|  | 3393100573 | |
|  | 3394927196 | |
|  | 3396393889 | |
|  | 3399951910 | |
|  | 3399962845 | |
|  | 3400020746 | |
|  | 3400645812 | |
|  | 3402594617 | |
|  | 3407523657 | |
|  | 3408765322 | |
|  | 3412210860 | |
|  | 3412947479 | |
|  | 3413384544 | |
|  | 3415464045 | |
|  | 3417020623 | |
|  | 3417857634 | |
|  | 3421227752 | |
|  | 3424025209 | |
|  | 3429697732 | |
|  | 3430370997 | |
|  | 3430575859 | |
|  | 3430642422 | |
|  | 3432086929 | |
|  | 3434359339 | |
|  | 3435437068 | |
|  | 3442795883 | |
|  | 3443041413 | |
|  | 3448041912 | |
|  | 3450512647 | |
|  | 3452424498 | |
|  | 3452535345 | |
|  | 3455614769 | |
|  | 3458774156 | |
|  | 3460898310 | |
|  | 3461773897 | |
|  | 3462333187 | |
|  | 3462479532 | |
|  | 3466404006 | |
|  | 3472639681 | |
|  | 3479423841 | |
|  | 3480856402 | |
|  | 3481960194 | |
|  | 3482873808 | |
|  | 3488664301 | |
|  | 3488928039 | |
|  | 3488951756 | |
|  | 3491109101 | |
|  | 3491543932 | |
|  | 3493793229 | |
|  | 3494024880 | |
|  | 3502700335 | |
|  | 3503842959 | |
|  | 3508865297 | |
|  | 3520289908 | |
|  | 3524145956 | |
|  | 3525819727 | |
|  | 3527448998 | |
|  | 3528907692 | |
|  | 3531241045 | |
|  | 3531536209 | |
|  | 3532053313 | |
|  | 3532733700 | |
|  | 3535578123 | |
|  | 3537119515 | |
|  | 3537123720 | |
|  | 3538861869 | |
|  | 3540287614 | |
|  | 3540401924 | |
|  | 3542045346 | |
|  | 3542456614 | |
|  | 3545353036 | |
|  | 3545365497 | |
|  | 3550478926 | |
|  | 3553818046 | |
|  | 3556587083 | |
|  | 3557152699 | |
|  | 3557699678 | |
|  | 3559852389 | |
|  | 3561006593 | |
|  | 3561946030 | |
|  | 3563227052 | |
|  | 3565565663 | |
|  | 3573166628 | |
|  | 3573188050 | |
|  | 3579962709 | |
|  | 3581812808 | |
|  | 3585531950 | |
|  | 3591356197 | |
|  | 3593041223 | |
|  | 3594356142 | |
|  | 3594646603 | |
|  | 3594696610 | |
|  | 3596485316 | |
|  | 3599133573 | |
|  | 3599391724 | |
|  | 3601928391 | |
|  | 3607319266 | |
|  | 3609483052 | |
|  | 3609735552 | |
|  | 3612926680 | |
|  | 3613617905 | |
|  | 3613721458 | |
|  | 3613916939 | |
|  | 3616878512 | |
|  | 3618831505 | |
|  | 3621923070 | |
|  | 3623063705 | |
|  | 3626680192 | |
|  | 3627524306 | |
|  | 3631761933 | |
|  | 3632350815 | |
|  | 3641117322 | |
|  | 3642583640 | |
|  | 3643491447 | |
|  | 3644941081 | |
|  | 3646640021 | |
|  | 3647526986 | |
|  | 3647731884 | |
|  | 3648820092 | |
|  | 3649454404 | |
|  | 3651174070 | |
|  | 3653560676 | |
|  | 3657328276 | |
|  | 3657471097 | |
|  | 3659125690 | |
|  | 3659806237 | |
|  | 3661262542 | |
|  | 3665875809 | |
|  | 3666567924 | |
|  | 3670614808 | |
|  | 3673736185 | |
|  | 3675674571 | |
|  | 3675951595 | |
|  | 3677836265 | |
|  | 3678650707 | |
|  | 3681261157 | |
|  | 3684013815 | |
|  | 3685257385 | |
|  | 3685328280 | |
|  | 3689791161 | |
|  | 3692055567 | |
|  | 3692176620 | |
|  | 3692680800 | |
|  | 3694199440 | |
|  | 3696402029 | |
|  | 3696580341 | |
|  | 3698257053 | |
|  | 3699369810 | |
|  | 3701504231 | |
|  | 3702168190 | |
|  | 3703934905 | |
|  | 3704548532 | |
|  | 3705139132 | |
|  | 3706229233 | |
|  | 3712137798 | |
|  | 3718064757 | |
|  | 3718757751 | |
|  | 3721278156 | |
|  | 3722806490 | |
|  | 3727977332 | |
|  | 3730894340 | |
|  | 3731979411 | |
|  | 3735247081 | |
|  | 3742418174 | |
|  | 3745247955 | |
|  | 3745470532 | |
|  | 3745584548 | |
|  | 3747360616 | |
|  | 3749319234 | |
|  | 3749587434 | |
|  | 3751064229 | |
|  | 3753068593 | |
|  | 3755128710 | |
|  | 3757038672 | |
|  | 3759586448 | |
|  | 3760180875 | |
|  | 3763383337 | |
|  | 3770784874 | |
|  | 3772302167 | |
|  | 3775468872 | |
|  | 3776148706 | |
|  | 3776905034 | |
|  | 3777168895 | |
|  | 3777243454 | |
|  | 3777584818 | |
|  | 3779753094 | |
|  | 3781885420 | |
|  | 3785696842 | |
|  | 3787488754 | |
|  | 3793196542 | |
|  | 3793515369 | |
|  | 3796359118 | |
|  | 3796841743 | |
|  | 3796970912 | |
|  | 3798139026 | |
|  | 3801831583 | |
|  | 3806389659 | |
|  | 3810633824 | |
|  | 3811375594 | |
|  | 3816921572 | |
|  | 3818546315 | |
|  | 3820070372 | |
|  | 3820826669 | |
|  | 3824063894 | |
|  | 3825284333 | |
|  | 3833071061 | |
|  | 3833608434 | |
|  | 3835273908 | |
|  | 3836081194 | |
|  | 3836607521 | |
|  | 3840147540 | |
|  | 3840182861 | |
|  | 3843666872 | |
|  | 3847232715 | |
|  | 3847608821 | |
|  | 3848368466 | |
|  | 3848534832 | |
|  | 3848614249 | |
|  | 3850856377 | |
|  | 3852169106 | |
|  | 3853122790 | |
|  | 3853295885 | |
|  | 3855048697 | |
|  | 3857712607 | |
|  | 3859681392 | |
|  | 3866935459 | |
|  | 3866957768 | |
|  | 3868182870 | |
|  | 3869579337 | |
|  | 3869852276 | |
|  | 3870683401 | |
|  | 3872837863 | |
|  | 3873306213 | |
|  | 3873322538 | |
|  | 3876289568 | |
|  | 3877806992 | |
|  | 3880022399 | |
|  | 3884031532 | |
|  | 3884454603 | |
|  | 3886201961 | |
|  | 3887717254 | |
|  | 3888537269 | |
|  | 3889565720 | |
|  | 3889738582 | |
|  | 3892941493 | |
|  | 3893640578 | |
|  | 3893870893 | |
|  | 3894273779 | |
|  | 3895633315 | |
|  | 3898652266 | |
|  | 3904721945 | |
|  | 3907878801 | |
|  | 3908647167 | |
|  | 3909974435 | |
|  | 3910725794 | |
|  | 3914946506 | |
|  | 3916124310 | |
|  | 3918336191 | |
|  | 3920365076 | |
|  | 3920625465 | |
|  | 3920757710 | |
|  | 3924990413 | |
|  | 3925172229 | |
|  | 3929393147 | |
|  | 3930312868 | |
|  | 3931049057 | |
|  | 3931877381 | |
|  | 3932135717 | |
|  | 3933537673 | |
|  | 3934886170 | |
|  | 3935018034 | |
|  | 3935867650 | |
|  | 3941024655 | |
|  | 3941342539 | |
|  | 3943444448 | |
|  | 3943465899 | |
|  | 3946510714 | |
|  | 3956360032 | |
|  | 3958495319 | |
|  | 3959936273 | |
|  | 3962114891 | |
|  | 3964189818 | |
|  | 3964845468 | |
|  | 3965194470 | |
|  | 3965767626 | |
|  | 3966612213 | |
|  | 3973543784 | |
|  | 3974650111 | |
|  | 3975275337 | |
|  | 3975295864 | |
|  | 3976623167 | |
|  | 3979703018 | |
|  | 3980805843 | |
|  | 3980948056 | |
|  | 3981688378 | |
|  | 3981762496 | |
|  | 3982076256 | |
|  | 3982379045 | |
|  | 3983062349 | |
|  | 3983190549 | |
|  | 3983669392 | |
|  | 3986688618 | |
|  | 3987247535 | |
|  | 3988343070 | |
|  | 3988861584 | |
|  | 3991719838 | |
|  | 3992087745 | |
|  | 3992410619 | |
|  | 3992474495 | |
|  | 3992800027 | |
|  | 3993319192 | |
|  | 3993796134 | |
|  | 3994088662 | |
|  | 3995043796 | |
|  | 3995362948 | |
|  | 3997240246 | |
|  | 3997483227 | |
|  | 3999906991 | |
|  | 4001028360 | |
|  | 4001515211 | |
|  | 4002801397 | |
|  | 4003049590 | |
|  | 4003054055 | |
|  | 4006278292 | |
|  | 4006557530 | |
|  | 4006968963 | |
|  | 4007822218 | |
|  | 4008337421 | |
|  | 4009983216 | |
|  | 4011293459 | |
|  | 4011460791 | |
|  | 4011603269 | |
|  | 4011776249 | |
|  | 4012121575 | |
|  | 4014642748 | |
|  | 4017958407 | |
|  | 4019034699 | |
|  | 4022103728 | |
|  | 4022716898 | |
|  | 4023654873 | |
|  | 4024144969 | |
|  | 4026101393 | |
|  | 4026625396 | |
|  | 4028939580 | |
|  | 4029716266 | |
|  | 4033380444 | |
|  | 4033841756 | |
|  | 4036277955 | |
|  | 4036774035 | |
|  | 4038637638 | |
|  | 4038740760 | |
|  | 4039085716 | |
|  | 4041573576 | |
|  | 4042373501 | |
|  | 4047334924 | |
|  | 4048747042 | |
|  | 4048813351 | |
|  | 4048848834 | |
|  | 4049095736 | |
|  | 4051518313 | |
|  | 4051585127 | |
|  | 4051720567 | |
|  | 4052636517 | |
|  | 4055698890 | |
|  | 4057700103 | |
|  | 4058563477 | |
|  | 4059302282 | |
|  | 4068239769 | |
|  | 4070786387 | |
|  | 4072159216 | |
|  | 4073938295 | |
|  | 4079389161 | |
|  | 4079699505 | |
|  | 4081436575 | |
|  | 4083722611 | |
|  | 4085949867 | |
|  | 4086265842 | |
|  | 4086696725 | |
|  | 4089138501 | |
|  | 4089587568 | |
|  | 4091723021 | |
|  | 4096713990 | |
|  | 4097902480 | |
|  | 4103497225 | |
|  | 4104822550 | |
|  | 4105666400 | |
|  | 4107532493 | |
|  | 4111903471 | |
|  | 4111957990 | |
|  | 4114110979 | |
|  | 4115839854 | |
|  | 4118787735 | |
|  | 4121384005 | |
|  | 4121755354 | |
|  | 4122381561 | |
|  | 4122971576 | |
|  | 4123143112 | |
|  | 4126242811 | |
|  | 4127728637 | |
|  | 4128659070 | |
|  | 4130057166 | |
|  | 4130182993 | |
|  | 4130205841 | |
|  | 4130863294 | |
|  | 4135058020 | |
|  | 4138040424 | |
|  | 4138330676 | |
|  | 4139827362 | |
|  | 4140034840 | |
|  | 4140340356 | |
|  | 4142616092 | |
|  | 4146681879 | |
|  | 4146741105 | |
|  | 4148053715 | |
|  | 4150358042 | |
|  | 4156406673 | |
|  | 4158944142 | |
|  | 4159583953 | |
|  | 4160330003 | |
|  | 4162388837 | |
|  | 4162508222 | |
|  | 4162551380 | |
|  | 4162818269 | |
|  | 4164236649 | |
|  | 4165661399 | |
|  | 4176971489 | |
|  | 4178682433 | |
|  | 4181168628 | |
|  | 4182960975 | |
|  | 4185026439 | |
|  | 4187426726 | |
|  | 4188525416 | |
|  | 4190889139 | |
|  | 4192150624 | |
|  | 4194366826 | |
|  | 4194776273 | |
|  | 4197146180 | |
|  | 4197577604 | |
|  | 4200113039 | |
|  | 4203454020 | |
|  | 4204573730 | |
|  | 4205526060 | |
|  | 4206526563 | |
|  | 4206592788 | |
|  | 4208121018 | |
|  | 4211279910 | |
|  | 4212647803 | |
|  | 4212936508 | |
|  | 4214532707 | |
|  | 4214573929 | |
|  | 4216335232 | |
|  | 4216649732 | |
|  | 4222060125 | |
|  | 4222851645 | |
|  | 4223817698 | |
|  | 4223976160 | |
|  | 4224153346 | |
|  | 4225511501 | |
|  | 4229088015 | |
|  | 4232435237 | |
|  | 4232722574 | |
|  | 4232889011 | |
|  | 4234407427 | |
|  | 4234795114 | |
|  | 4235022035 | |
|  | 4239911899 | |
|  | 4240174685 | |
|  | 4242283111 | |
|  | 4242425044 | |
|  | 4245637465 | |
|  | 4250585145 | |
|  | 4252736910 | |
|  | 4253494739 | |
|  | 4256629345 | |
|  | 4256676941 | |
|  | 4257302536 | |
|  | 4260819931 | |
|  | 4261830945 | |
|  | 4262385000 | |
|  | 4262623969 | |
|  | 4263186572 | |
|  | 4264403900 | |
|  | 4264854005 | |
|  | 4265716399 | |
|  | 4266745719 | |
|  | 4267579308 | |
|  | 4268661680 | |
|  | 4271883048 | |
|  | 4271990713 | |
|  | 4272110579 | |
|  | 4275613329 | |
|  | 4277394373 | |
|  | 4278941385 | |
|  | 4279758435 | |
|  | 4282315164 | |
|  | 4285418163 | |
|  | 4287125113 | |
|  | 4292539202 | |
