## Supplementary figures and images for "Combining Multi-Dimensional Molecular Fingerprints to Predict hERG Cardiotoxicity of Compounds"

### a.bmp

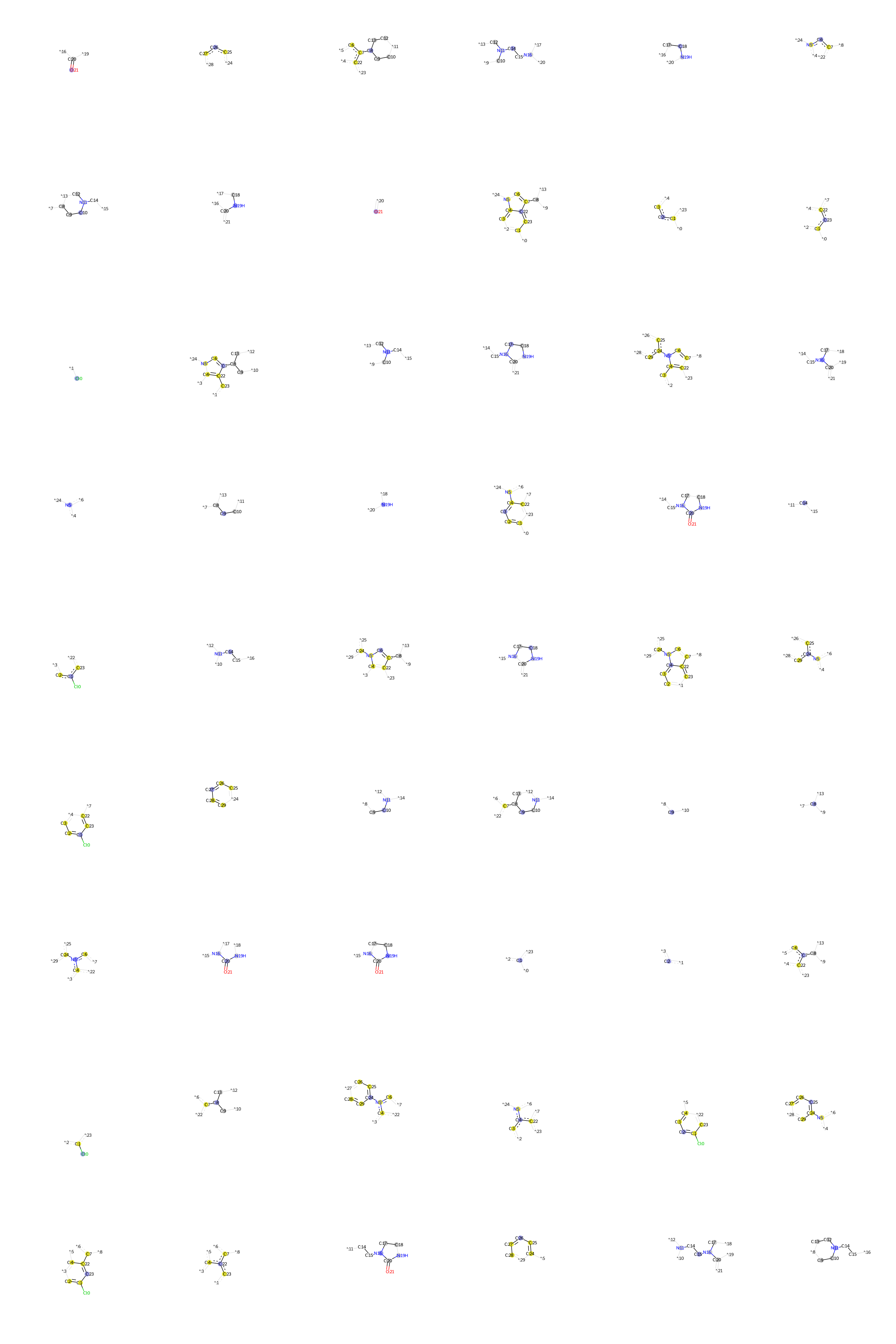

### Ajmaline .bmp

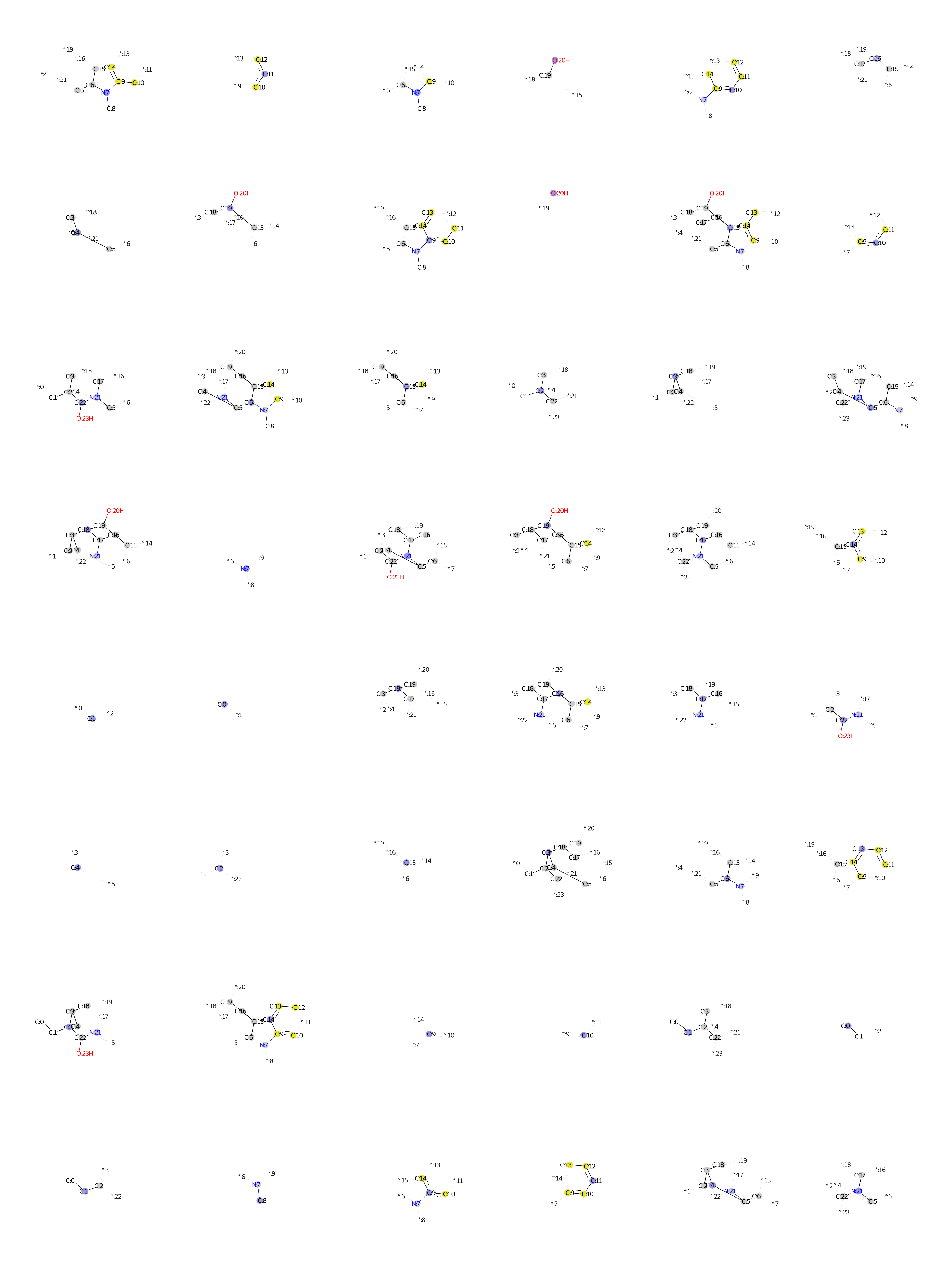

### Alfuzosin.bmp

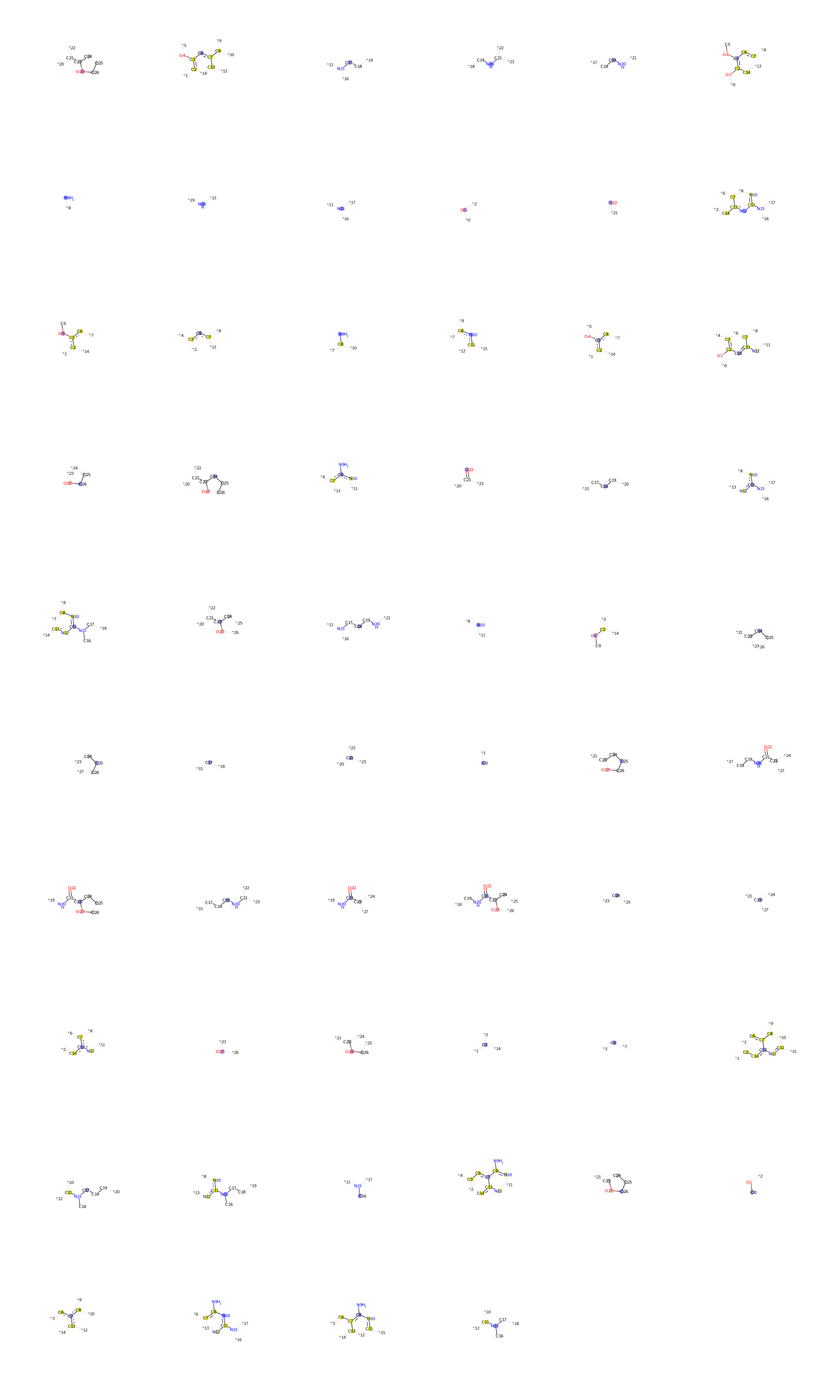

### Alosetron.bmp

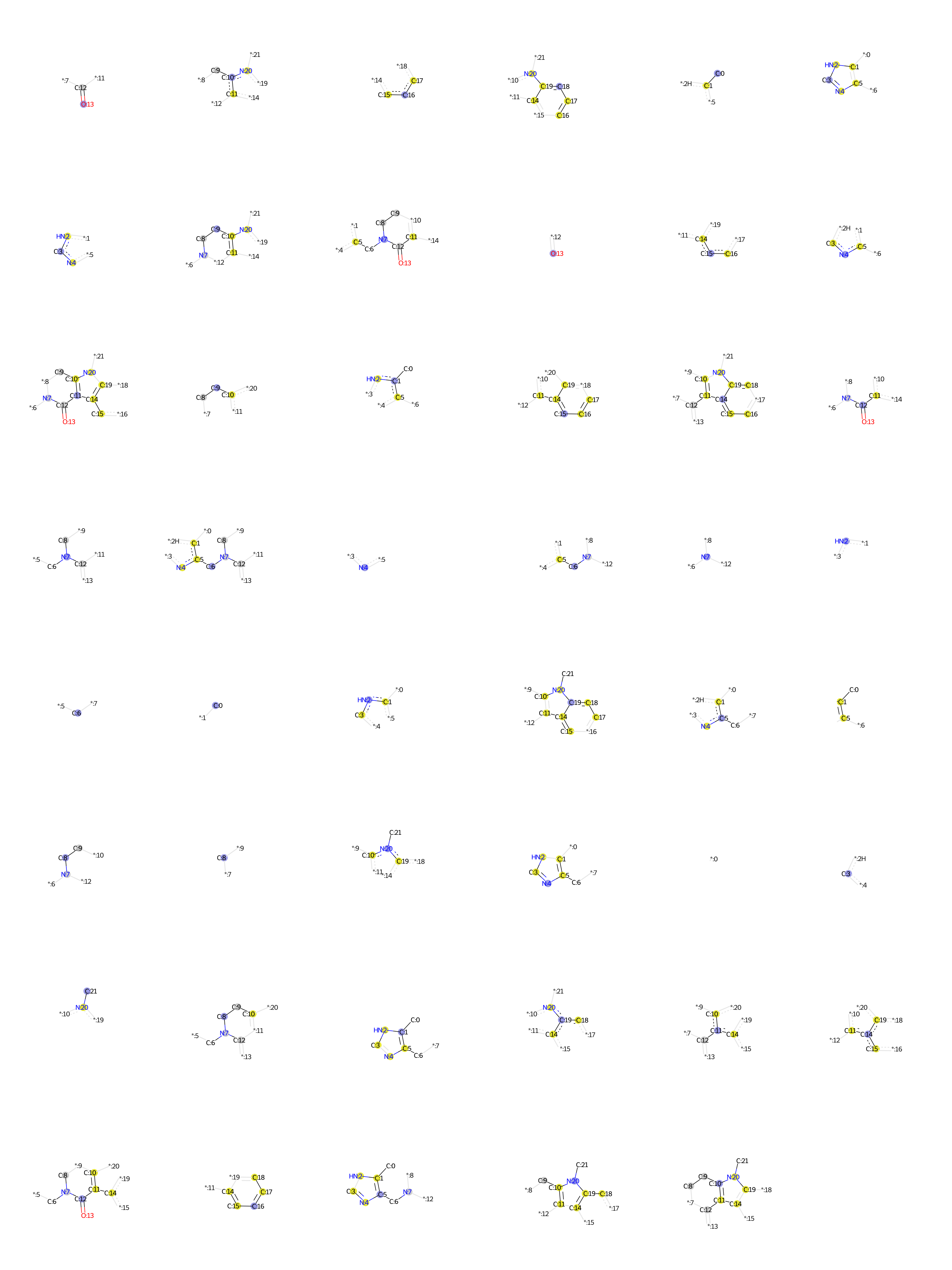

### Alphaacetylmethadol.bmp

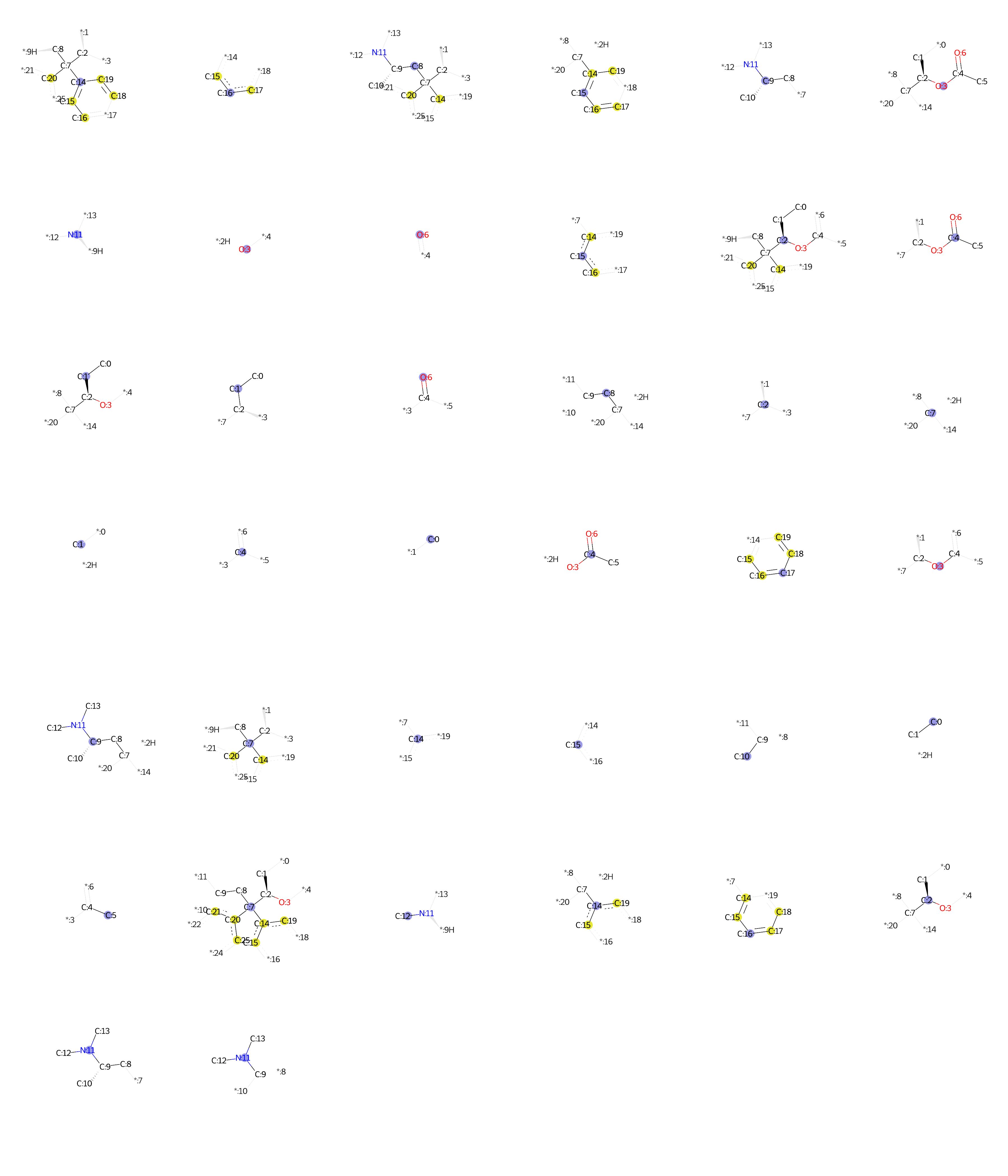

### Ambasilide.bmp

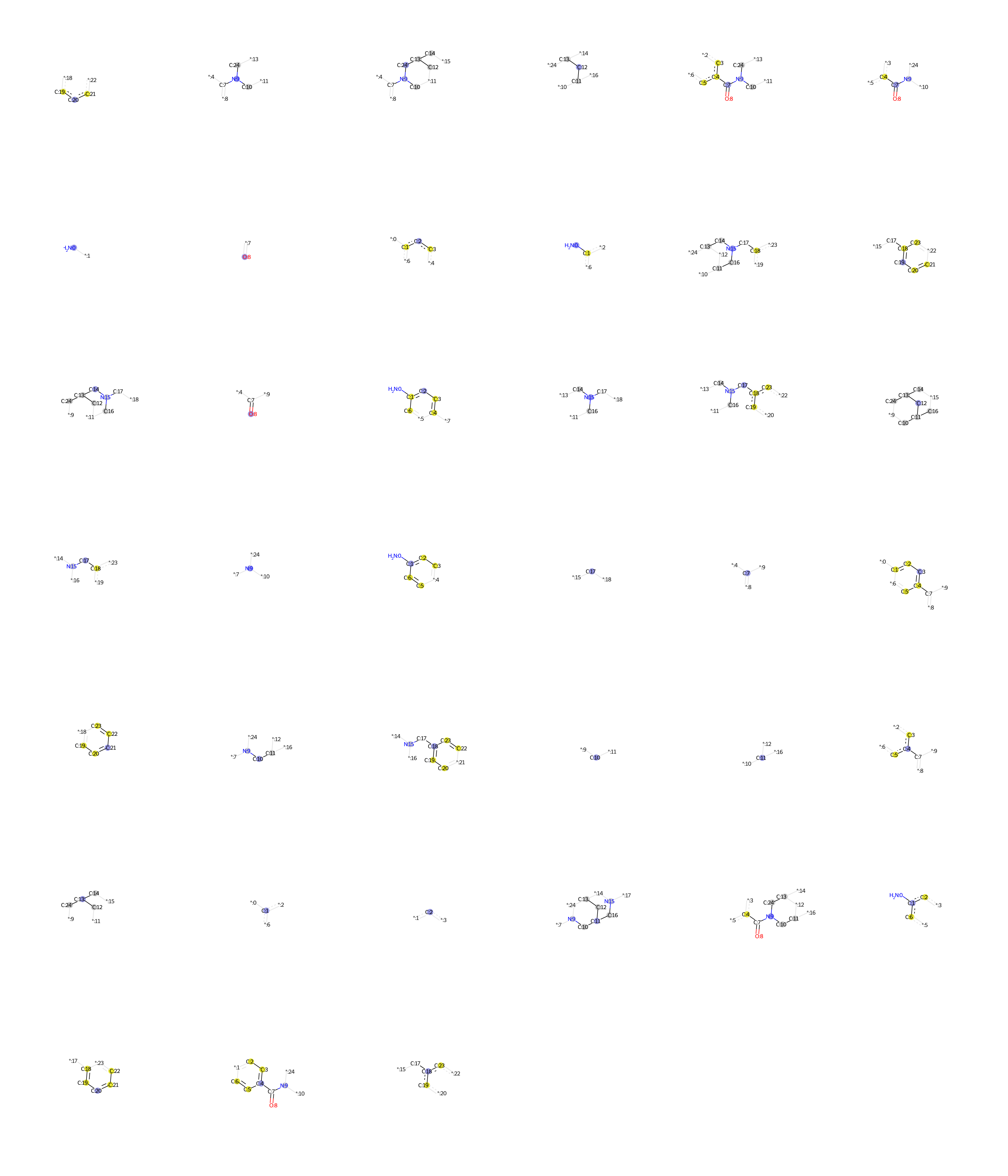

### Amiodarone.bmp

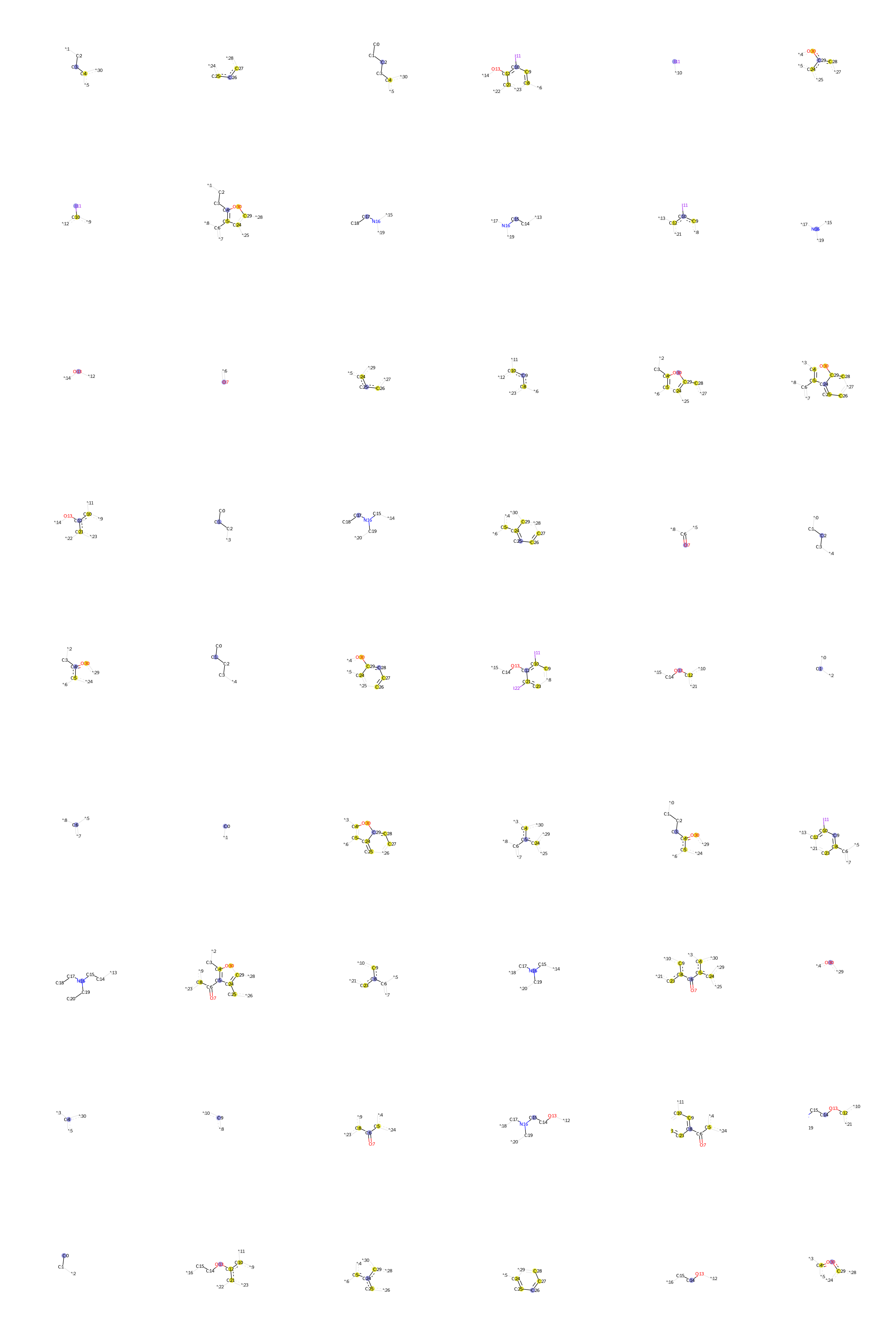

### Amitriptyline.bmp

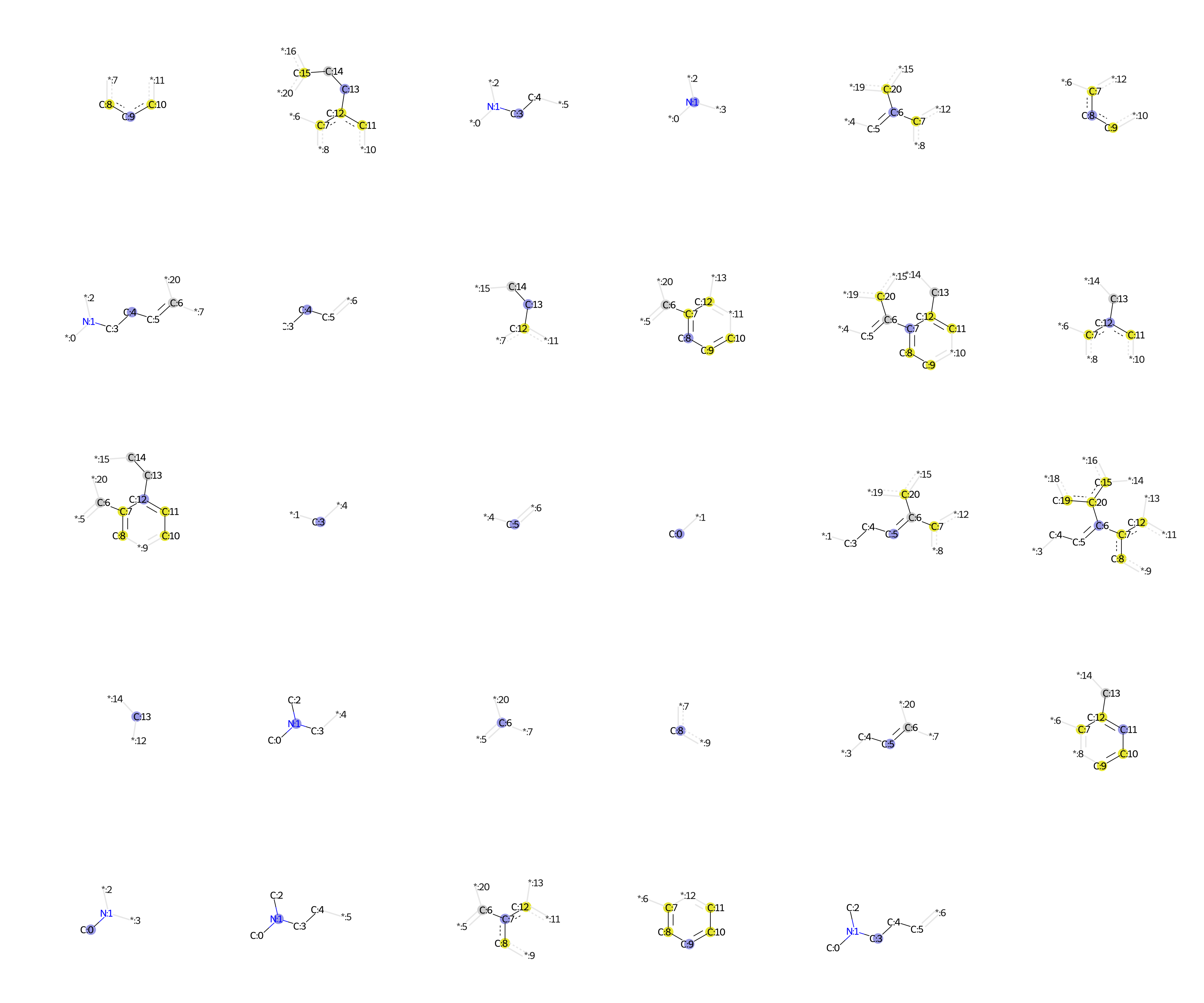

### Amsacrine.bmp

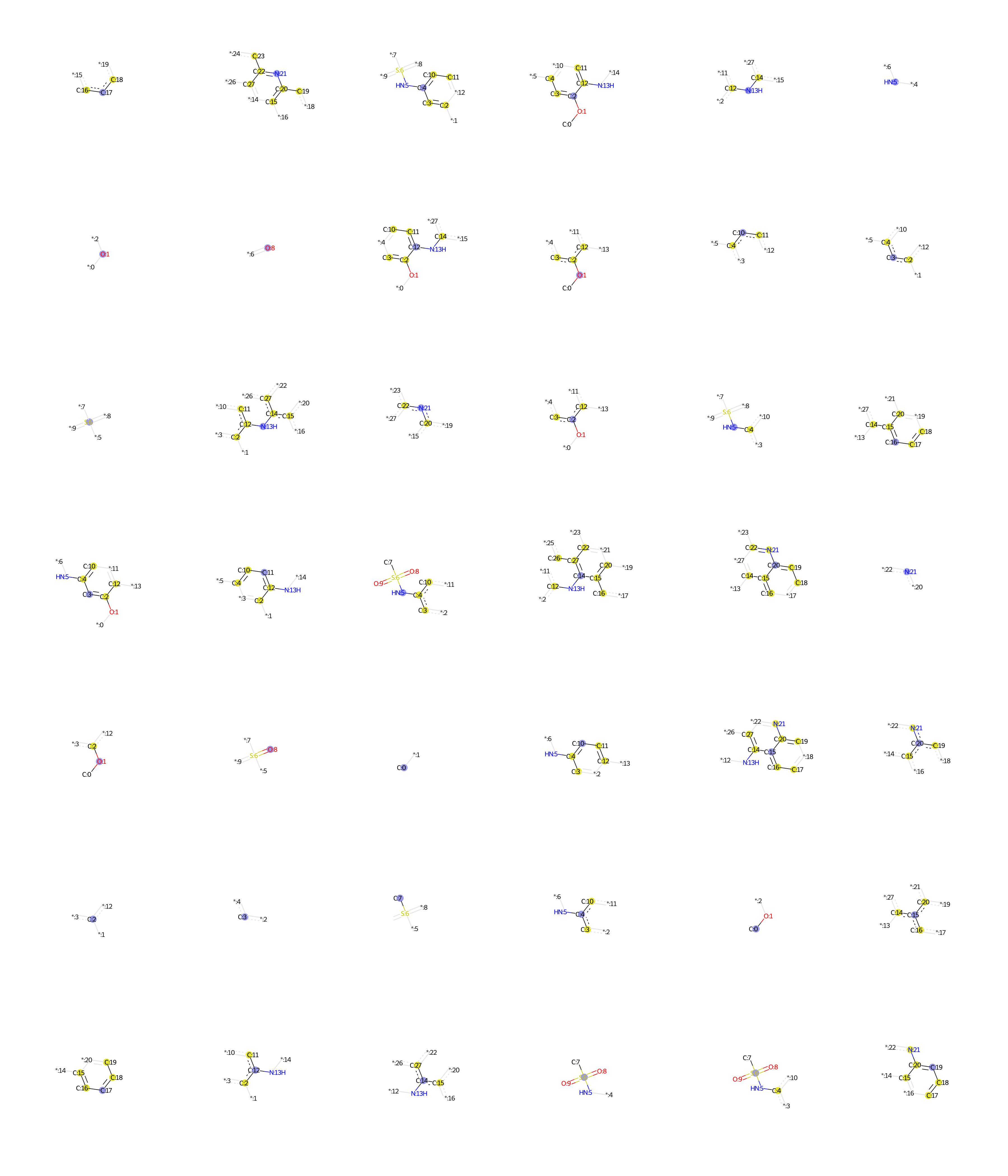

### Apomorphine.bmp

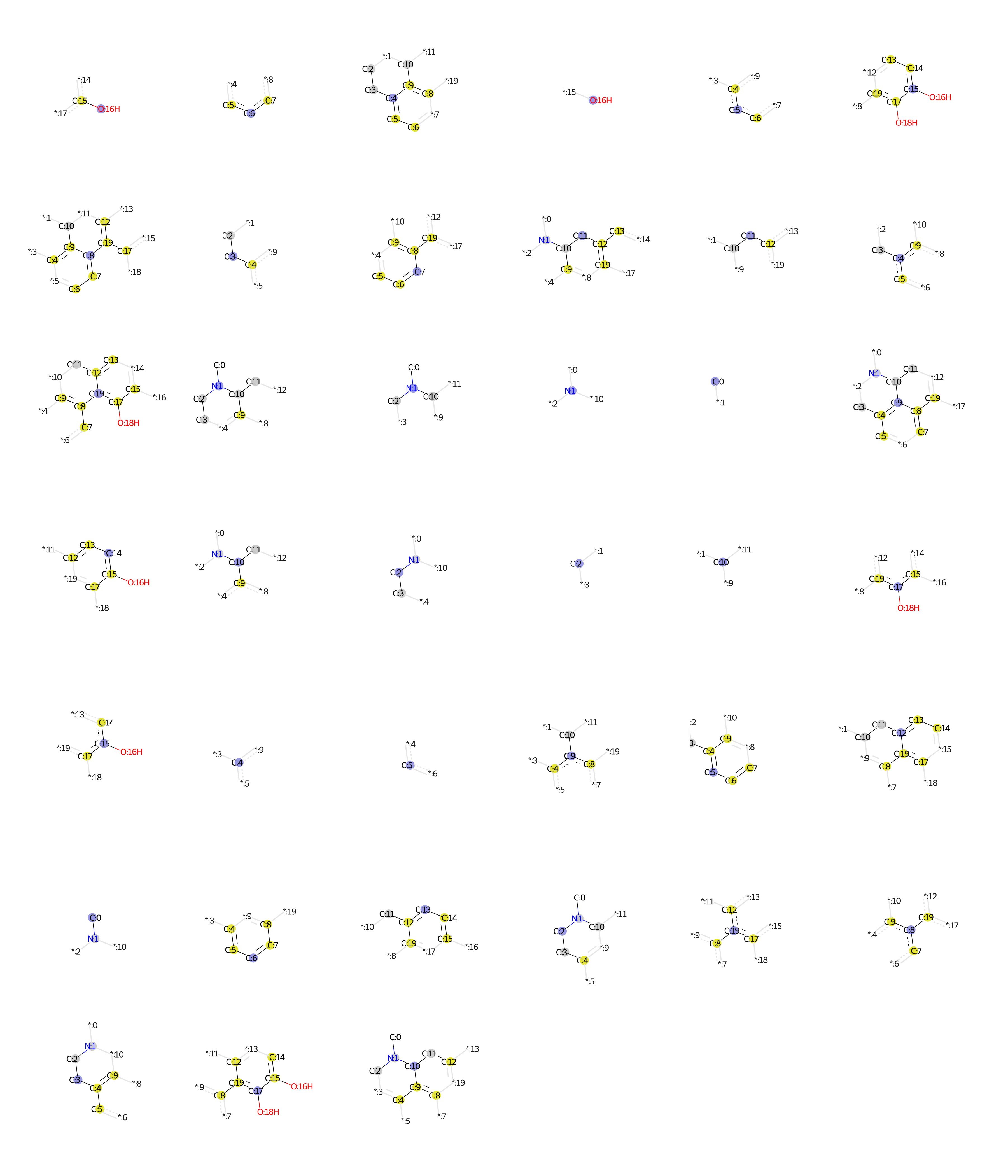

### Articaine.bmp

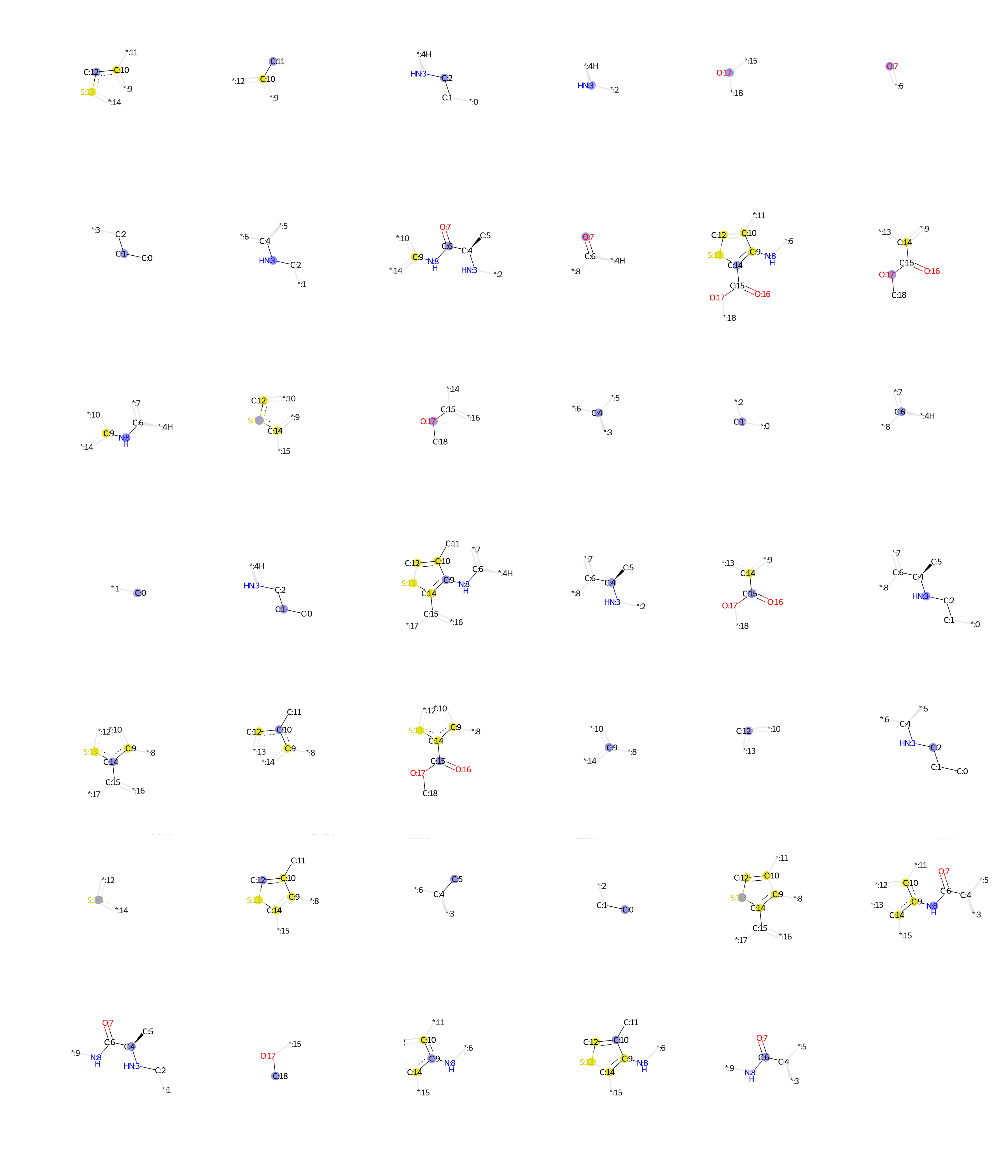

### Astemizole.bmp

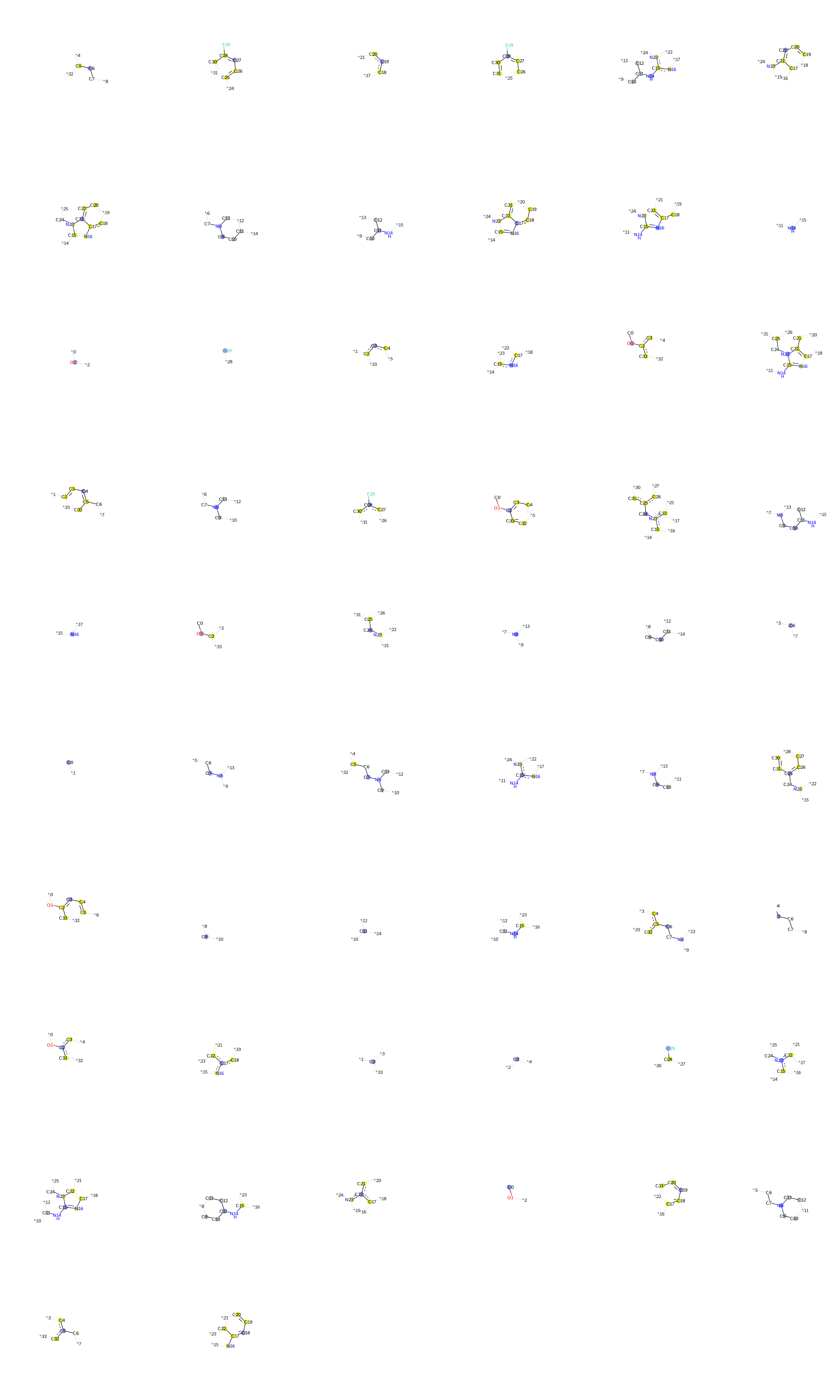

### Azimilide.bmp

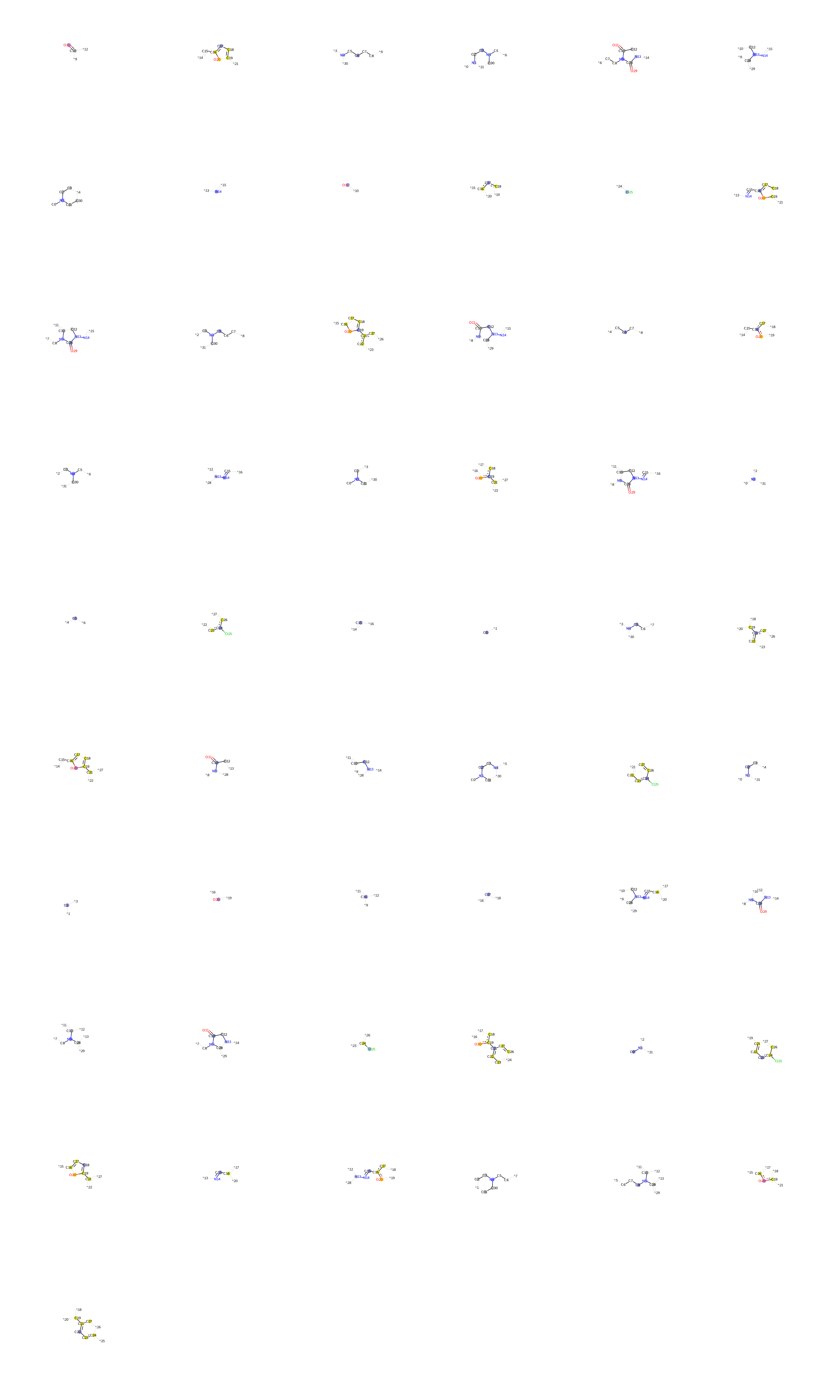

### b.bmp

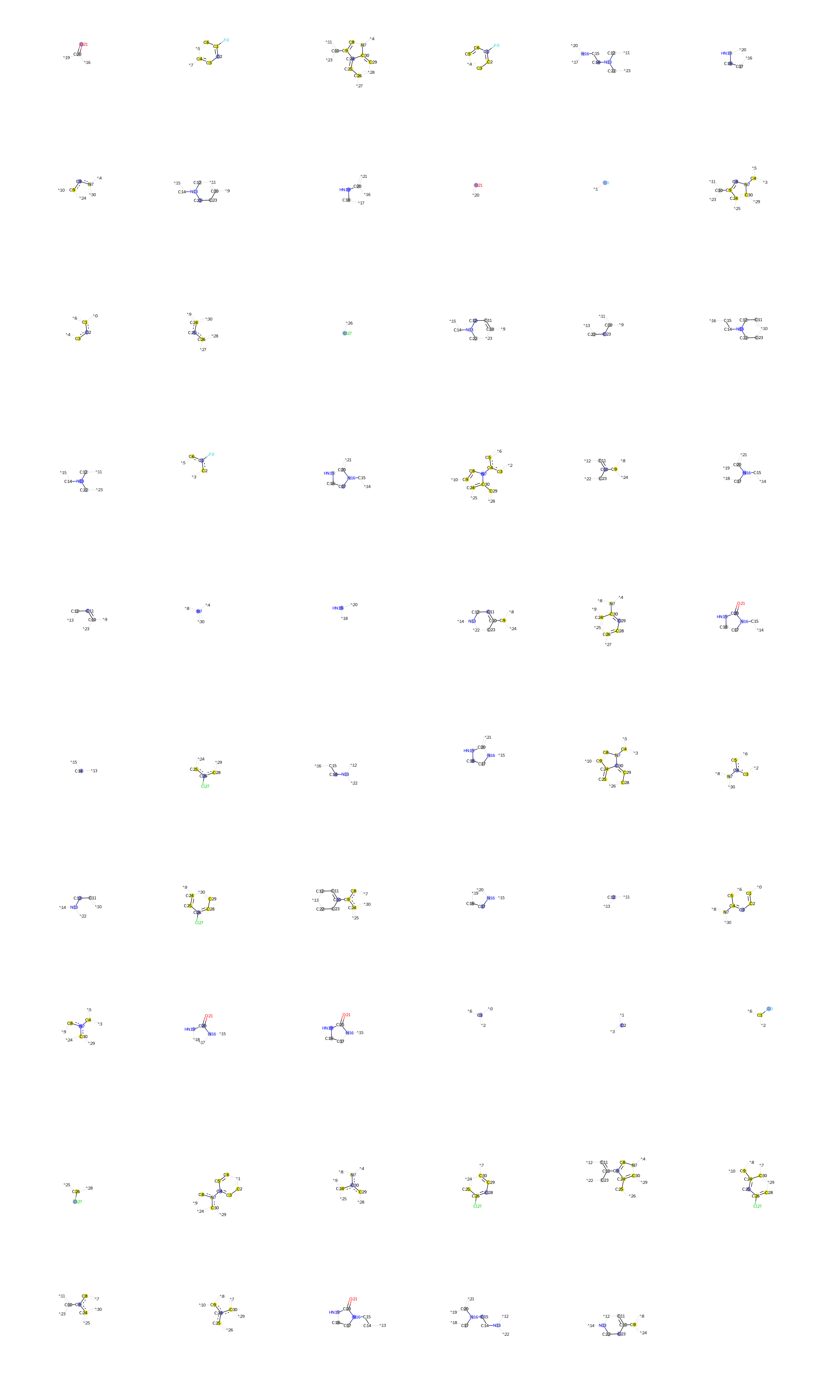

### Balofloxacin.bmp

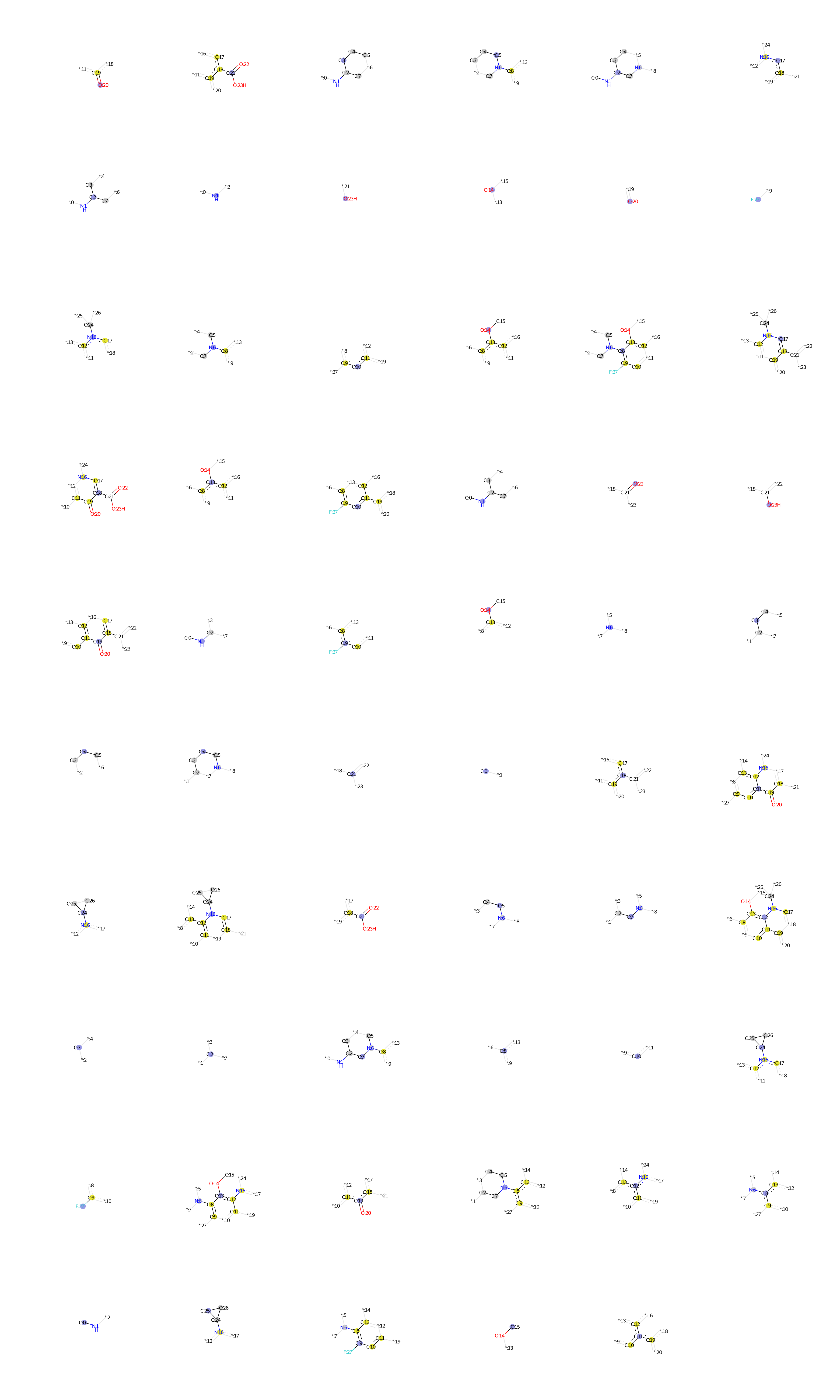

### Benperidol.bmp

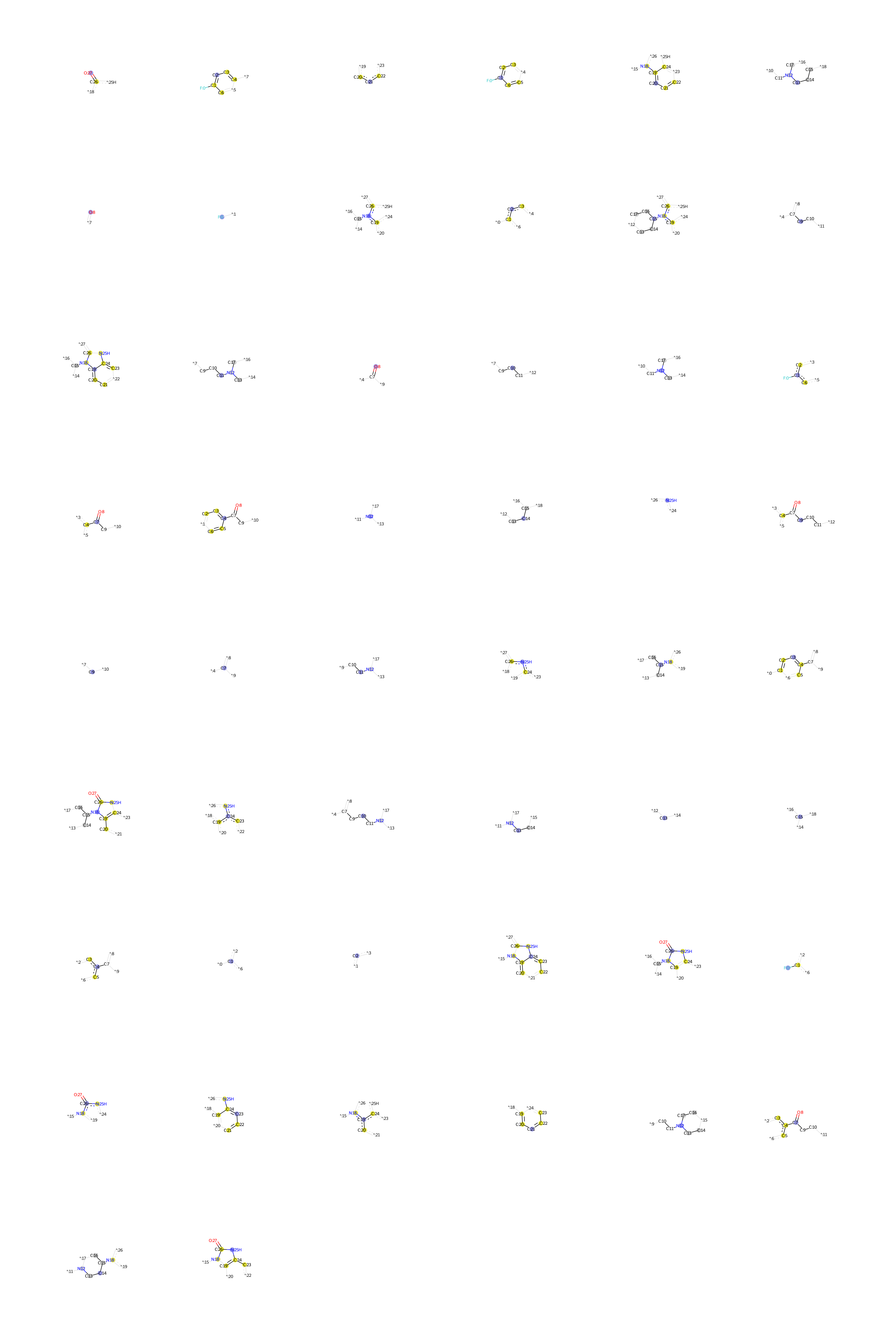

### Bepridil.bmp

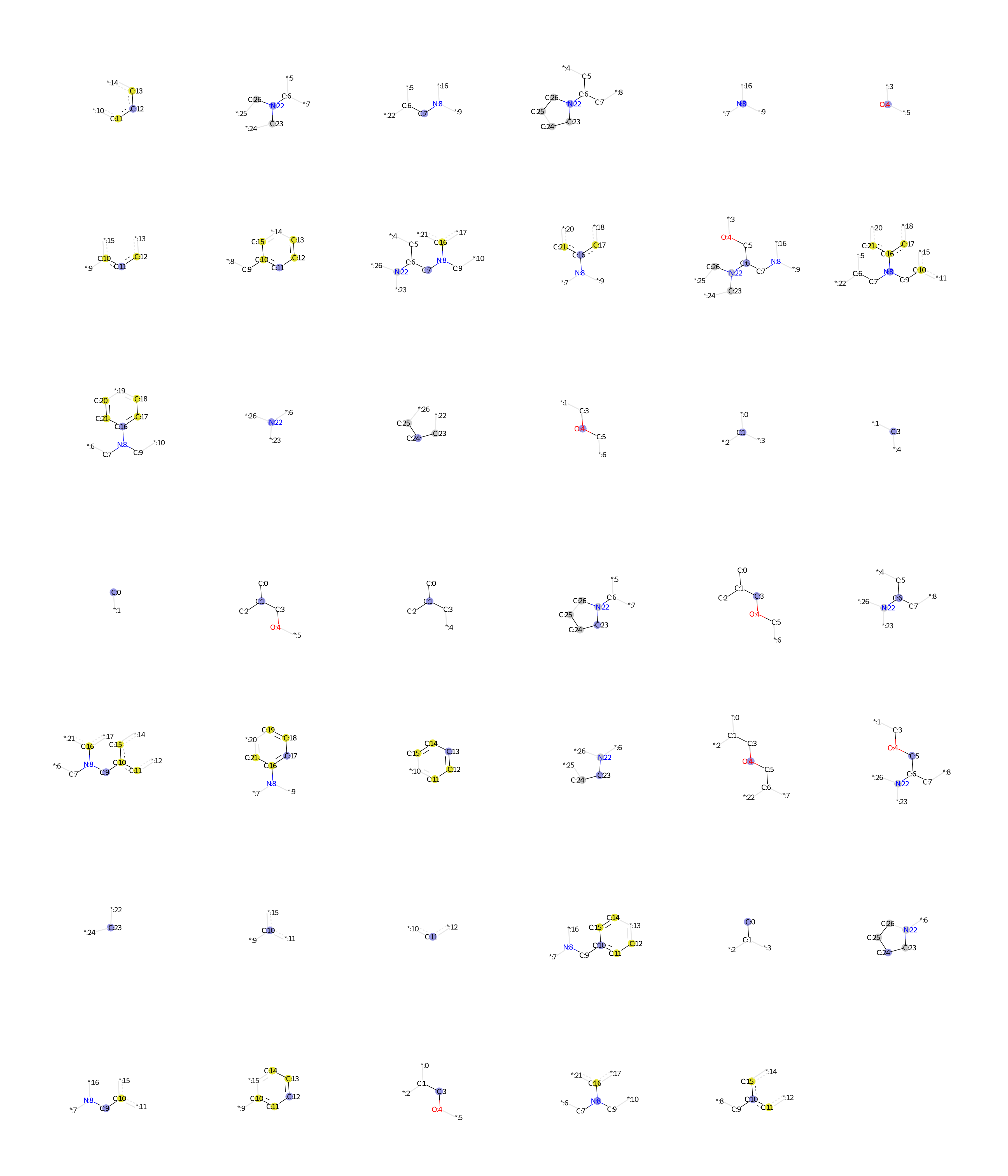

### Bisindolylmaleimide.bmp

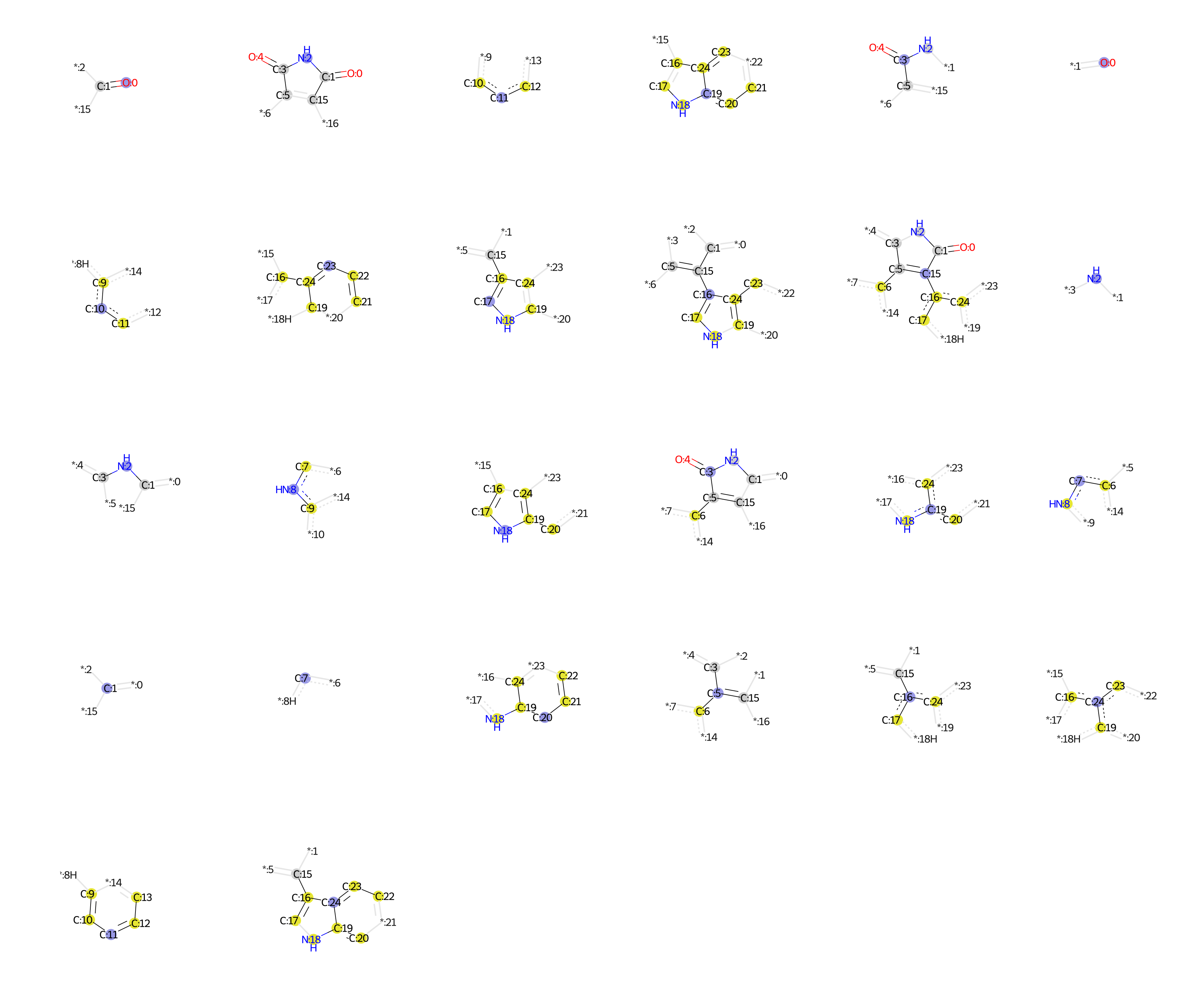

### BRL.bmp

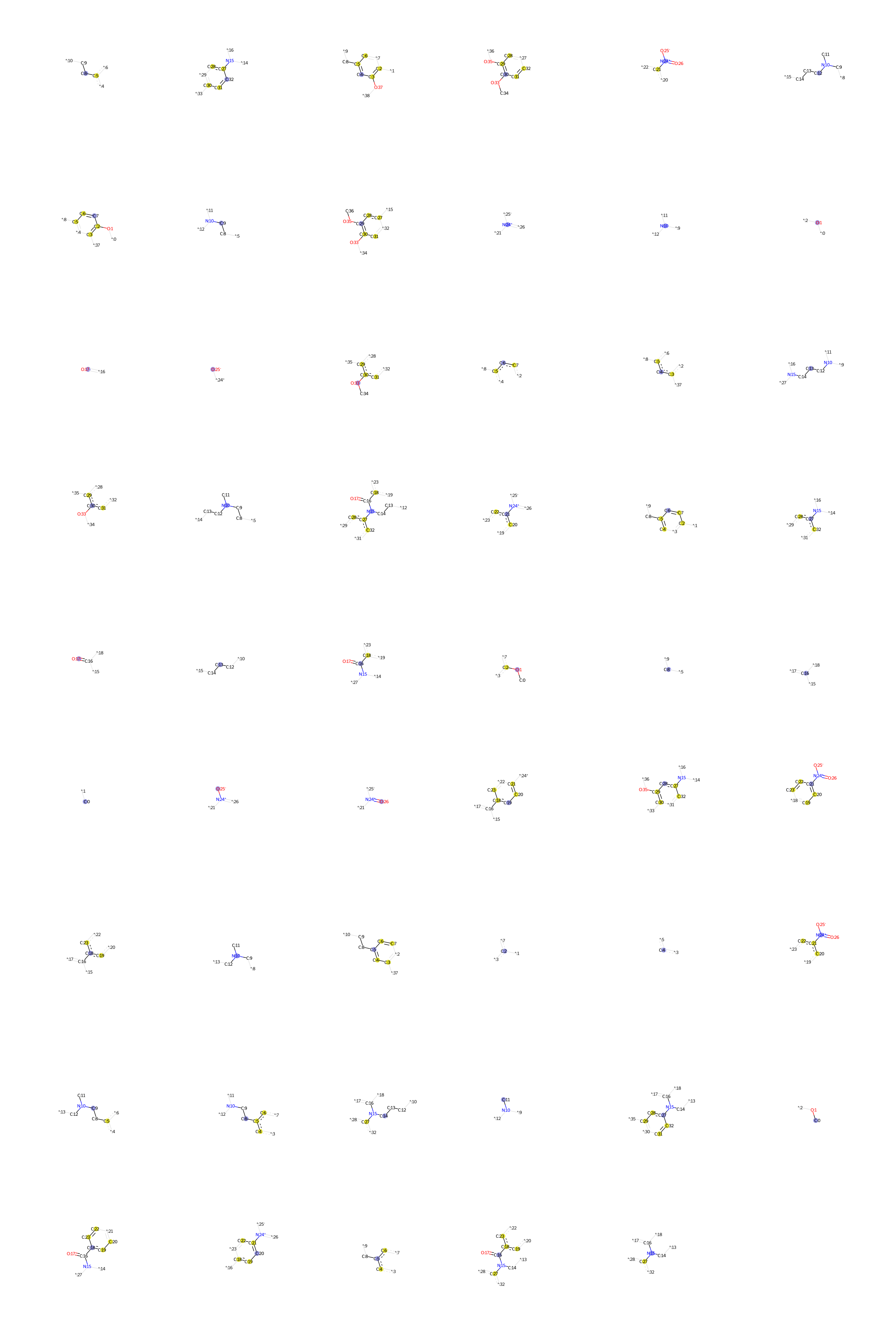

### Brompheniramine.bmp

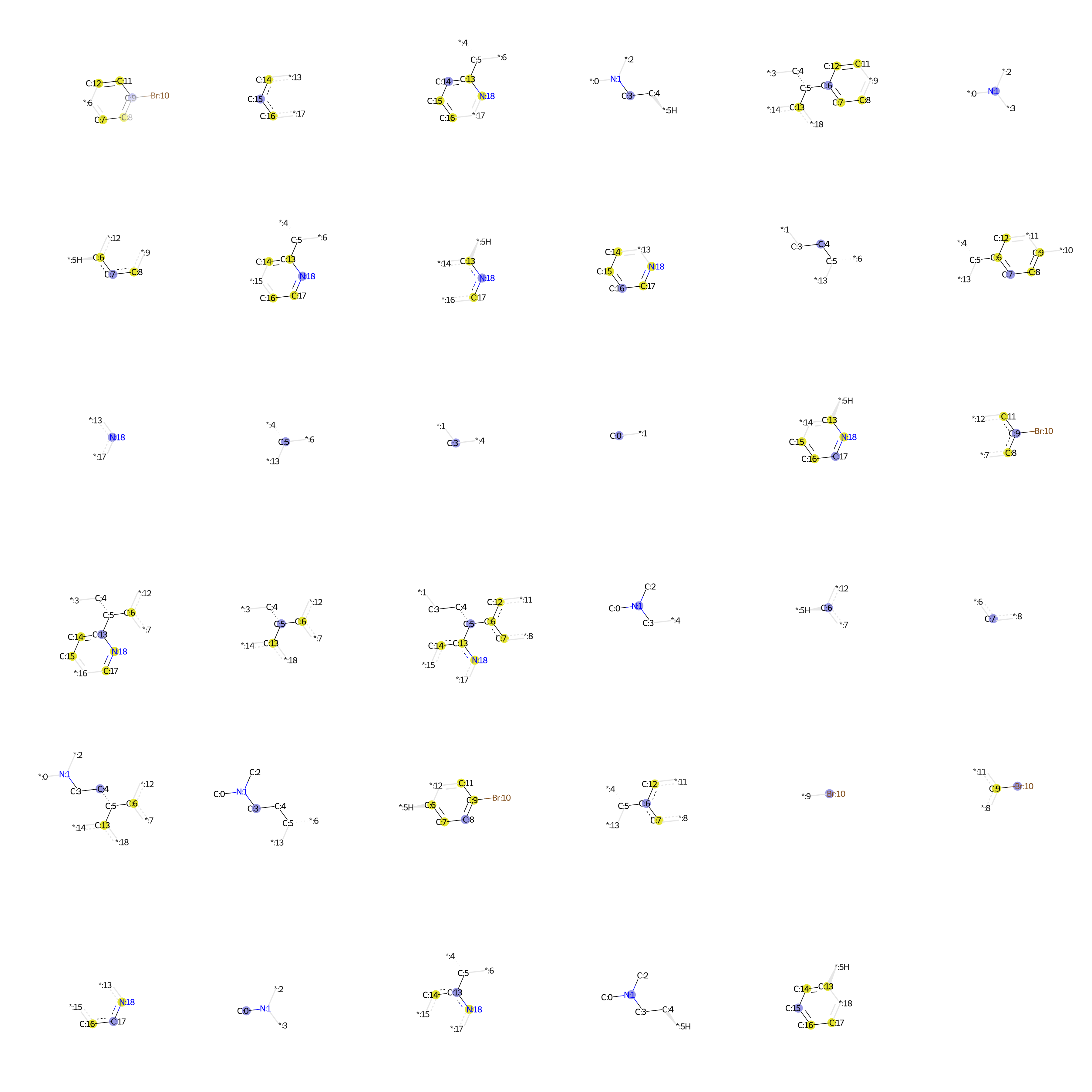

### Bupivacaine.bmp

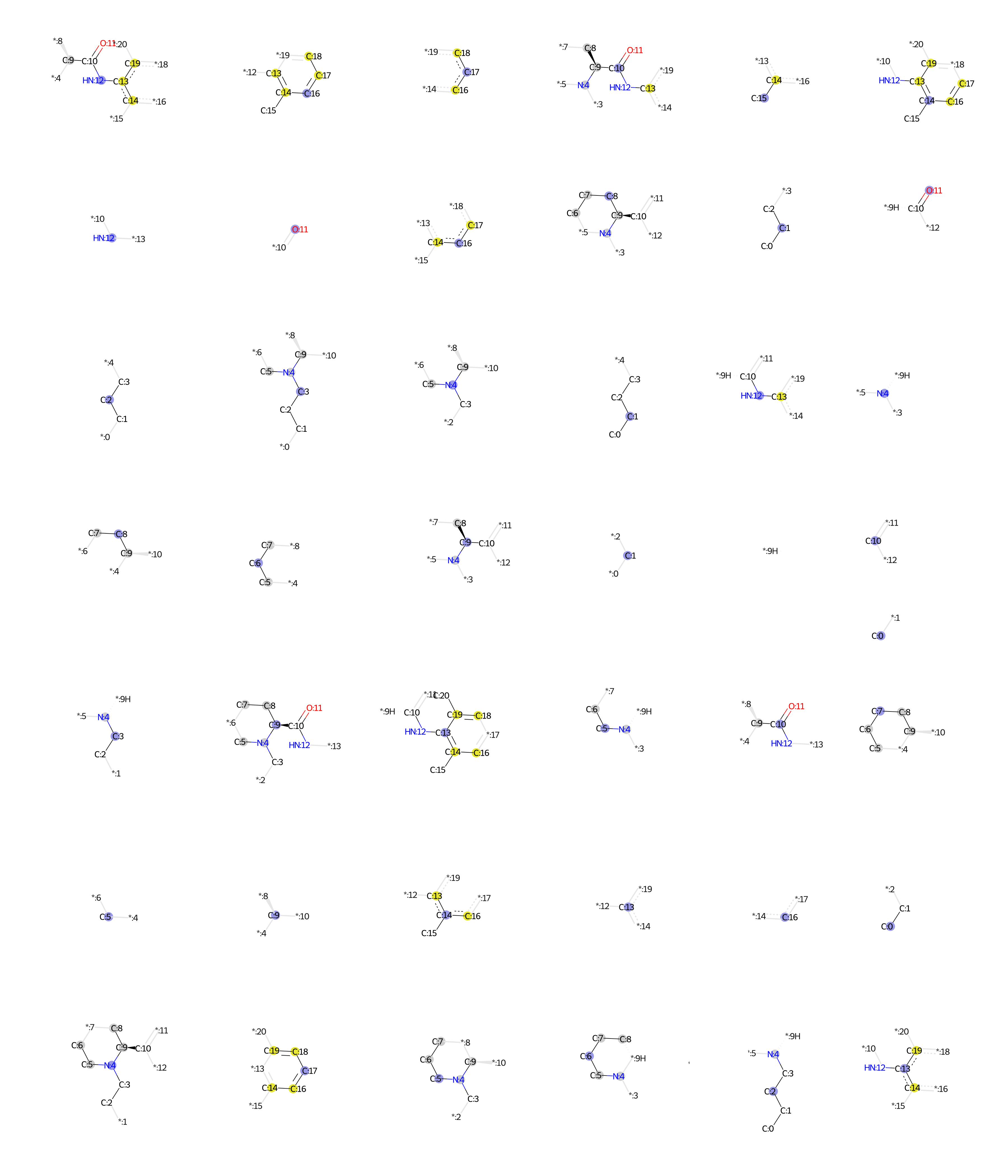

### c.bmp

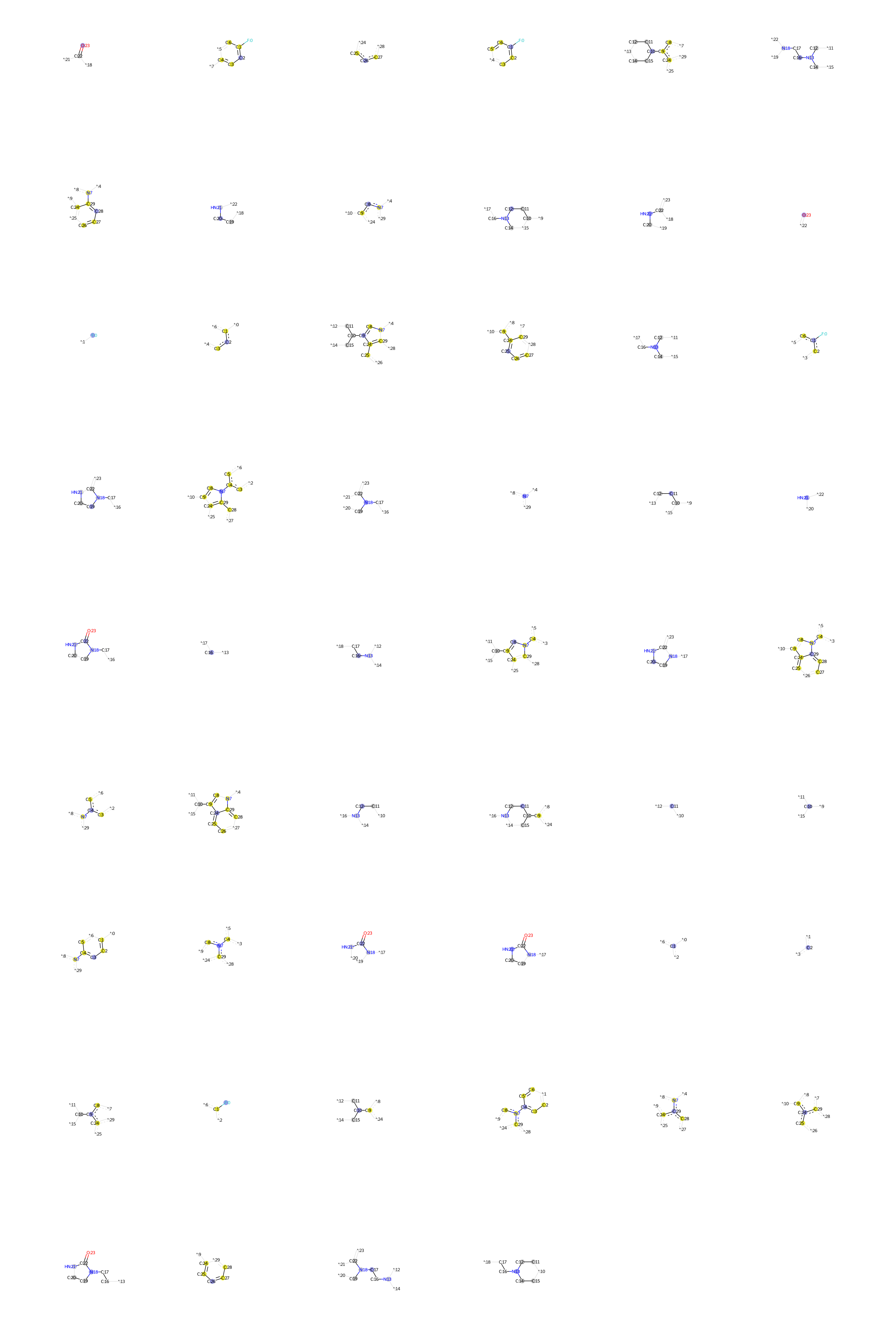

### Carbamazepine.bmp

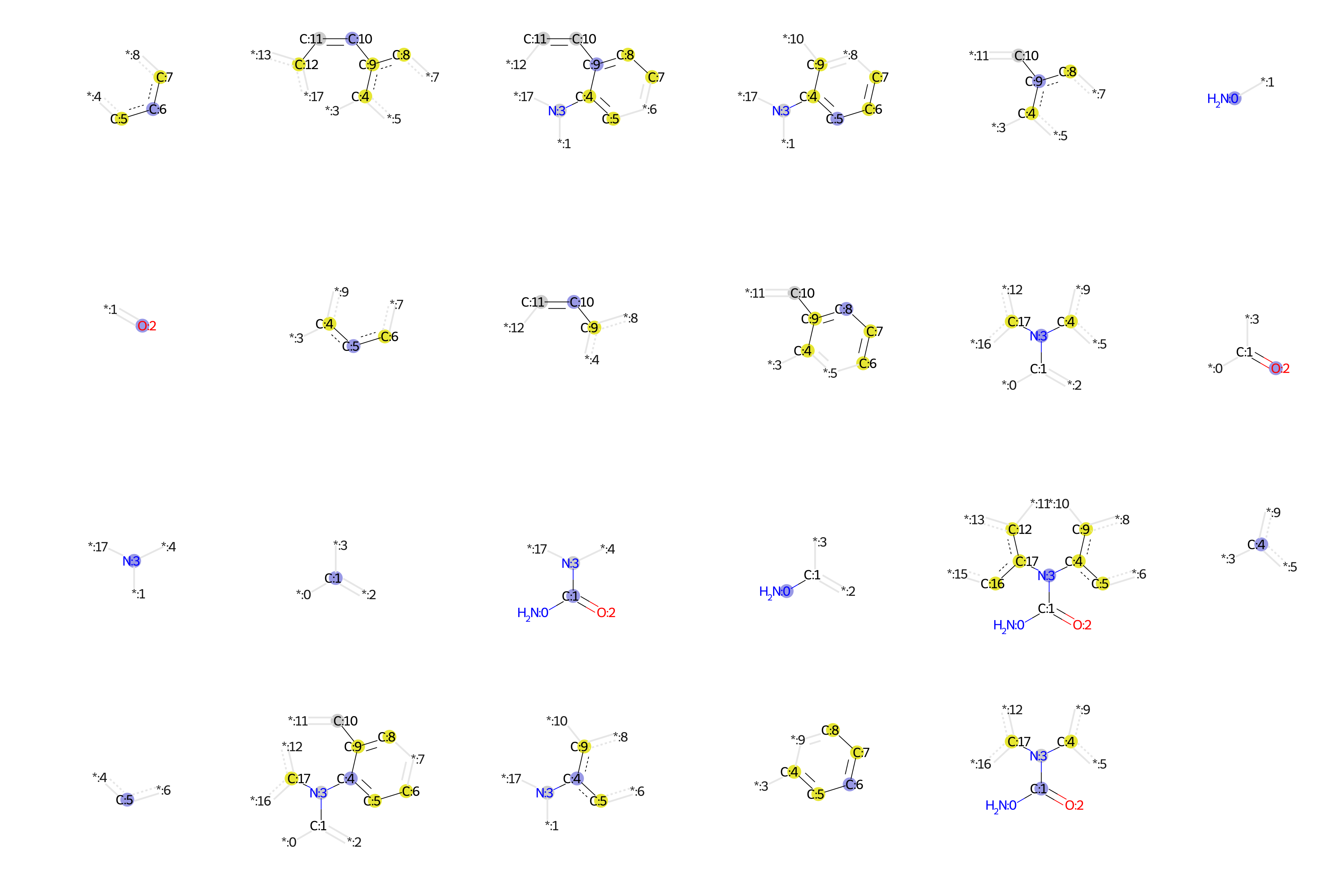

### Carvedilol.bmp

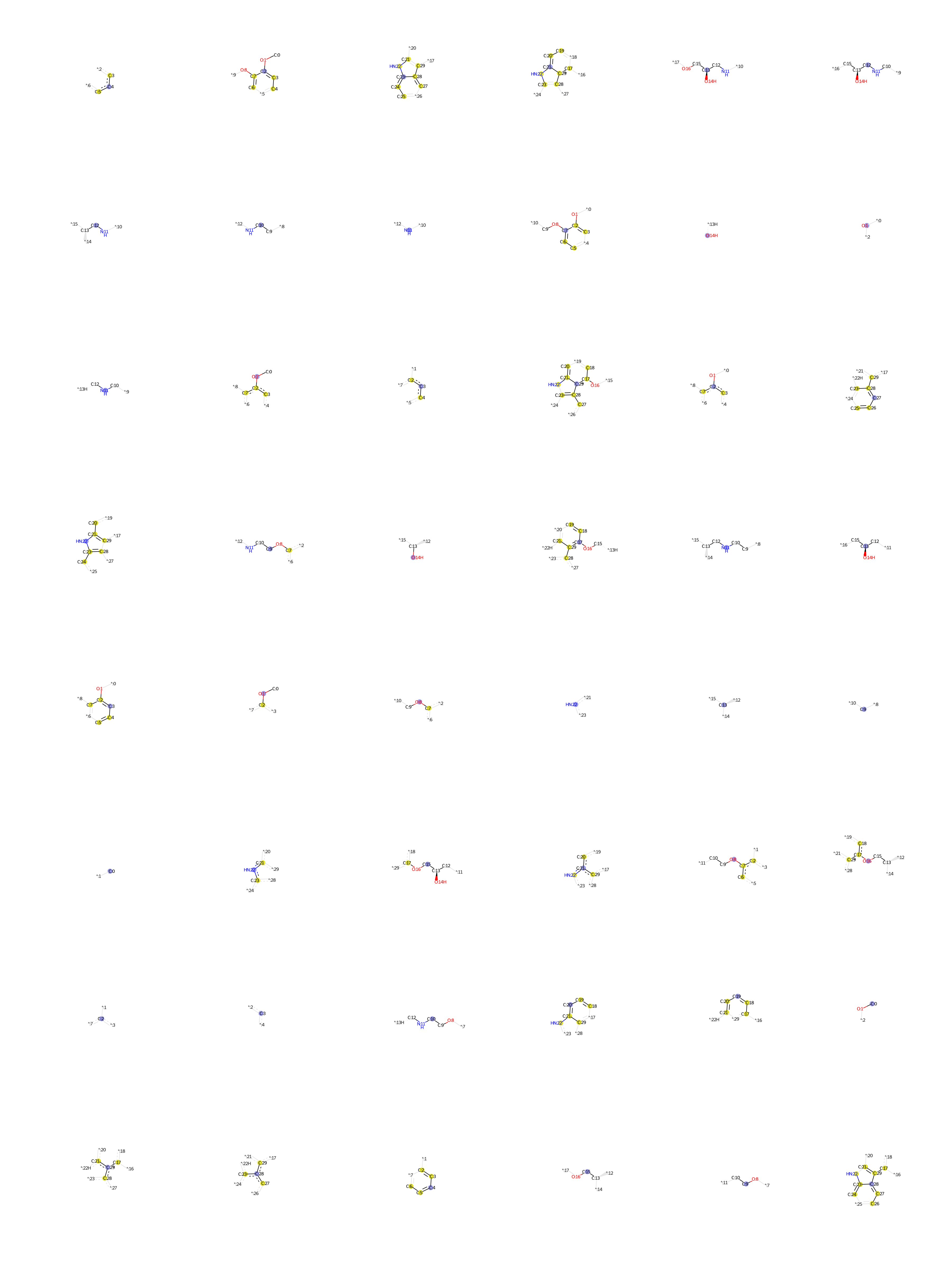

### Chloroquine.bmp

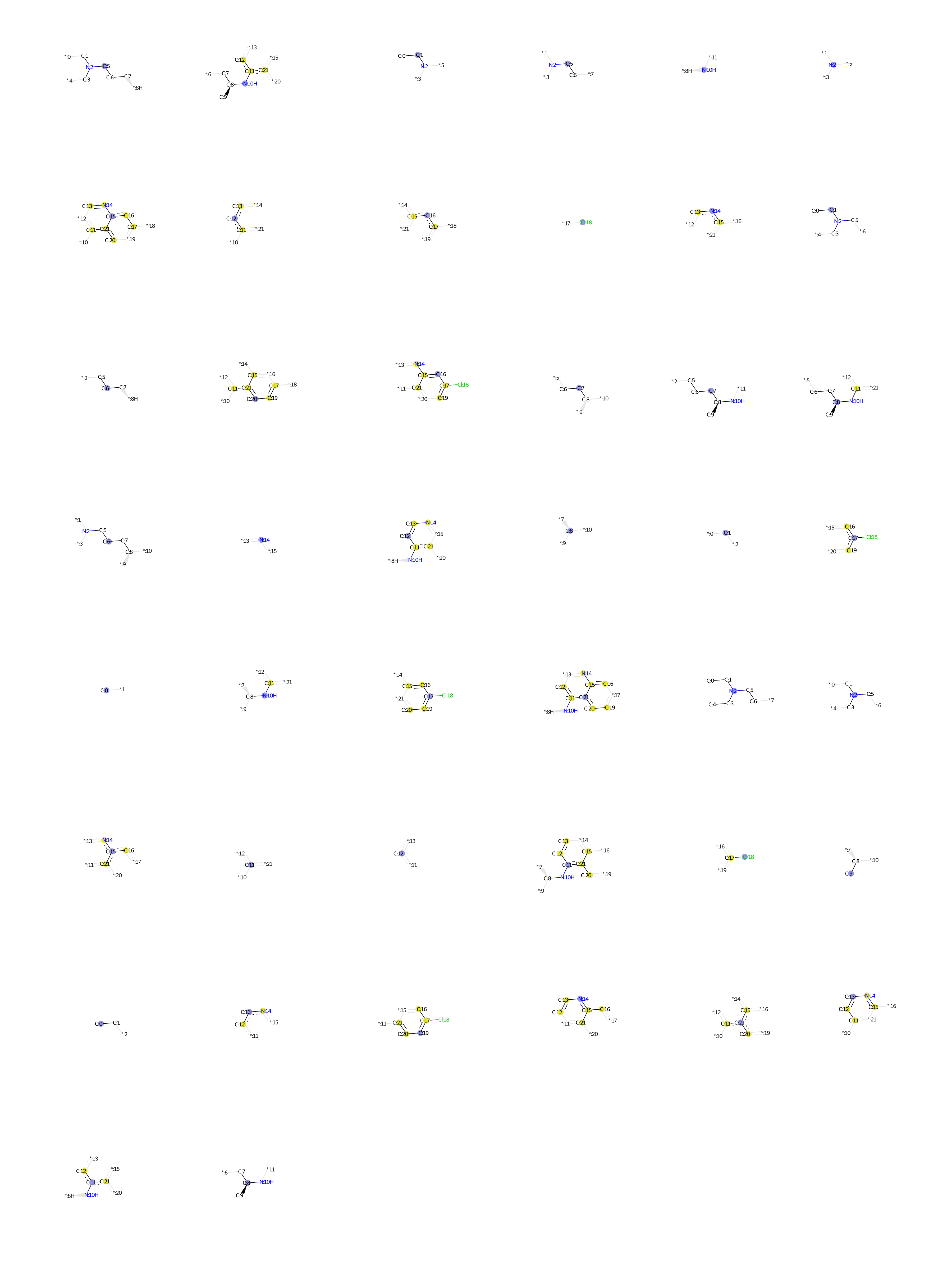

### Chlorpheniramine.bmp

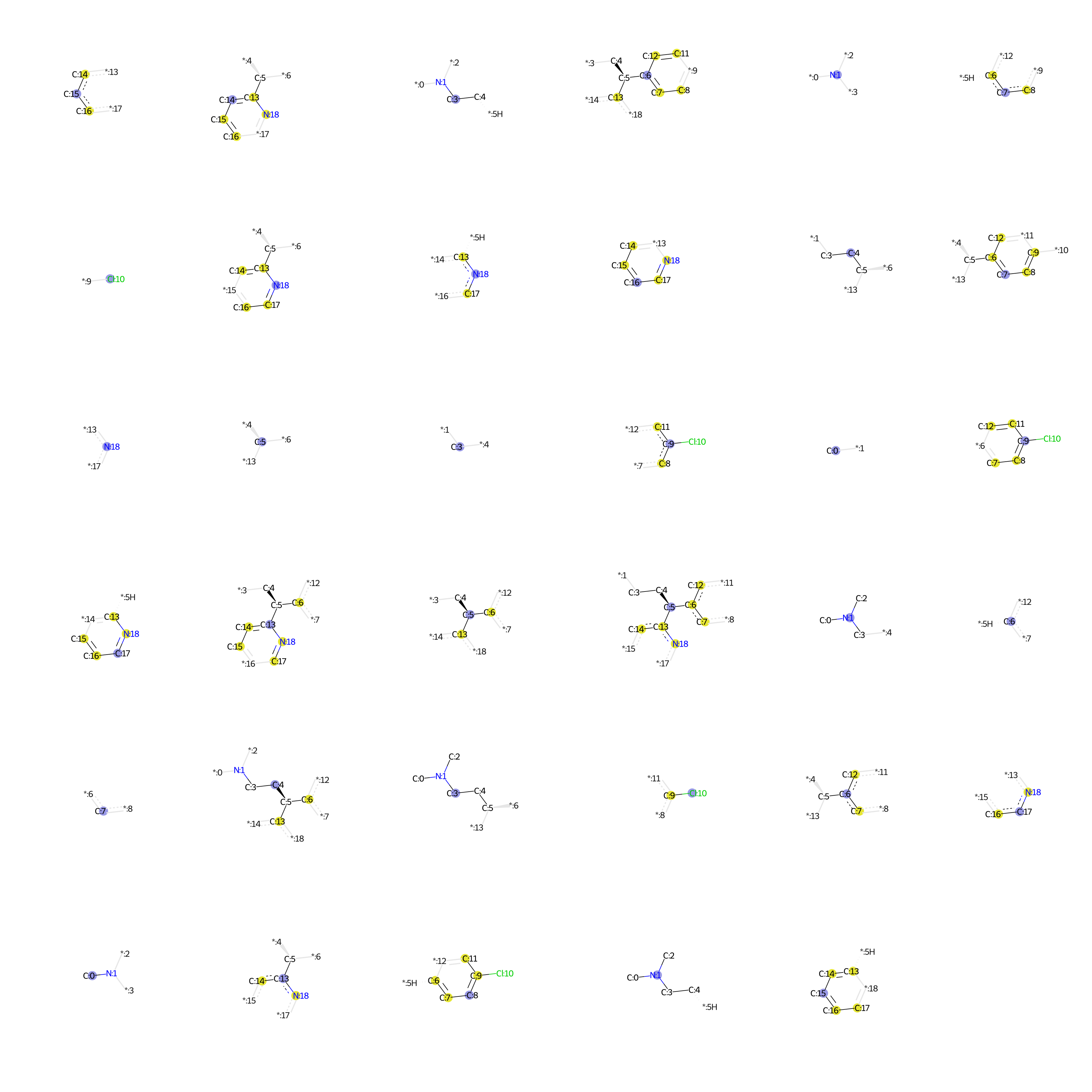

### Chlorpromazine.bmp

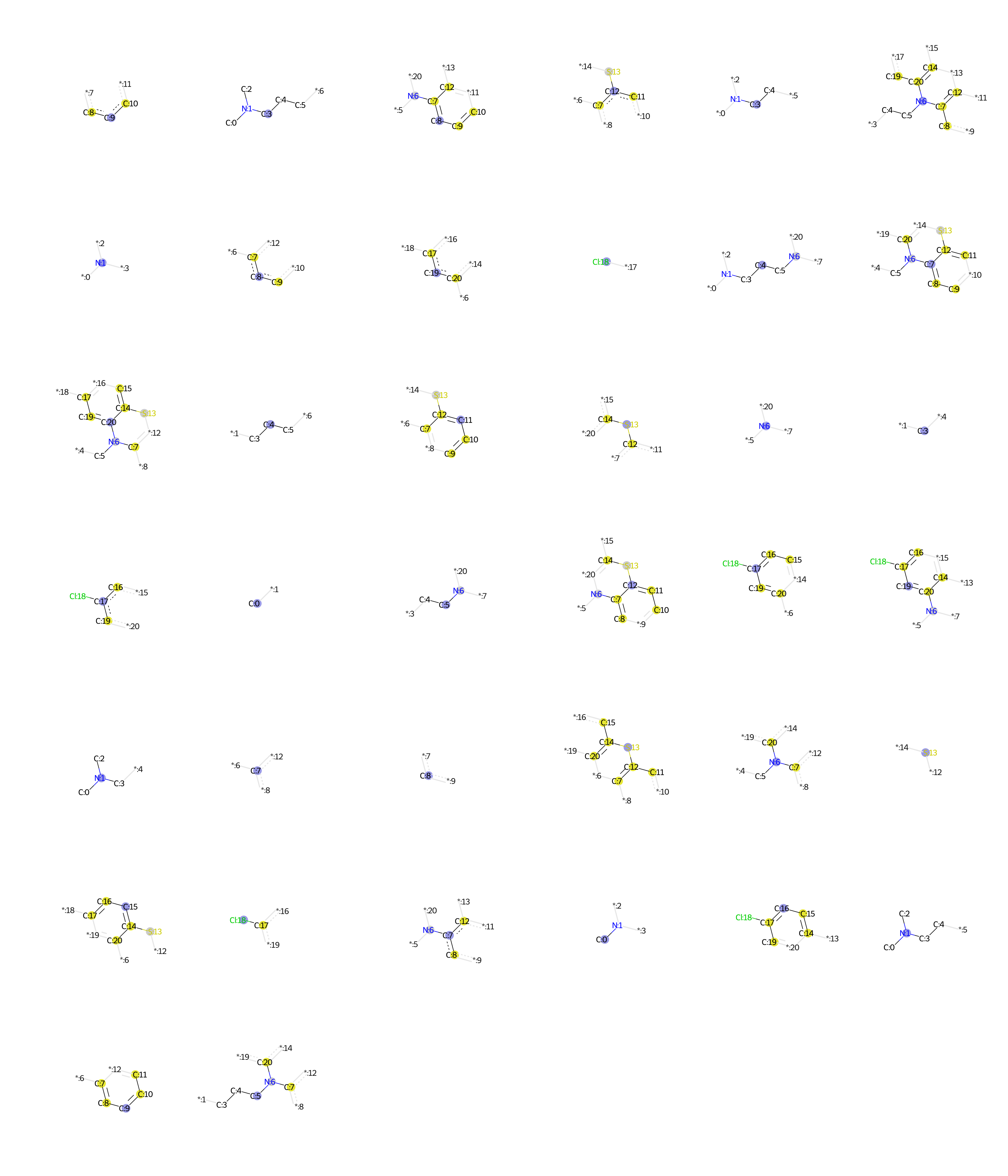

### Cibenzoline.bmp

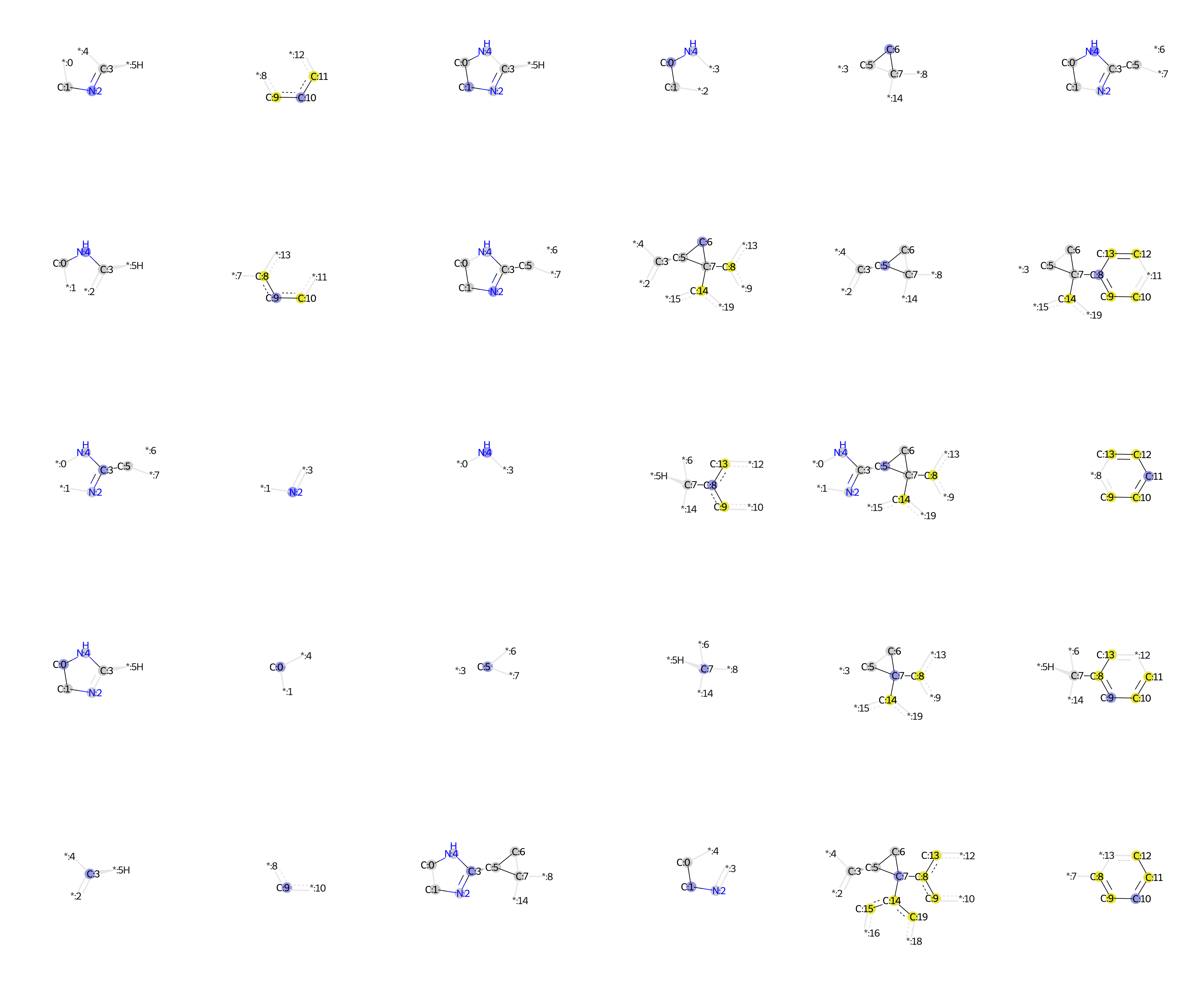

### Ciprofloxacin.bmp

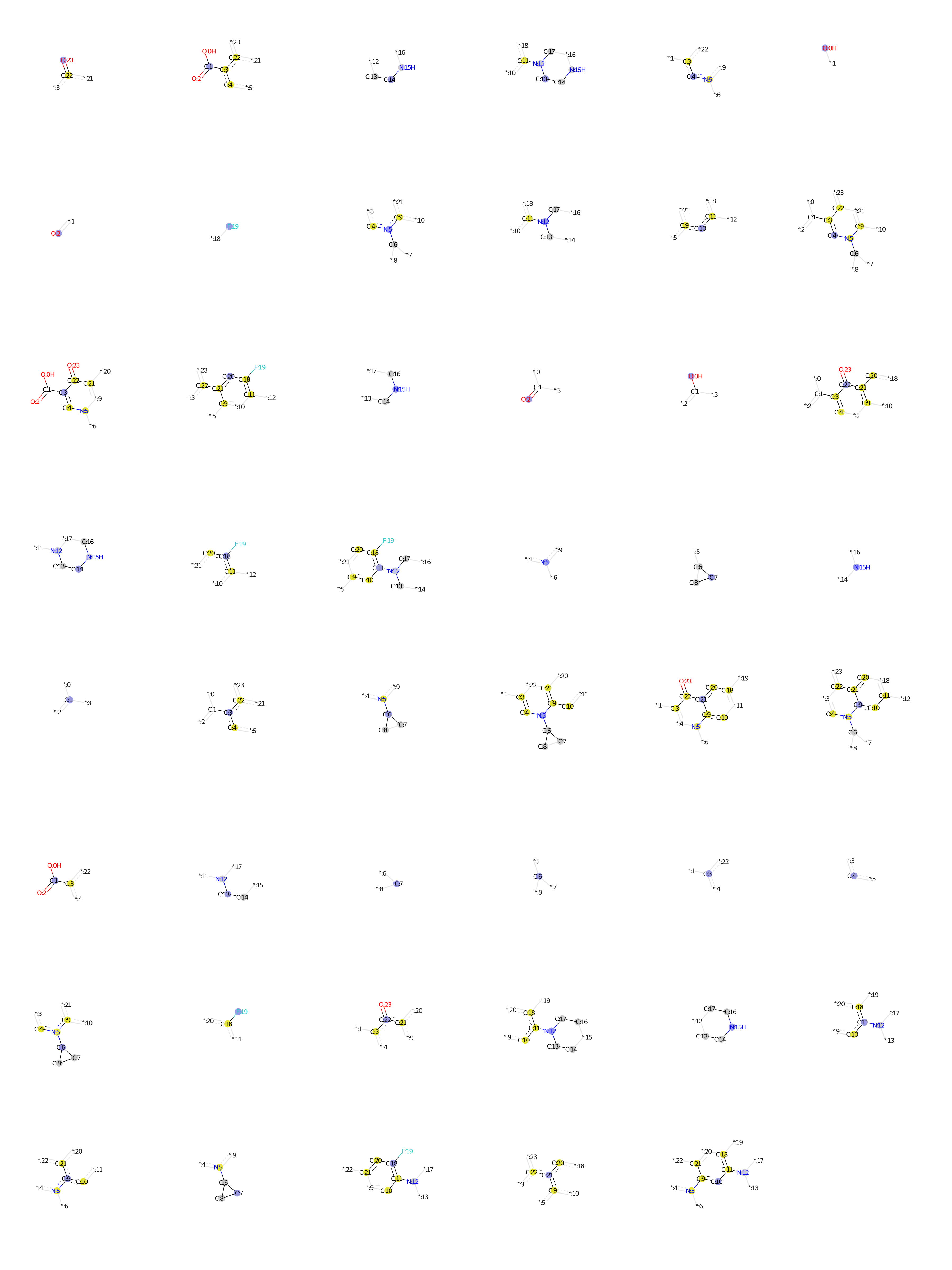

### Citalopram.bmp

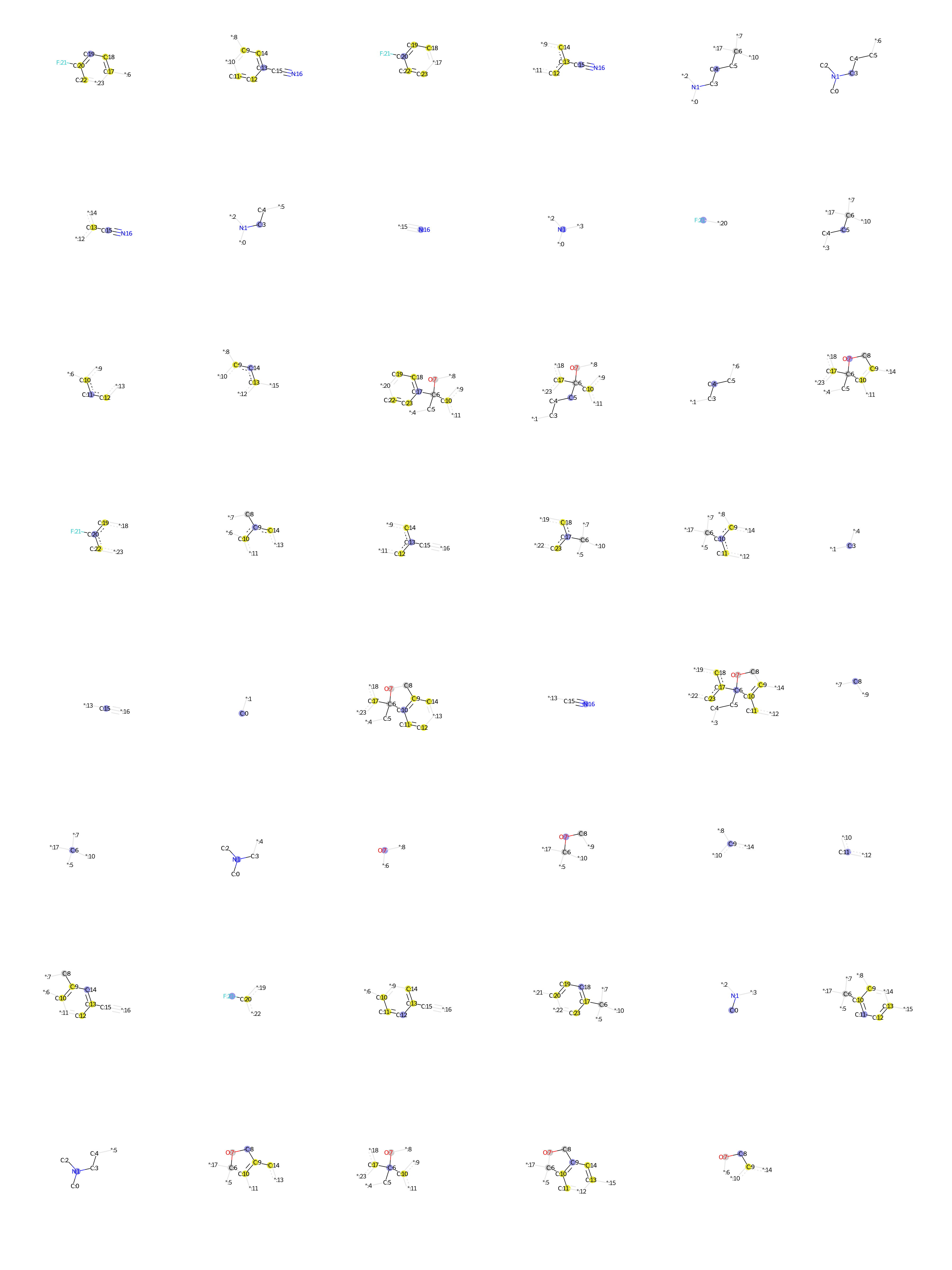

### Clebopride.bmp

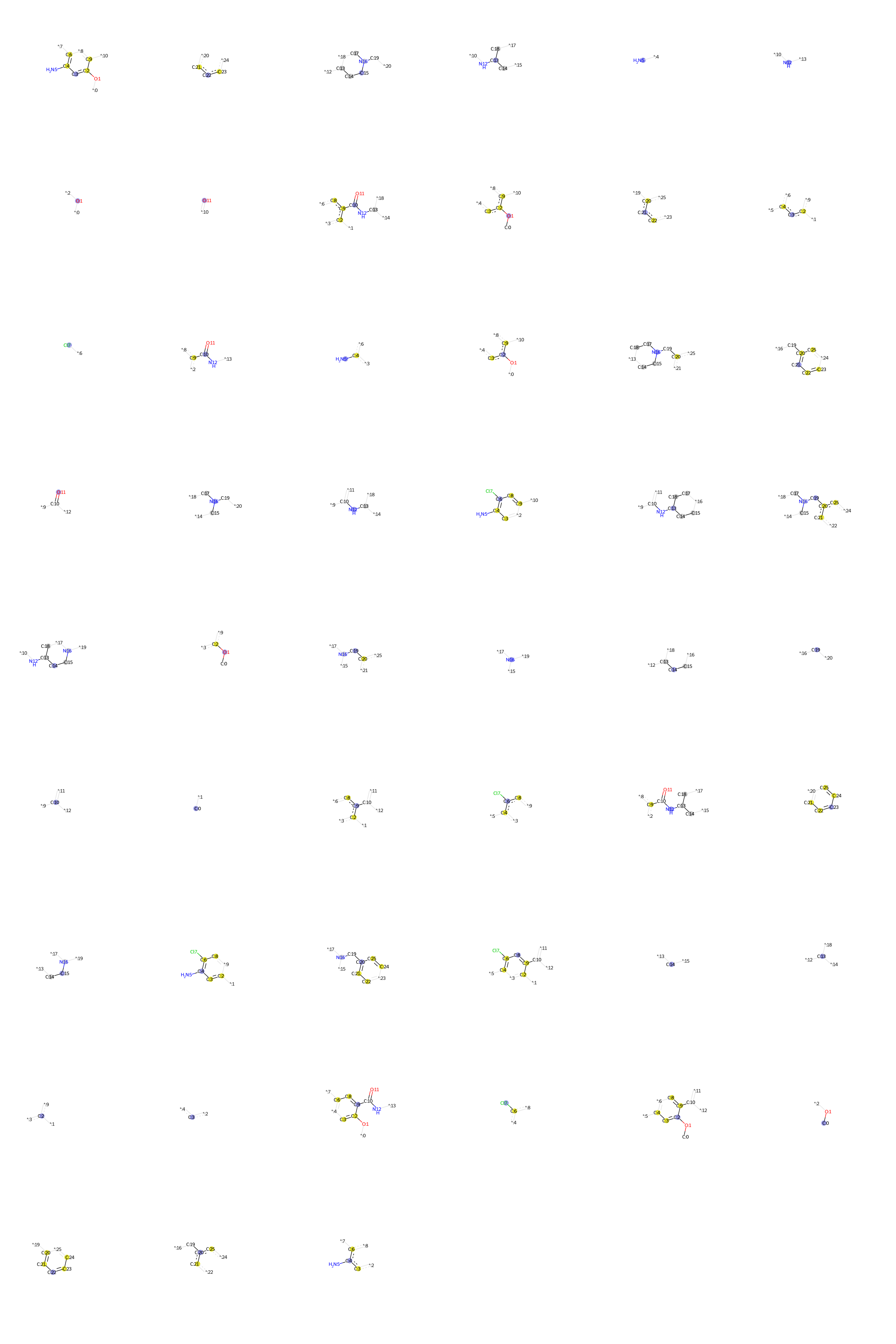

### Clemastine.bmp

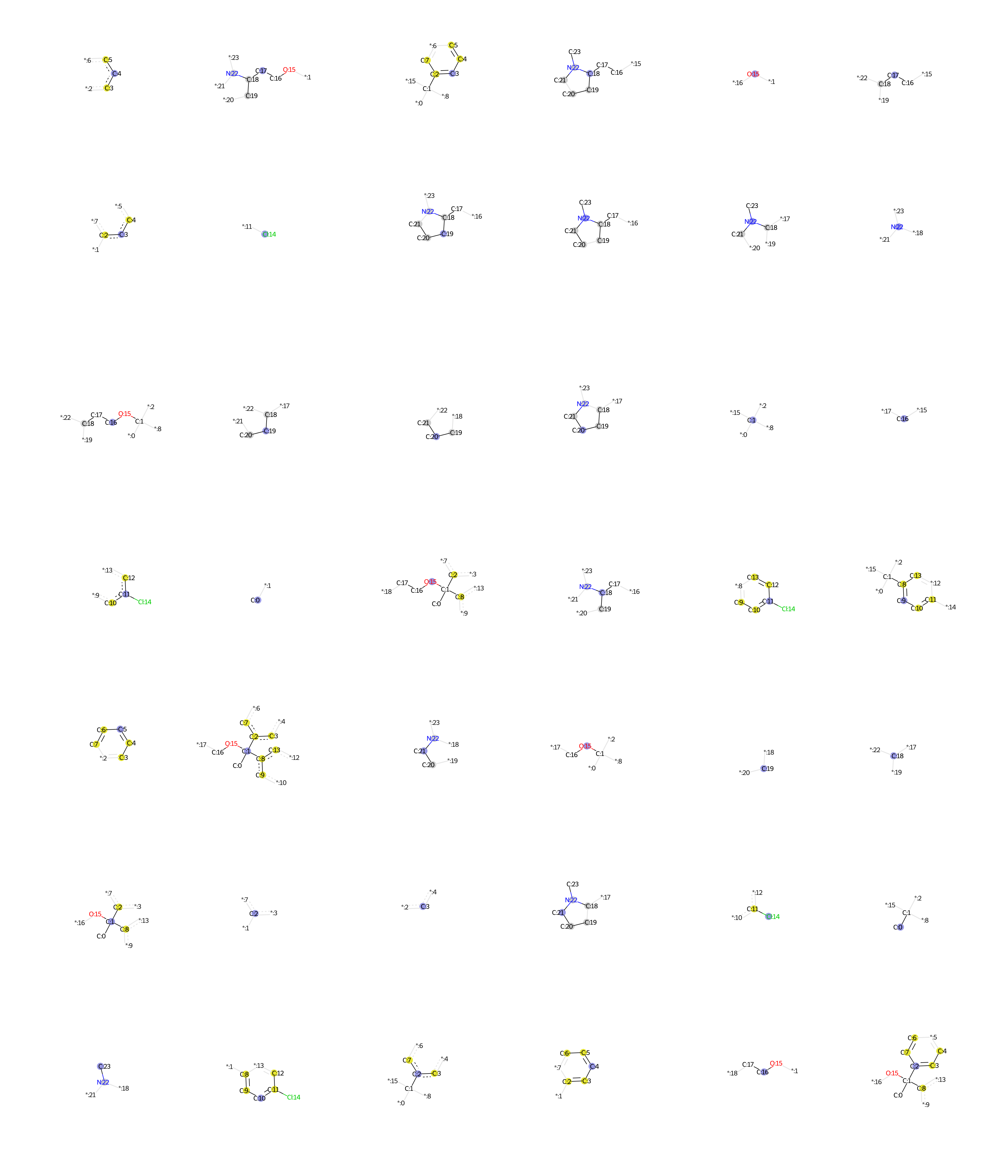

### Clomiphene.bmp

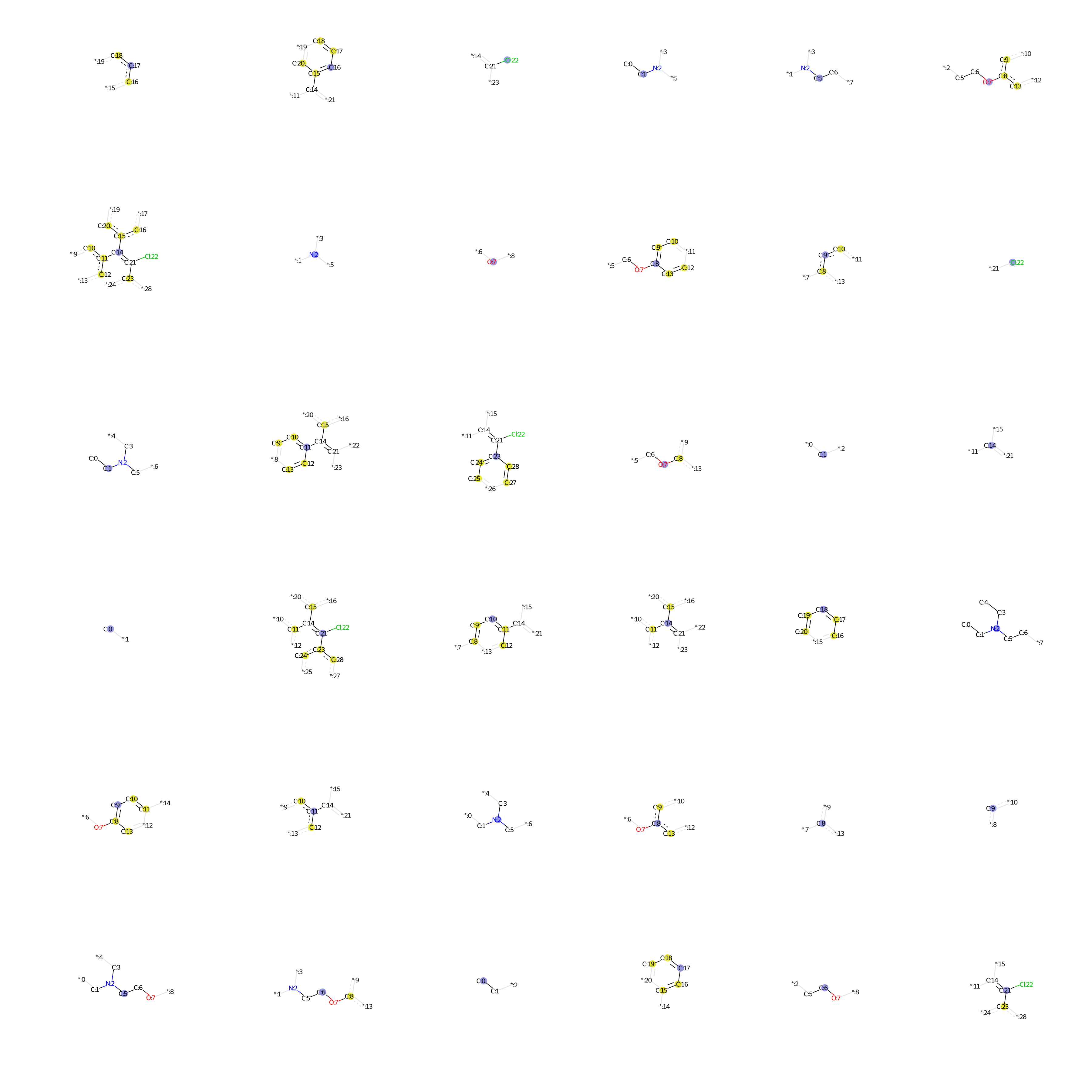

### Clotrimazole.bmp

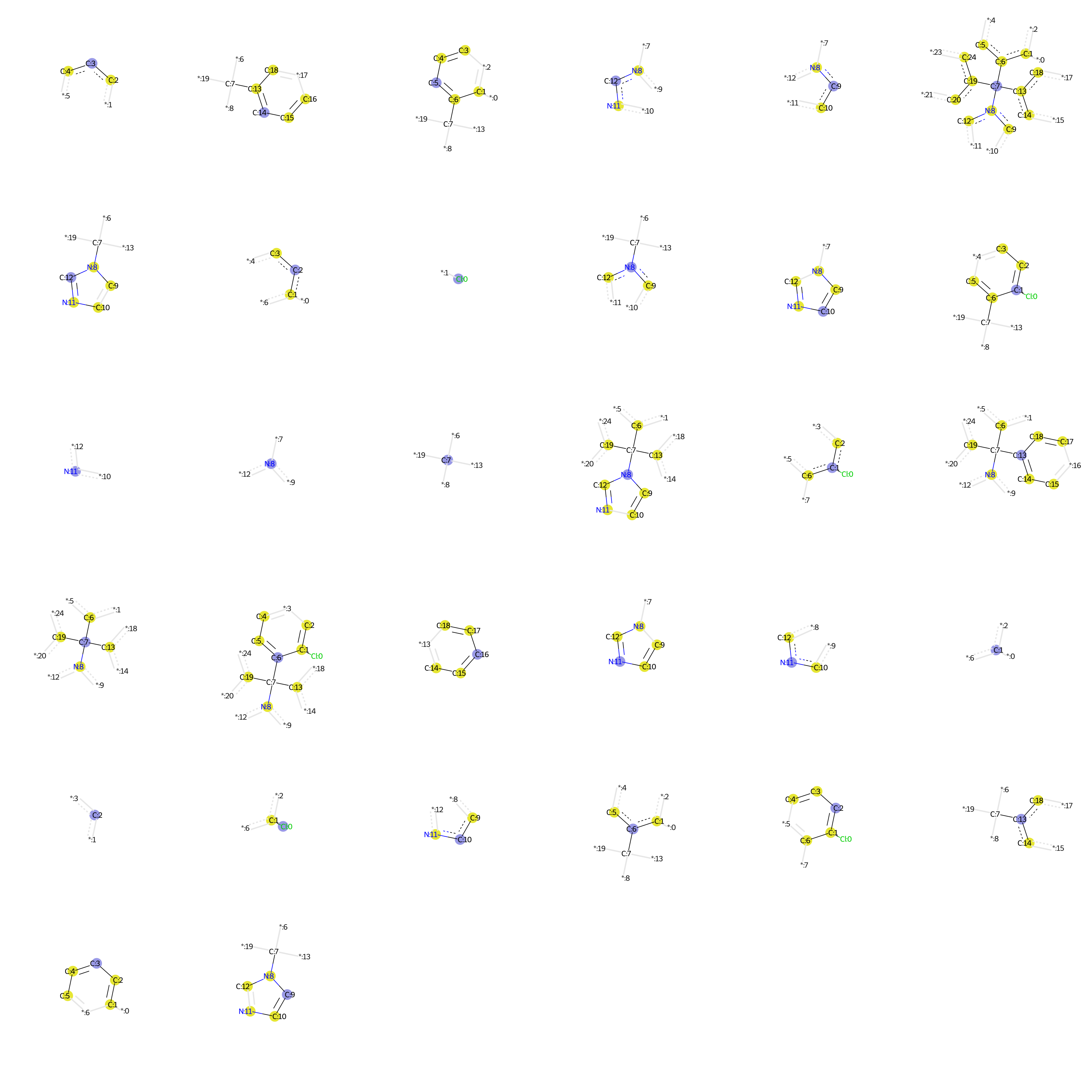

### Clozapine.bmp

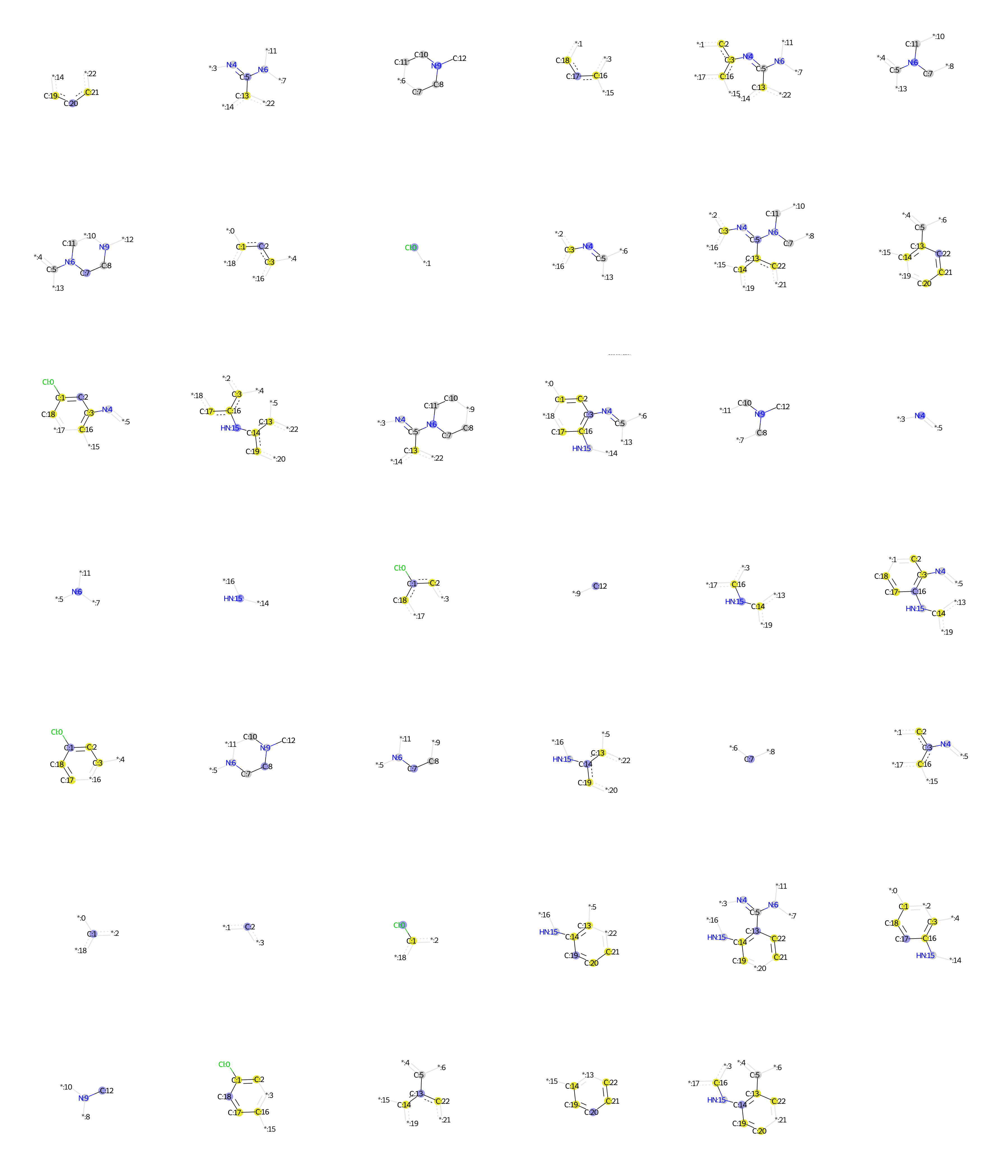

### ClozapineNoxide.bmp

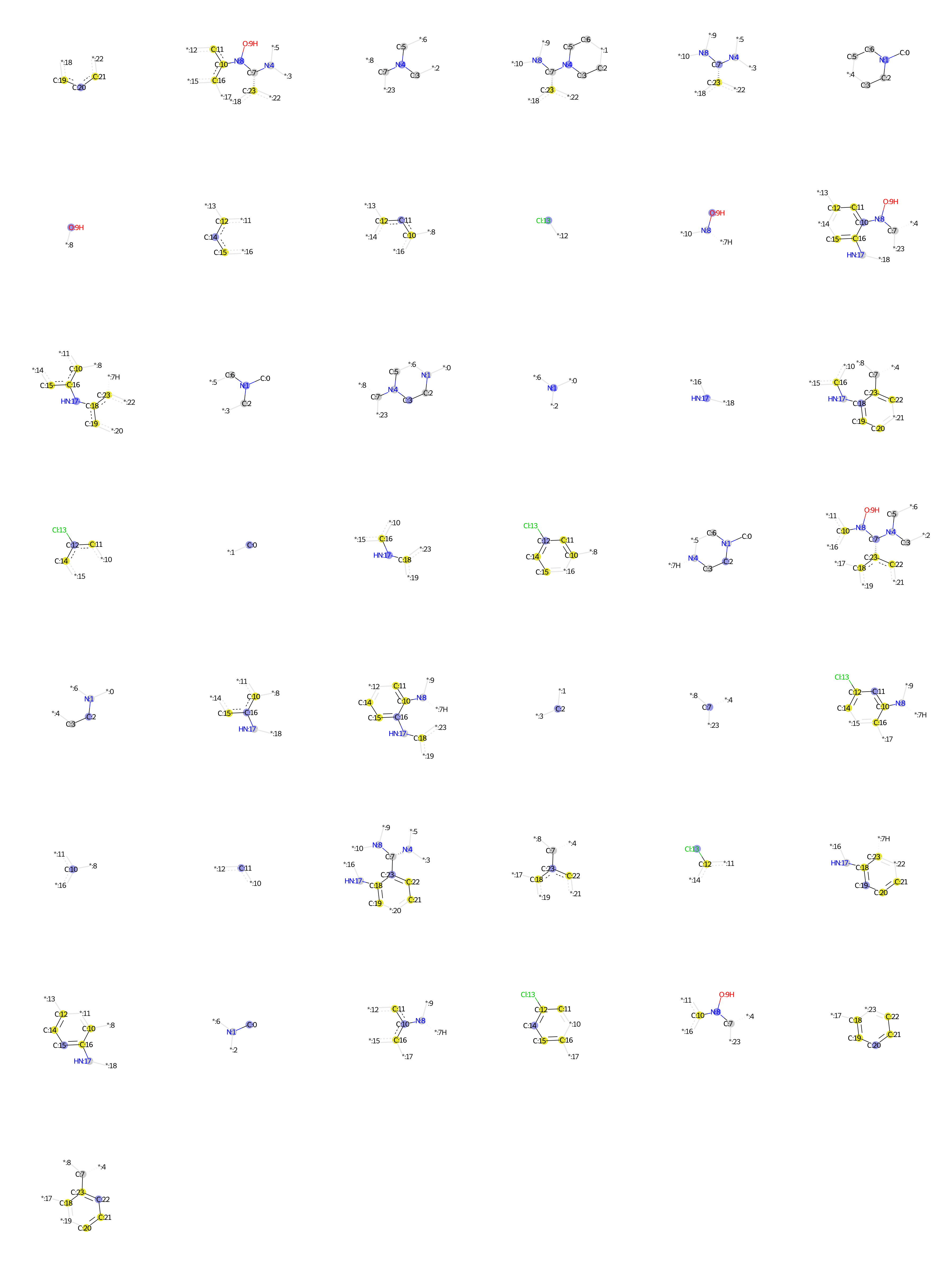

### Cocaethylene.bmp

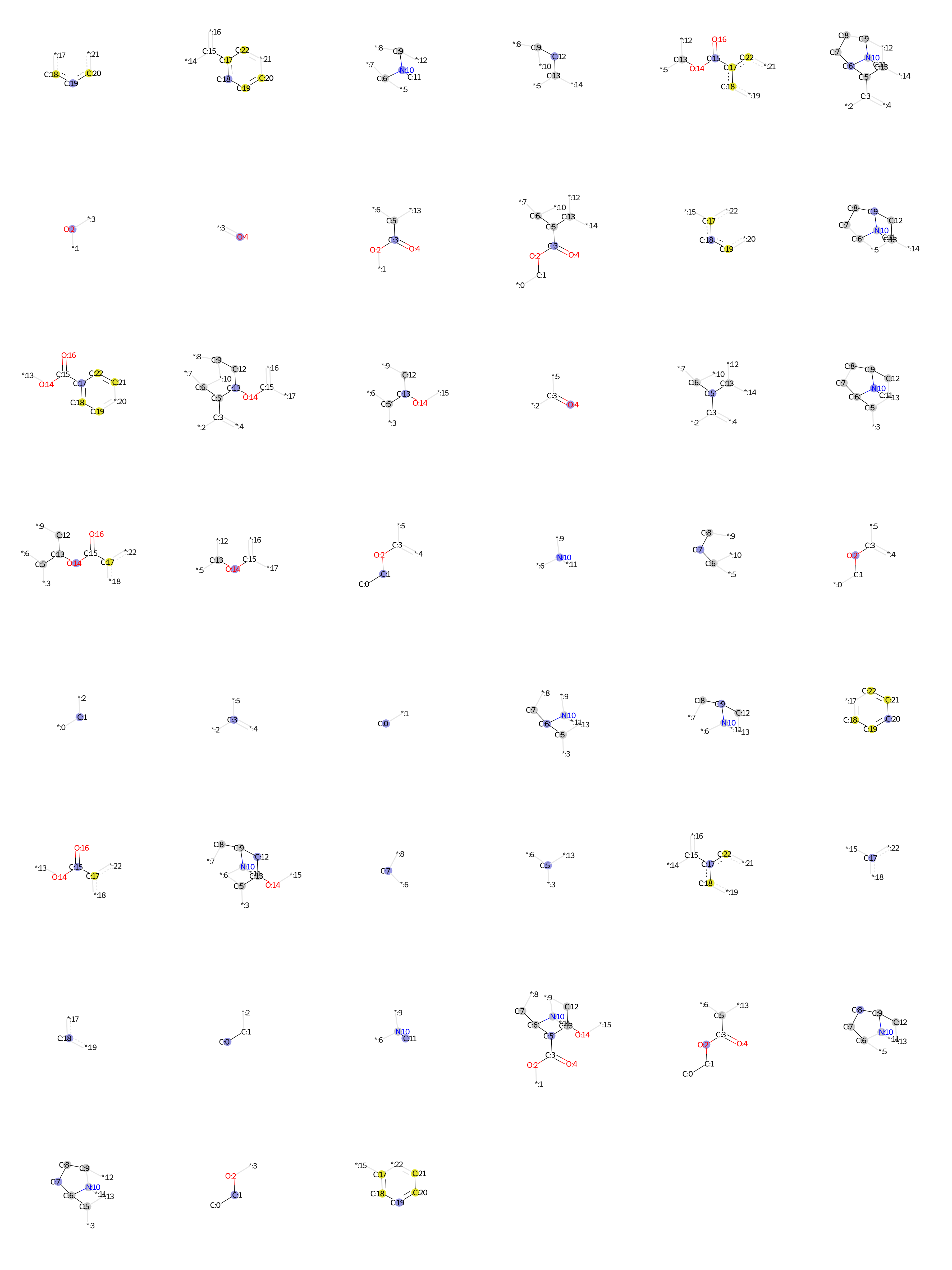

### Cocaine.bmp

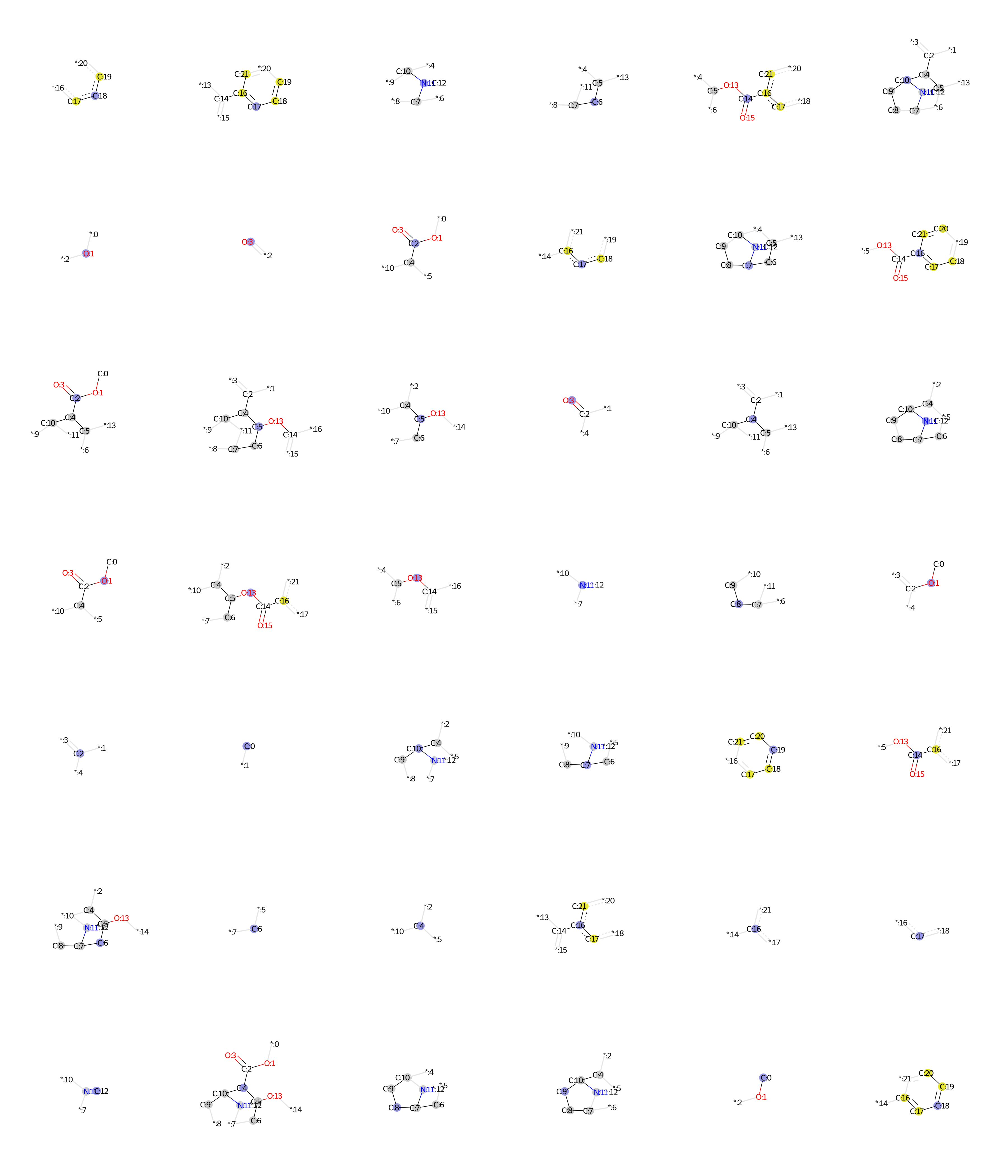

### Codeine.bmp

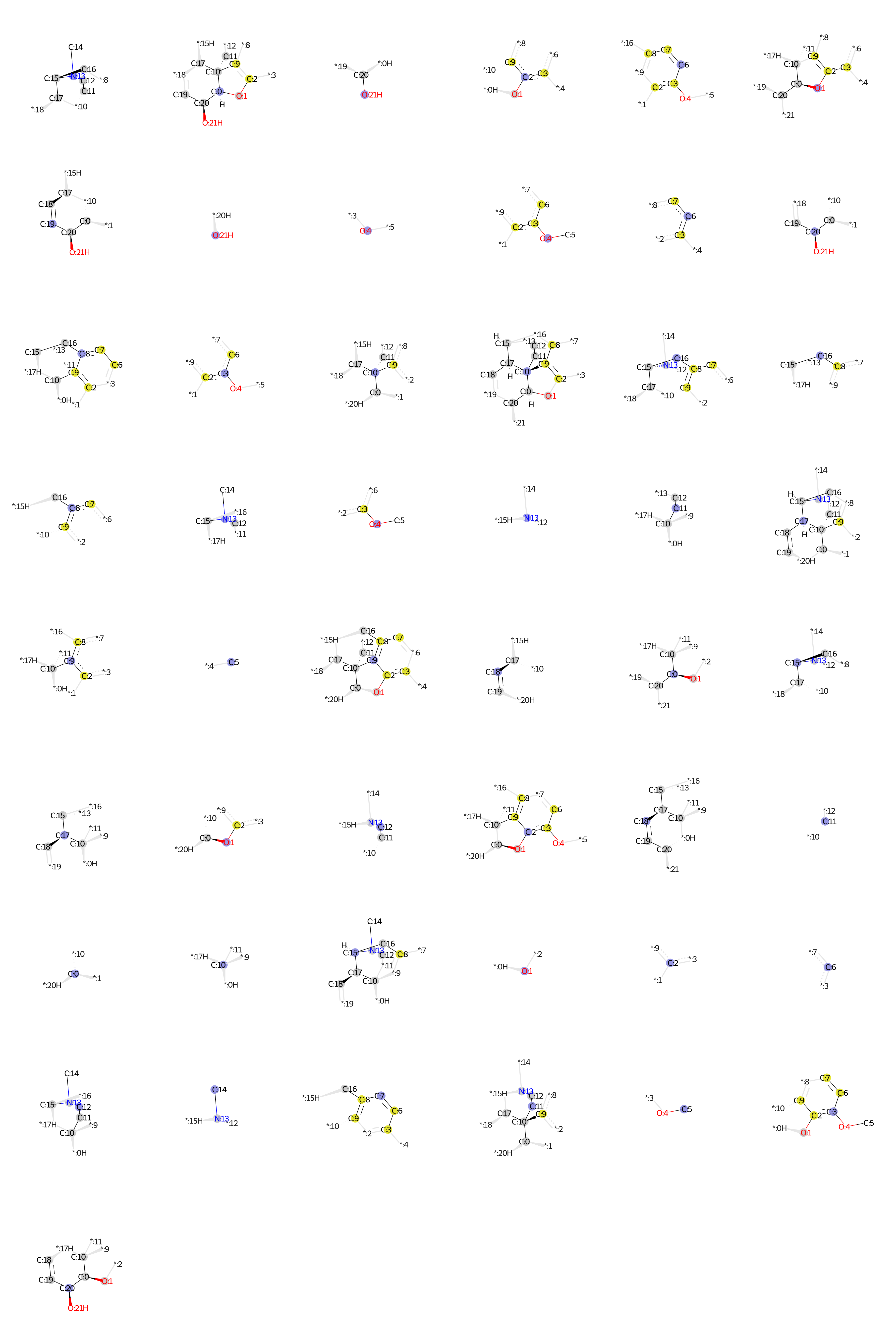

### d.bmp

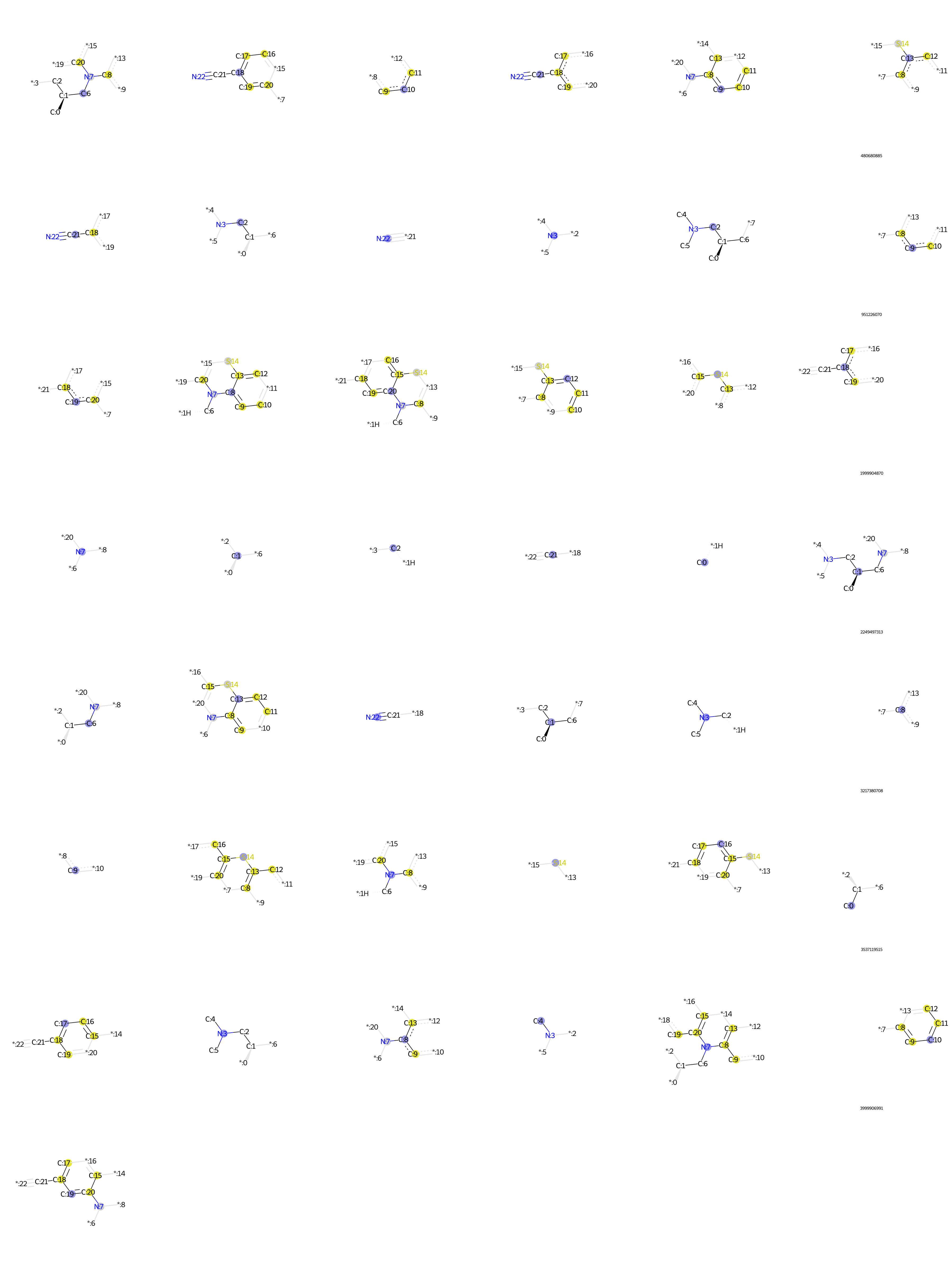

### Demethylolanzapine.bmp

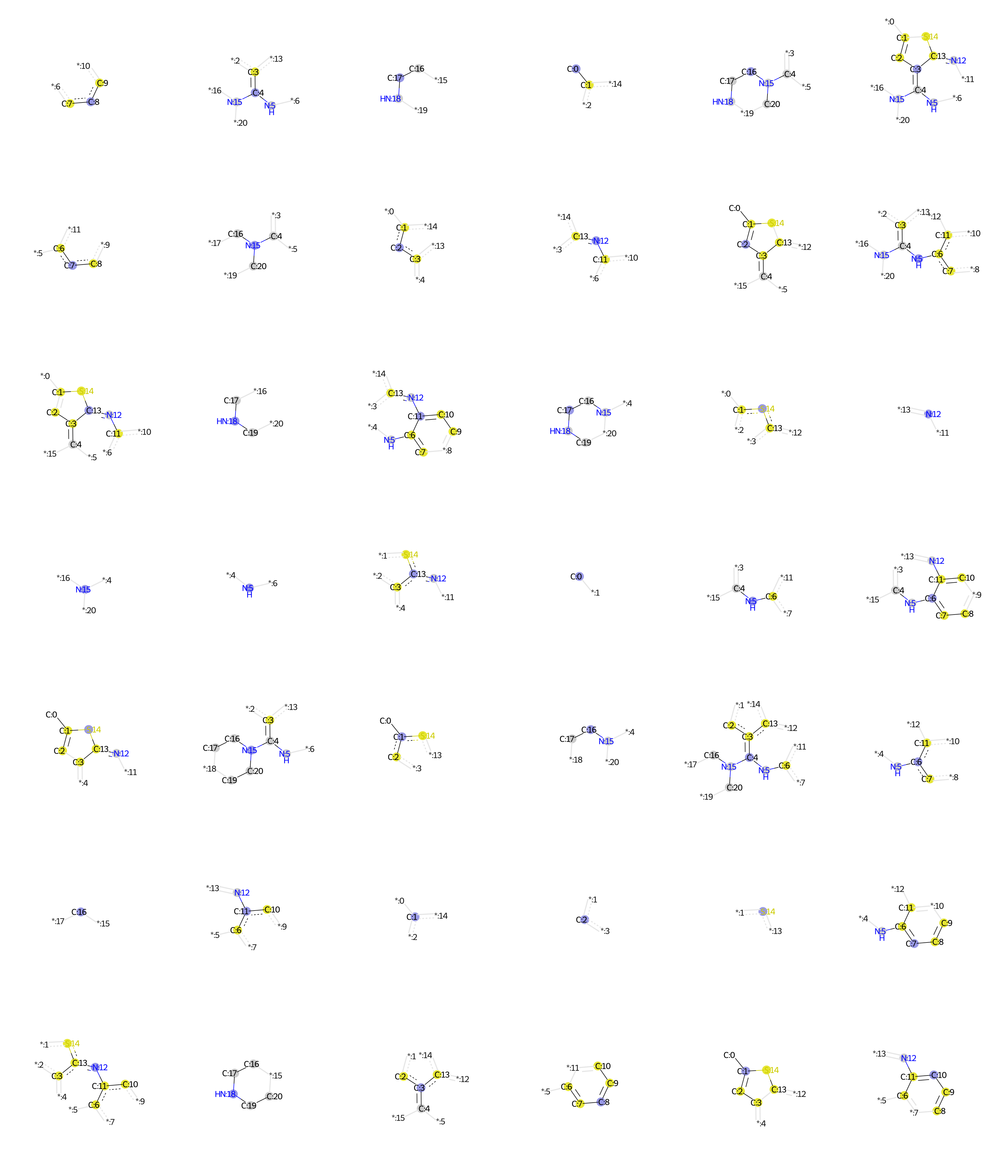

### Desbutyllumefantrine.bmp

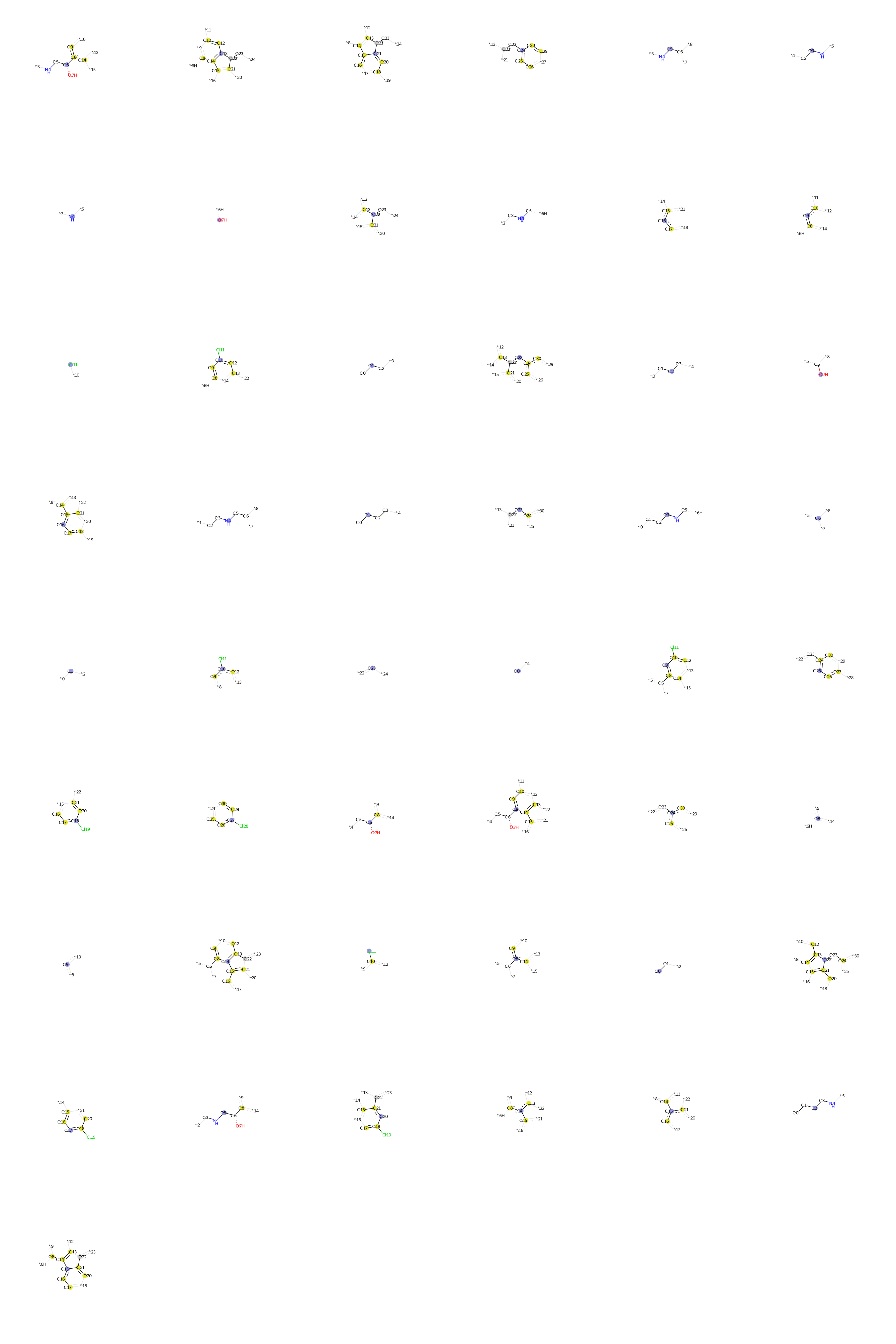

### Desipramine.bmp

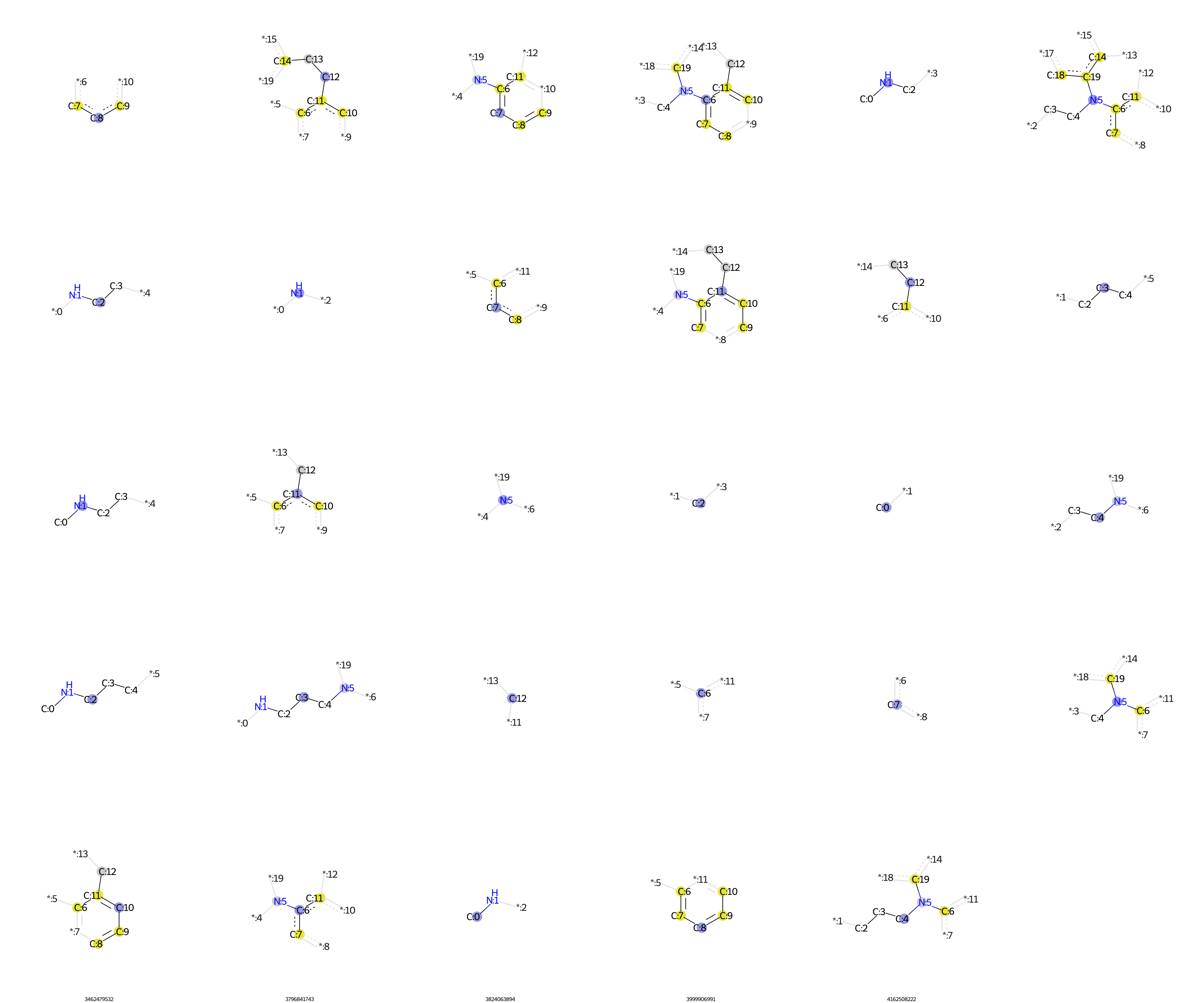
